# Ter-Seq: A high-throughput method to stabilize transient ternary complexes and measure associated kinetics

**DOI:** 10.1101/2022.05.18.492574

**Authors:** Gopinath Chattopadhyay, Shahbaz Ahmed, Nonavinakere Seetharam Srilatha, Apana Ashok, Raghavan Varadarajan

## Abstract

Regulation of biological processes by proteins often involves the formation of transient, multimeric complexes whose characterisation is mechanistically important but challenging. The bacterial toxin CcdB binds and poisons DNA Gyrase. The corresponding antitoxin CcdA extracts CcdB from its complex with Gyrase through formation of a transient ternary complex, thus rejuvenating Gyrase. We describe a high throughput methodology called Ter-Seq to stabilize probable ternary complexes and measure associated kinetics using the CcdA-CcdB-GyrA14 ternary complex as a model system. The method involves screening a YSD saturation mutagenesis library of one partner (CcdB) for mutants that show enhanced ternary complex formation. We also isolated CcdB mutants that were either resistant or sensitive to rejuvenation, and used SPR with purified proteins to validate the kinetics measured using surface display. Positions where CcdB mutations lead to slower rejuvenation rates are largely involved in CcdA-binding, though there were several notable exceptions. Mutations at these positions reduce the affinity towards CcdA, thereby slowing down the rejuvenation process. Mutations at GyrA14-interacting positions significantly enhanced rejuvenation rates, either due to reduced affinity or complete loss of CcdB binding to GyrA14. We examined the effect of different parameters (CcdA affinity, GyrA14 affinity, surface accessibilities, evolutionary conservation) on the rate of rejuvenation. Finally, we further validated the Ter-Seq results by monitoring kinetics of ternary complex formation for individual CcdB mutants in solution by FRET studies.

## Introduction

Regulation of several biological processes involves formation of short-lived transient multi-meric complexes. Characterization of such complexes is challenging, but mechanistically important. The binding of transcription factors to DNA, regulation of interferon signalling, and hypoxic response induced by transcription factors are examples of regulatory mechanisms which involve the formation of transient ternary complexes (Berlow et al., 2017; Kamar et al., 2017; Li et al., 2017). There have been few studies in which the molecular details of such short lived complexes have been characterised (Berlow et al., 2017; Kamar et al., 2017; Li et al., 2017; Schreiber, 2017).

The ccd toxin-antitoxin system is involved in plasmid maintenance and bacterial persistence (Bernard and Couturier, 1992; Gupta et al., 2017; Tripathi et al., 2012). The CcdB toxin binds and inhibits the function of homodimeric DNA Gyrase (Bernard and Couturier, 1992). DNA gyrase is an important, validated drug target (Collin et al., 2011). Both fluoroquinolone drugs and the bacterial toxin CcdB cause cell death through DNA gyrase poisoning (Aldred et al., 2014; Critchlow et al., 1997; Dao-Thi et al., 2005; Drlica and Zhao, 1997; Heddle and Maxwell, 2002; Heddle et al., 2000; Van Melderen, 2001; Shen et al., 1989). The intrinsically disordered C-terminal domain of the cognate antitoxin CcdA, facilitates the rejuvenation of the poisoned Gyrase-CcdB complex and thus neutralizes CcdB (De Jonge et al., 2009; Maki et al., 1996). Previously, we have used FRET studies to demonstrate that rejuvenation begins with the binding of CcdA to the CcdB-Gyrase complex to form a short-lived transient ternary complex, which subsequently collapses to release functionally restored Gyrase and CcdA-CcdB complex (Aghera et al., 2020). Rejuvenation is significantly enhanced by subsequent binding of a second CcdA molecule to the pre-formed CcdA-CcdB-Gyrase ternary complex.

In the present study, we have developed high throughput methodology to isolate probable stabilised ternary complexes using YSD coupled to Sanger sequencing. The CcdA:CcdB:Gyrase system was used as the first test system. Here we used the GyrA14 variant of Gyrase (Dao-Thi et al., 2005) and full-length CcdA as well as three variants of CcdA peptides (CcdA_37-72_, CcdA_45-72_ and CcdA_61-72_), which contain the necessary binding domains that mediate dissociation of the CcdB-GyrA14 complex (Aghera et al., 2020). The rejuvenation experiments on the yeast cell surface validated the earlier observations which demonstrated that the CcdA_37-72_ and CcdA_45-72_ peptides can rejuvenate the CcdB:GyrA14 complex to the same extent as full-length CcdA, whereas the CcdA_61-72_ peptide failed to rejuvenate. Further, a CcdB library containing ∼1600 point mutants was displayed on the yeast cell surface, and bound GyrA14 was rejuvenated with different concentrations of CcdA_45-72_ peptide for different time points. Sorting was carried out for two rounds to enrich for CcdB mutants having rejuvenation rate constants faster or slower than WT CcdB. Following sorting and plating, mutants identified from Sanger sequencing of individual colonies were revalidated by yeast surface display (YSD) experiments. CcdB mutants in regions involved in either CcdA or GyrA14 binding, as well as mutants at non-interacting sites were also purified, and their rejuvenation rate constants were determined using Surface Plasmon Resonance (SPR). From both YSD and SPR studies, it was observed that mutants in the CcdA interacting regions (hereafter referred to as CcdA interacting-site mutants) had slower rejuvenation rate constants than WT, whereas mutants in the Gyrase binding regions (hereafter referred to as Gyrase interacting-site mutants) had relatively faster rejuvenation rate constants. The affinities of a few of the CcdB mutants with CcdA_45-72_ peptide and GyrA14 were further determined using YSD and SPR respectively. It was observed that the rejuvenation rate constants of CcdA interacting site-mutants were weakly dependent of the strength of interaction with CcdA. However, for the Gyrase interacting-site mutants, rejuvenation rate constants were inversely correlated with the strength of interaction with Gyrase. It was also observed that the CcdB residues involved in GyrA14 interaction were more conserved than the CcdA interacting residues. Finally, we monitored the formation of probable ternary complexes for a few of the CcdB mutants identified above, using FRET (Fluorescence Resonance Energy Transfer). We observe stabilization of the ternary complex for most mutants at residues that are involved in interaction with CcdA. Conversely mutants at sites involved in Gyrase interaction decrease the lifetime of the ternary complex, relative to WT. Interestingly, we found two mutants at residues that are neither involved in interaction with CcdA nor Gyrase, to form a stabilised ternary complex relative to WT. In summary, we experimentally describe and validate methodology that can be used to stabilise and characterize transient, macromolecular complexes in a high throughput fashion.

## Results

### CcdB rejuvenation is independent of the length of the CcdA peptide

To determine the effect of different lengths of CcdA on CcdB rejuvenation from the CcdB:GyrA14 complex, the rejuvenation of WT CcdB was carried out with different concentrations of full-length CcdA, CcdA_37-72_, CcdA_45-72_ and CcdA_61-72_ peptides on the yeast cell surface. Consistent with a previous study (Aghera et al., 2020), it was observed that the CcdA_37-72_ and CcdA_45-72_ peptides rejuvenated WT CcdB with rates similar to that of full-length CcdA, whereas CcdA_61-72_ failed to rejuvenate WT CcdB, consistent with earlier SPR and FRET studies (Figure S1).

### TER-Seq : A high throughput methodology to identify stable ternary complexes

In an attempt to identify mutants which can form stabilised CcdA:CcdB:GyrA14 ternary complexes, a CcdB library containing ∼1600 single-site mutants was displayed on the yeast cell surface (Figure 1). The library was first incubated with the cognate partner GyrA14 for an hour. The excess unbound GyrA14 was removed and the library was then rejuvenated from the bound Gyrase by addition of the second cognate partner, CcdA_45-72_ (Figure 1). The reaction was quenched after 5 s and 30 s by adding a 10 fold excess of purified WT CcdB to identify mutants having faster and slower rejuvenation rate constants than the WT respectively (Figure 1). The population showing binding to both GyrA14 and CcdA_45-72_ was sorted for two rounds and the different sorted populations were regrown, the isolated plasmids were transformed in *Top10Gyrase*, and the colonies sent for Sanger sequencing for identification of mutants which have a slower rejuvenation rate than WT CcdB (Figure 2A). Additionally, the mutants with faster rejuvenation rate constants, displaying a higher bound CcdA signal than WT were subjected to two rounds of sorting. The final sorted mutants were subjected to Sanger sequencing in a similar fashion as described above (Figure 2B).

**Figure 1.**
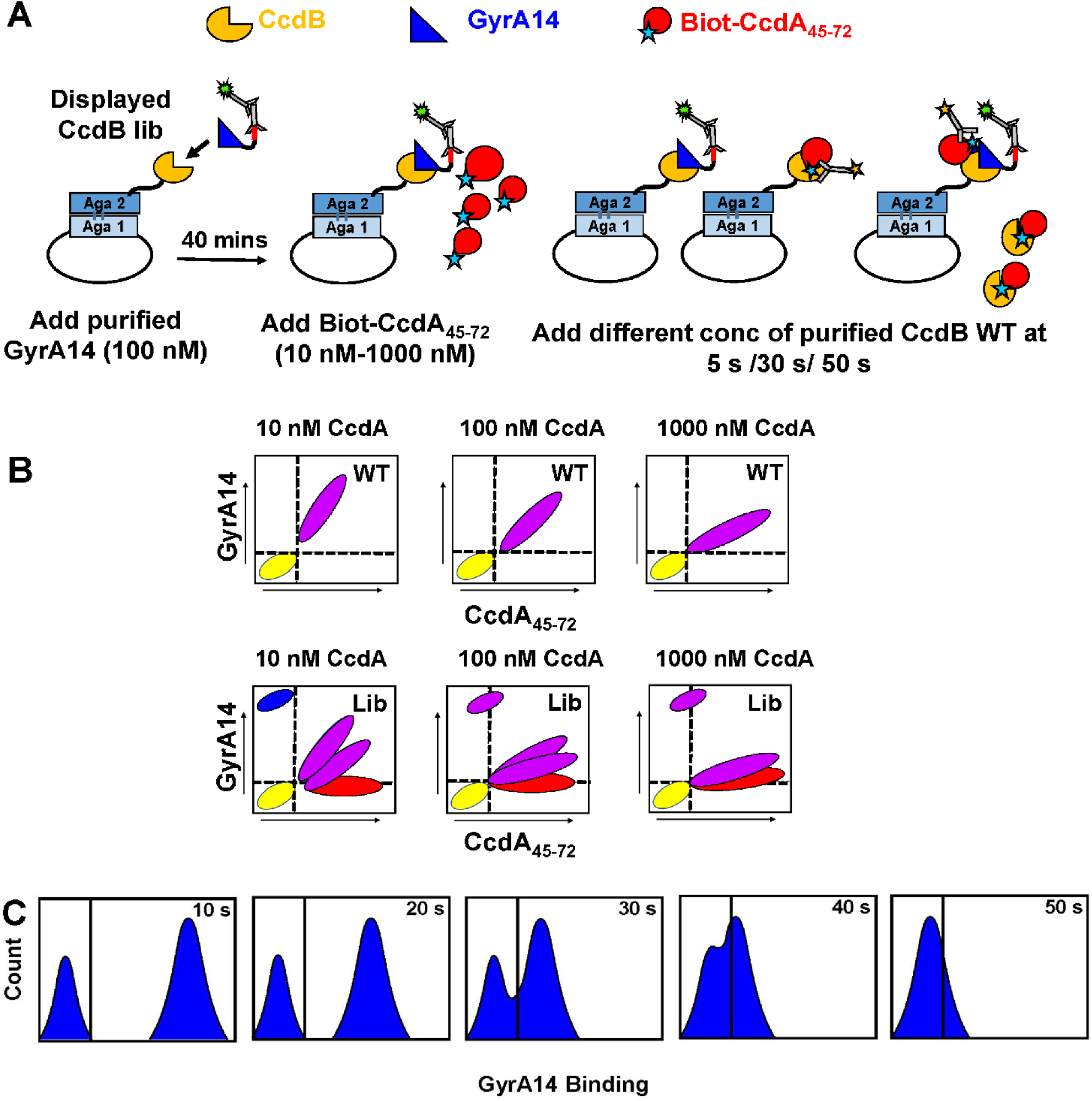
Schematic representation of the methodology to isolate a stabilised ternary complex. (A) The CcdB library is displayed as a fusion protein on the surface of yeast cells and binding to its cognate partner GyrA14 is observed by flow cytometry. Introduction of a second binding partner CcdA_45-72_ may generate different populations as a function of time, depending on the relative affinities of the mutants towards GyrA14 and CcdA_45-72_. The reaction is quenched by the addition of a molar excess of the purified WT CcdB at different time points to prevent further rejuvenation of GyrA14 from its complex with YSD CcdB. (B) (Top) The CcdA_45-72_ and GyrA14 binding of WT is shown as a function of different CcdA_45-72_ peptide concentrations. The reaction was quenched after 30 secs. (Bottom) The CcdA_45-72_ and GyrA14 binding of the CcdB library is shown as a function of different CcdA_45-72_ peptide concentrations. The reaction was quenched after 30 secs. Mutations at the Gyrase-interacting sites will likely cause faster rejuvenation upon addition of CcdA_45-72_ (red populations). However, mutations at the CcdA-interacting interface will likely result in different populations. Mutants at the hot-spot residues will lose CcdA binding and thereby fail to rejuvenate (blue populations) whereas mutants at other interfacial residues may show reduced binding resulting in slower rejuvenation (purple populations). Adding higher concentrations of CcdA_45-72_ or performing the reaction for longer time periods may permit the rejuvenation of CcdB mutants with mutations at CcdA binding site residues, though at a significantly slower rate than that observed for WT CcdB. Populations that show significant amounts of both bound CcdA and GyrA14 may be enriched in ternary CcdA_45-72_:CcdB:GyrA14 complexes. (C) Schematic of the rejuvenation rate calculation using Yeast Surface Display (YSD). The CcdB mutants were displayed as a fusion protein on the yeast cell surface and incubated with the cognate partner GyrA14 for 40 minutes, followed by addition of a second binding partner CcdA_45-72_. The reaction was then quenched by the addition of ten fold molar excess of the purified WT CcdB over CcdA_45-72_ at five different time points, and cells subjected to FACS. The extent of GyrA14 binding on the yeast cell surface was estimated by incubating the induced CcdB mutants with 100 nM FLAG tagged GyrA14, followed by incubation with mouse anti-FLAG antibodies (1˸300 dilution) and rabbit anti-mouse antibodies conjugated to Alexa Fluor 633 (1:1600 dilution). *See also Figure S1*.

**Figure 2.**
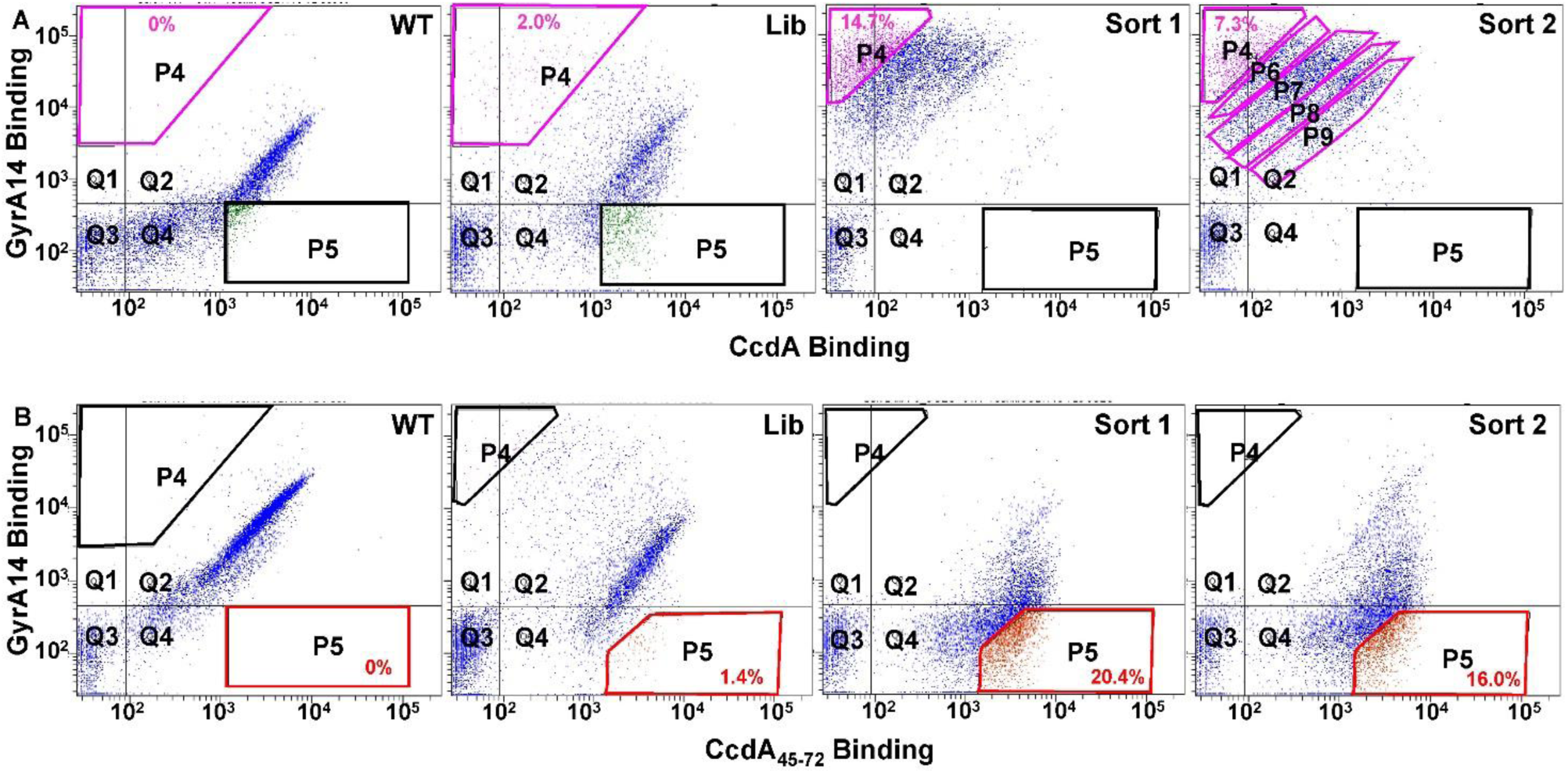
Enrichment of CcdB mutants with slower and faster rejuvenation rate constants (relative to the WT) following multiple rounds of FACS. (A) WT CcdB, CcdB library, Sort-1 CcdB library (sorted in gate P4) and Sort-2 CcdB library (sorted in gate P4, P6, P7, P8 and P9) rejuvenated with 100 nM CcdA_45-72_ for 30 s from left to right. (B) WT CcdB, CcdB library, Sort-1 CcdB library (sorted in gate P5) and Sort-2 CcdB library (sorted in gate P5) rejuvenated with 10 nM CcdA_45-72_ for 5 s from left to right. 100 nM of GyrA14-3xFLAG was used in all the cases for analysis. 100 nM of CcdA_45-72_ for 30 s (A) and 10 nM of CcdA_45-72_ for 5 s (B) were used for sorting mutants with slower and faster rejuvenation rate constants than the WT CcdB respectively. The populations gated in purple and blue (A) shows enrichment of mutants with slower rejuvenation rate than WT CcdB, while those in red (B) show enrichment of mutants with faster rejuvenation rate than the WT. *See also Figure S2-S6*.

### Mutants at CcdA binding sites have a slower rejuvenation rate than WT CcdB whereas mutations at Gyrase binding sites rejuvenate faster than the WT

Mutants enriched after two rounds of sorting, that displayed rejuvenation rate constants slower than the WT were residues largely involved in CcdA-binding (Figure S2). Mutations at these positions reduced the affinity towards CcdA, thereby slowing down the rejuvenation process (Figure S3). However, the expression and GyrA14-binding activities of these mutants were similar to WT (Figure S3). For example, mutants showing the slowest rejuvenation such as Y8D, Y8E, S12W, P35W, A69R and A69W have either lost their binding or bind very weakly to CcdA (Figure S2). Other mutants such as L42K, L42W, V46R, V46Y, M64E, M64R and T66E have significant reductions in their affinities to CcdA and therefore, rejuvenate slower than the WT (Figure S2, Figure S3).

Mutations at Gyrase-interacting positions significantly enhanced the rejuvenation rate constants either because of reduced affinities or complete loss of binding towards Gyrase (Figure S4). However, these mutants showed similar expression levels and CcdA-binding abilities, relative to that of WT CcdB (Figure S5). Mutants such as F17I, S22G, I24E, I24G, I24R, I24T, I25G, N95L, N95R, W99C and G100A have lost their binding to Gyrase and therefore, rejuvenate faster than the WT (Figure S4, Figure S5).

For a few selected CcdB mutants, with reduced affinity to CcdA_45-72_, rejuvenation was carried out for longer time periods (Figure S6). Most mutants retained significant GyrA14 binding even after 200 s (Figure S6).

### The rejuvenation rate constants of CcdB mutants are weakly dependent of the strength of their interactions with CcdA

The rejuvenation rate constants of individual CcdB mutants were determined on the yeast cell surface (Figure 3). GyrA14 bound to CcdB mutants on the yeast cell surface, was rejuvenated by addition of a second binding partner, CcdA_45-72_. This was followed by quenching the rejuvenation reaction by addition of a ten-fold molar excess of purified WT CcdB protein at five different time points (Figure 3). For CcdA interacting-site CcdB mutants, rejuvenation was carried out with 100 nM CcdA_45-72_ (Figure 3A-B), except for R10G, for which it was carried out with 10 nM CcdA_45-72_ (Figure 3C). For GyrA14 interacting-site CcdB mutants, rejuvenation was carried out with 10 nM of CcdA_45-72_ (Figure 3D, Table S2). CcdB mutants at residues which are not involved in interaction with either CcdA or GyrA14 were efficiently rejuvenated with both 100 (Figure 3E) and 10 nM of CcdA_45-72_ (Figure 3F, Table S2).

**Figure 3.**
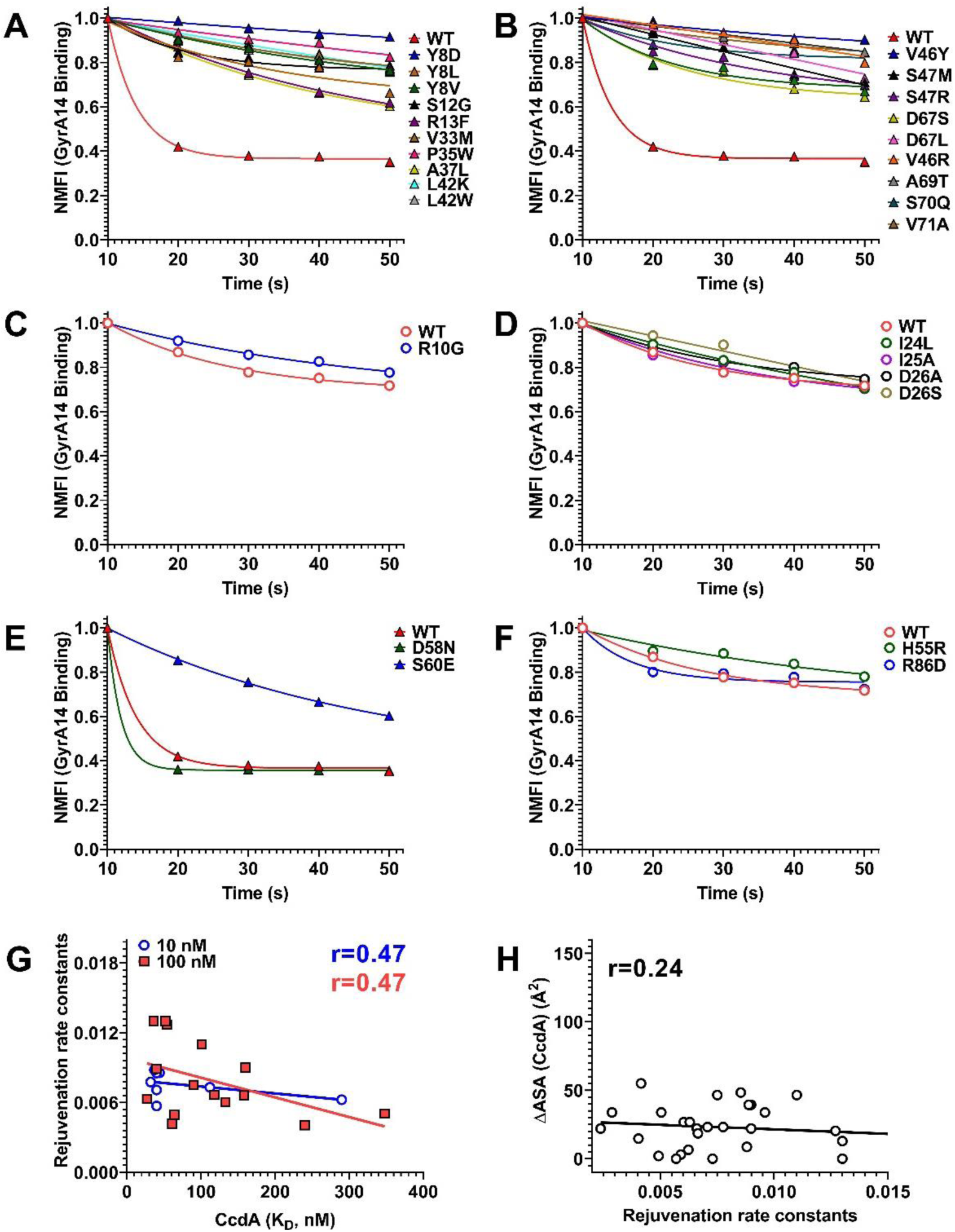
Rejuvenation rate constant is moderately correlated with the strength of interaction with CcdA. (A-C) CcdB mutants involved in CcdA interaction with slower rejuvenation rate constants than WT were rejuvenated with (A-B) 100 and (C) 10 nM CcdA_45-72_ peptide. (D) CcdB mutants involved in GyrA14 interaction with faster rejuvenation rate constants than the WT were rejuvenated only with 10 nM CcdA_45-72_ peptide. (E-F) CcdB mutants which do not interact with either CcdA or GyrA14 and with largely similar rejuvenation rate constants as the WT were rejuvenated with (E) 100 and (F) 10 nM CcdA_45-72_ peptide. Though not involved in interaction with CcdA or GyrA14, S60E had a slower rejuvenation rate constant than WT. The rate of rejuvenation was calculated from the Mean Fluorescence Intensity (MFI) values of Gyrase-binding obtained after 10, 20, 30, 40 and 50 s of rejuvenation. Further, the MFI values at each time point were normalised with the MFI value obtained after 10 s of rejuvenation (also listed in Table S2). (G) The rate constants of rejuvenation calculated with 100 nM CcdA_45-72_ (red) and with 10 nM CcdA_45-72_ (blue) are weakly dependent on the affinity of the mutants towards CcdA_45-72_. (H) The rejuvenation rate constants are independent of their ΔASA (CcdA) values, where ΔASA (CcdA) is the difference between solvent accessible surface area of a particular CcdB residue in the (fictitious) free form (removing CcdA from the CcdB-CcdA complex) and CcdA-bound form (PDB ID: 3G7Z). *See also Figure S7*.

To determine if rejuvenation rate constants of CcdB mutants were dependent on the strength of interaction with their binding partner, CcdA, the affinities for some of the CcdB mutants with CcdA_45-72_ peptide were also measured by YSD (Figure S7, Table S2). In all cases, the CcdB mutants were displayed on the yeast cell surface and titrated serially with CcdA_45-72_ peptide. The binding was measured on a BD Accuri flow cytometer. For several CcdA interacting-site CcdB mutants, the affinities towards CcdA were two to thirty fold lower than that of WT CcdB (Figure S7A-C, Table S2), except for R13F, L42W, V46L and S47R (similar affinity relative to the WT). Further, some CcdA interacting site, CcdB mutants such as Y8D, S12W, P35W and A69R had completely lost their binding towards CcdA (Figure S7D, Table S2). CcdA-binding affinities of CcdB mutants in the GyrA14 interaction sites were similar to that of WT CcdB even though there is partial overlap of both binding sites (Figure S7E, Table S2). For CcdB mutants, at residues which are not involved in CcdA or GyrA14 binding, such as H55R, D58N and S60E, affinities towards CcdA remained unchanged except for R86D (Figure S7F, Table S2).

Regression analysis between the rejuvenation rate constants and the CcdA-binding affinities of the CcdB mutants showed a weak correlation of 0.47 between the two parameters (Figure 3G). Further, there was no correlation between the ΔASA (difference between the solvent accessible surface areas of a CcdB residue in its free form (3VUB) and CcdA-bound form (3G7Z)) calculated using NACCESS, and the rejuvenation rate constants of the CcdB mutants (Figure 3H). Overall, rejuvenation rate constants of the CcdB mutants are moderately correlated with the strength of interaction with CcdA and there are other factors which influence the process of CcdA-mediated rejuvenation of the CcdB:GyrA14 complex.

### Mutational effects on rejuvenation rate constants measured by SPR are consistent with those from YSD

To further validate results from YSD, a few CcdB mutants at either CcdA or GyrA14 interacting positions, as well as a few CcdB mutants that do not interact with either CcdA or GyrA14 were purified and their rejuvenation rate constants were determined by SPR (Figure S8, Table S3-S5). In all cases, rejuvenation was carried out with four different concentrations (10, 25, 100 and 200 nM) of CcdA_45-72_ peptide (Figure 4A-D). The rejuvenation rate constants of the CcdB mutants determined at each CcdA_45-72_ peptide concentration were further normalised with respect to the values for WT CcdB at the same CcdA_45-72_ peptide concentration as described in the SI methods and these values are referred to as Fold change. For CcdA interacting-site CcdB mutants, rejuvenation rate constants with CcdA were approximately two to thirty-fold lower than for WT CcdB except for R10K, P28N and P28Y, (Figure 4A-B, Figure S9, Table S3). The most significant effect of mutations on rejuvenation rate constants was observed for residues Y8, S12, Y14 in the 8-14 loop region, residues A37, L42, S43, V46, S47 and L50 in loop 37-50 and residues D67, A69, S70 and V71 in loop 66-72 (Figure 4A, S9, Table S3). For GyrA14 interacting-site CcdB mutants, the rejuvenation rate constants were higher that of the WT CcdB, except at higher CcdA_45-72_ concentrations (≥100 nM), where everything rejuvenates to the same extent as the WT (Figure 4A-B, Figure S10, Table S4). The rejuvenation rate constants for CcdB mutants at residues which do not interact with either CcdA or GyrA14, were either similar to or faster than for WT CcdB (Figure 4C-D, S11, Table S5) with the exception of the following mutants discussed below. Interestingly, mutants like A39R, V53P, V54G, G57C, E59P, E59Q, S60E, E79I and R86E at non active-site residues showed twenty and four fold lower rejuvenation rate constants respectively than the WT, even though their dissociation rates in the absence of CcdA were similar to that of WT CcdB (Figure 4C-D, Figure S11, Table S5). Some of the mutants at non active-site residues like K9R, D44V, D58N, S60G, S84G and R86D on the other hand have much faster rejuvenation rate constants as compared to the WT. Mutations were mapped onto the CcdB structure to emphasize the above points (Figure 4E). The likely structural basis for these anomalous effects is discussed in a later section.

**Figure 4.**
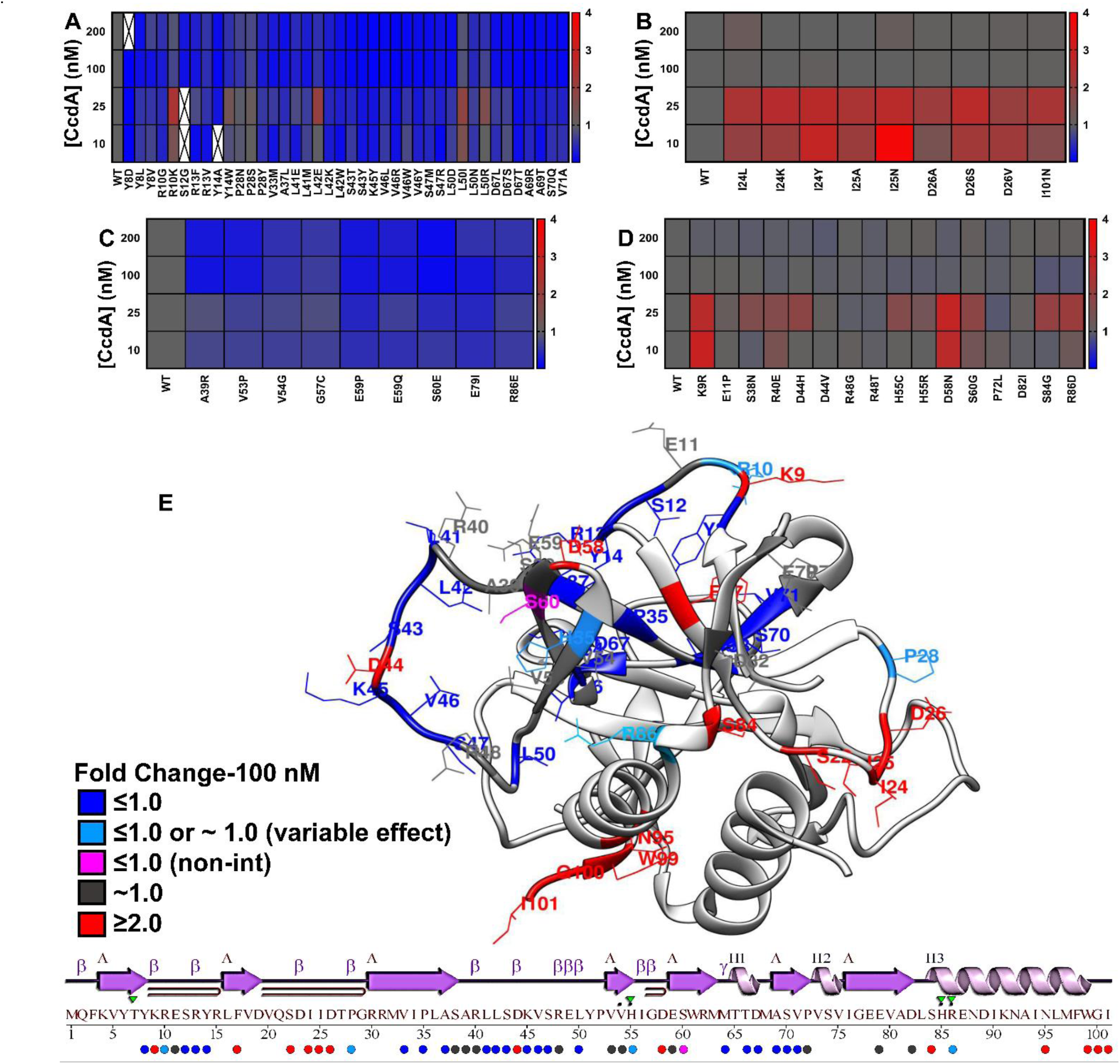
Analysis of rejuvenation rate constants of different CcdB mutants by SPR and YSD. (A-D) Heatmaps are coloured according to the magnitude of the nomalized dissociation rate constants of the CcdB mutants mediated by various concentrations of CcdA_45-72_. Blue, grey and red colour represent mutants having rejuvenation rate constants slower, similar and faster than WT respectively. All the rejuvenation rate constants presented in the heatmaps are normalised with respect to the WT as described in SI Methods as fold change. (A) Fold change of rejuvenation rate constants mediated by CcdA_45-72_ in CcdA interacting regions, where most mutations affect the rejuvenation process. The rate constants are approximately 5-30 fold lower for the CcdA-interacting-site mutant CcdB:GyrA14 complexes than that seen for the WT CcdB:GyrA14 complex. The white cells with a cross in the heatmap indicates that data is not available for that condition. (B) Fold change of rejuvenation rate constants mediated by CcdA_45-72_ for CcdB mutants at residues involved in interaction with GyrA14. All these mutations accelerate the rejuvenation process. At higher concentration of CcdA_45-72_≥100nM, the rejuvenation rate constants, are similar to WT. (C) Fold change of rejuvenation rate constants mediated by CcdA_45-72_ in regions not involved in interaction with either CcdA or GyrA14, but where mutations slow down the rejuvenation process. (D) Fold change of rejuvenation rate constants mediated by CcdA_45-72_ in regions not involved in interaction with either CcdA or GyrA14 but where mutations have either a WT like effect or rejuvenate faster than WT. The faster rejuvenation rate constants of such mutants are largely due to their altered binding kinetics to GyrA14. (E) Positions where mutations alter rejuvenation rate constants as determined from both YSD and SPR studies are mapped on the crystal structure of CcdB (PDB ID:3VUB). For clarity, residues of only one chain is color coded. Positions where most mutations slow down the rejuvenation rate constant are shown in blue and are involved in CcdA interaction. Four residues R10, P28 (involved in CcdA binding) and H55, R86 (not involved in interaction with either Gyrase or CcdA), where mutations have variable effects are shown in sky blue. The residues where most mutations have enhanced the rejuvenation rate constant are shown in red and are involved in Gyrase interaction, except for a few residues such as K9, D44, D58, S60, S84 and R86 which are not involved in interaction with either Gyrase or CcdA. Rejuvenation experiments for mutants at positions F17, S22, N95, W99 and G100 were determined from YSD only, and owing to their poor GyrA14 binding, rejuvenation rate constants were not determined for mutants at these positions. Mutations at positions that are not involved in either CcdA or Gyrase binding but rejuvenate slower than WT are shown in magenta. All the positions where mutations have WT like rejuvenation rate constants are shown in dark grey. The colour keys in square boxes show the fold change in rejuvenation rate constants for each category at 100 nM CcdA_45-72_. Shown below the crystal structure is the one-dimensional representation of the secondary structure of CcdB aligned to its amino acid sequence, with spheres below the sequence highlighting the positions where mutations were made in this study. The colour of the spheres represents the mutational effect as described in the above crystal structure. *See also Figure S8-S14*.

Mutations were classified into aliphatic, aromatic, charge and polar categories to further delineate the effect of different types of substitutions on rejuvenation rate constants (Table S6). The rejuvenation rate constants for the CcdB mutants were normalized as discussed above (Table S6). For all four concentrations of CcdA_45-72_ peptide used, it was observed that aromatic substitutions have the highest effect on the rejuvenation rate constants, followed by charged and aliphatic substitutions. Substitutions to polar amino acid had little effect on the rejuvenation rate constants (Table S6). Overall, the rate constants measured by YSD and SPR are in good agreement (Table S2).

### The strength of interaction with Gyrase strongly influences the rate constant of rejuvenation

The affinities of all CcdB mutants with GyrA14 were also measured by SPR as described previously (Chattopadhyay et al., 2021) (Figure S11-S14, Table S7-S9). The rejuvenation rate constants measured by SPR correlated well with rejuvenation rate constants determined from YSD. There was no correlation between the ΔASA (GyrA14) and the rejuvenation rate constants for the GyrA14 interacting CcdB mutants (Figure 5A). However, the normalised rejuvenation rate constant of CcdB mutants at GyrA14 interacting residues, and affinities of the mutants towards GyrA14 were well correlated at 10 and 25 nM CcdA_45-72_ concentration (Figure 5B). At higher concentrations of CcdA the correlation becomes moderate. There was no such correlation observed for CcdB mutants at residues not involved in binding with GyrA14. An exception to this was mutations at P28. Although not involved in GyrA14 binding, mutations at this residue resulted in a 10-20 fold difference in affinity to GyrA14 (Table S7).This might arise from the unique conformational restrictions associated with Proline (Bajaj et al., 2007) .

**Figure 5.**
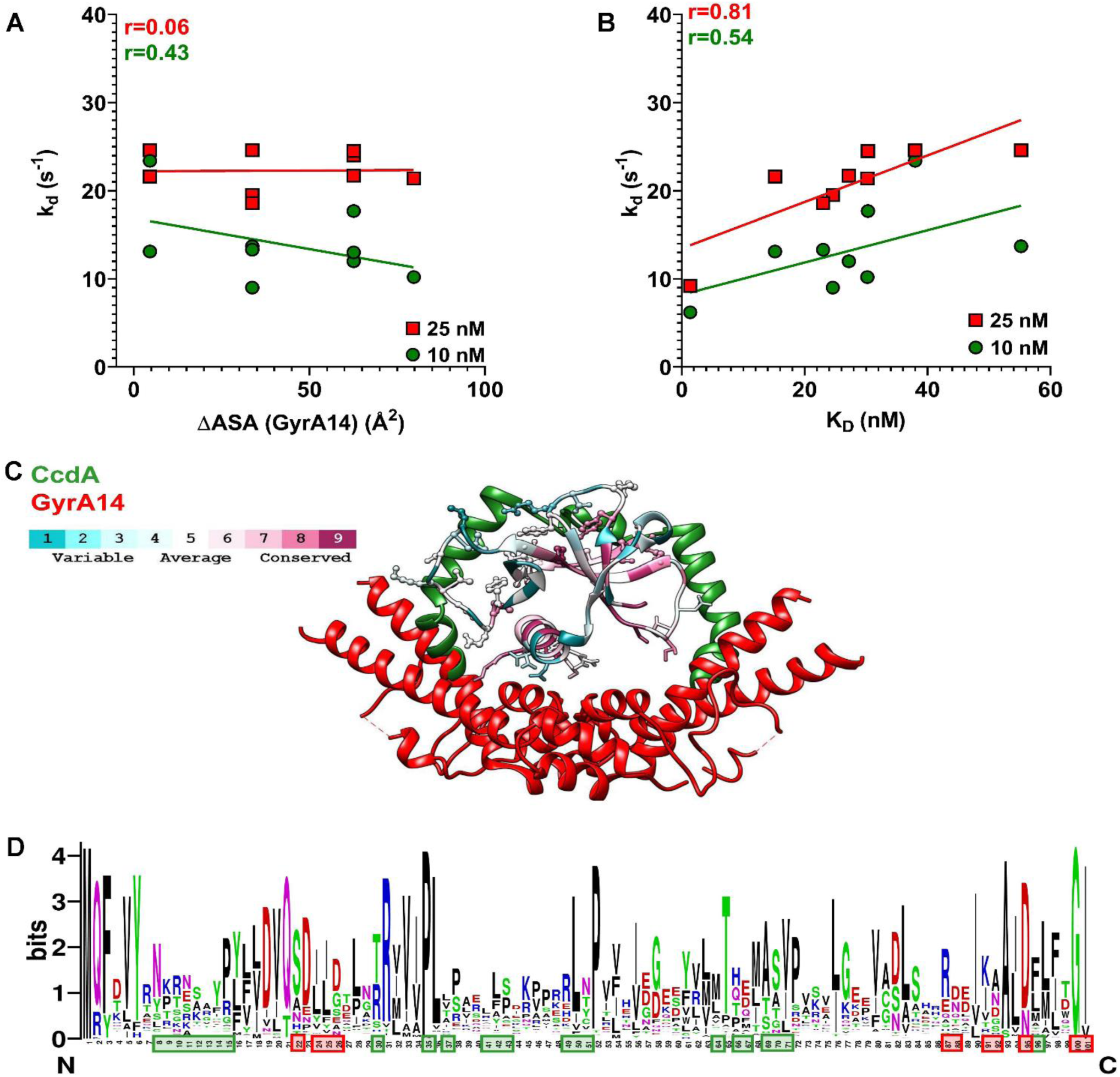
Correlation of rejuvenation rate constants of mutants at Gyrase interacting residues with various parameters. (A) The rate constants of rejuvenation calculated by SPR with 10 CcdA_45-72_ (green) and 25 nM CcdA_45-72_ CcdA_45-72_ (red) are independent of the ΔASA (GyrA14) values for the corresponding positions, where ΔASA (GyrA14) is the difference between solvent accessible surface areas of a CcdB residue in the (fictitious) free form (removing GyrA14 from CcdB:GyrA14 complex) and GyrA14-bound form (PDB ID: 1X75). (B) The correlation between rate constants of rejuvenation of the Gyrase interacting CcdB mutants calculated from SPR with the affinity of interaction of the CcdB mutants with GyrA14, using 10 nM CcdA_45-72_ (green) and 25 nM CcdA_45-72_ (red). (C) The conservation scores of residues calculated by Consurf are mapped on the CcdB crystal structure (PDB ID: 3VUB). Superposition of the crystal structures of the CcdA:CcdB complex (PDB ID: 3G7Z) with the CcdB:GyrA14 (PDB ID: 1X75) complex highlighting the relative binding positions of the CcdA (green) and GyrA14 (red) on CcdB. The structure was generated by aligning the CcdB homodimers in both the structures. Only a single chain of CcdB is shown for clarity. The residues involved in GyrA14 interaction are highly conserved as compared to the CcdA interacting residues. (D) Web logo showing residue conservation for the CcdB protein. Residues that are involved in GyrA14-interaction (red boxes) are more conserved as compared to CcdA-interacting residues (green boxes). *See also Figure S9-S14*.

### The CcdB residues involved in GyrA14 interaction are evolutionarily conserved

The evolutionary conservation scores (at each position) were also calculated for the CcdB protein (Figure 5C). The overall conservation across all the positions in the CcdB protein revealed that residues involved in Gyrase interaction were more conserved than the CcdA-interacting and non-interacting residues (Figure 5D). This is consistent with the fact that the rejuvenation rate constants for mutants at Gyrase interacting residues correlate well with their affinity of interaction with GyrA14 and also with the greater conservation of Gyrase relative to CcdA. The binding of CcdB to Gyr14 is much more sensitive to mutation than to CcdA, consistent with the smaller interface in the former.

### Probing the oligomeric states of purified proteins and complexes of different CcdB mutants by Size Exclusion Chromatography-Multi Angle Light Scattering (SEC-MALS)

The oligomeric states of WT CcdB and a few of the selected CcdB mutants in complex with GyrA14, CcdA_45-72_ peptide or both the partners, namely, GyrA14 and CcdA_45-72_ peptide, were probed using SEC-MALS (Figure S15, Table S10). The CcdB mutants Y8D, Y8L, Y8V and A69R which showed slow rejuvenation rate constants *in vitro*, as observed by both SPR and YSD, were selected for the study. These mutants showed varying affinities towards CcdA_45-72_, whereas their affinities towards GyrA14 were similar to WT. Approximately 10 µM of each of the proteins and their complexes were analysed under non-denaturing conditions by SEC-MALS in 1XPBS, pH 7.4 buffer at room temperature. The refractive index (RI) traces for all the purified proteins and *in vitro* reconstituted complexes are plotted (Figure S15). In all cases, we used CcdA_45-72_ peptide instead of the full-length CcdA as the full-length CcdA is known to degrade and form aggregates (Dao-Thi et al., 2002). Calculated molecular weights of the peaks are shown in Table S10.

In the case of purified CcdB WT (red), a single peak corresponding to the mass of dimeric CcdB with a molecular weight of 24 kDa was observed (Figure S15A). The purified GyrA14 (magenta) protein is a dimer in solution with a molecular weight of 40 kDa (Figure S15A). We also probed the oligomeric status of *in vitro* assembled CcdB WT:CcdA_45-72_ (black) and CcdB WT:GyrA14 (blue) complexes (Figure S15A). The observed stoichiometries of the *in vitro* reconstituted complexes were in concordance with that observed in their crystal structures (Figure S15A). At equimolar concentrations of the components in both complexes, namely, CcdB and CcdA_45-72_ (10 µM each) or CcdB and GyrA14 (10 µM each), we observed the formation of hetero-tetrameric complexes (Figure S15A). The *in vitro* reconstituted CcdB WT:GyrA14 complex (with 10 µM of each of the proteins) was rejuvenated with 10 µM of CcdA_45-72_ and the oligomeric status of the rejuvenated complex was determined (green). We observed that most of the CcdB:GyrA14 complexes had been rejuvenated, thereby showing two peaks corresponding to free dimeric GyrA14 and the hetero-tetrameric CcdB WT:CcdA_45-72_ complex (Figure S15A).

The purified CcdB mutants Y8D, Y8L, Y8V and A69R, all showed incomplete rejuvenation (Figure S15B-E). A single dimeric peak with a molecular weight of 24 kDa was observed for all the purified CcdB mutants in isolation (Figure S15B-E). The oligomeric status of both *in vitro* assembled CcdB:CcdA_45-72_ (black) and CcdB:GyrA14 (blue) complexes, formed with equimolar concentrations of the components, namely, CcdB and CcdA_45-72_ (10 µM each) and CcdB and GyrA14 (10 µM each), were hetero tetramers (Figure S15B-E). The *in vitro* reconstituted CcdB:GyrA14 complex (10 µM of each of the proteins) was rejuvenated with 10 µM of CcdA_45-72_ and the oligomeric status of the rejuvenated complex was determined (green). We observed that in all cases, the CcdB:GyrA14 complex had not been fully rejuvenated, therefore showing three peaks corresponding to the formation of a possible ternary complex of CcdA_45-72_:CcdB:GyrA14 or a hetero-tetrameric CcdB:GyrA14 complex, the free dimeric GyrA14 and the tetrameric CcdB:CcdA_45-72_ complex (Figure S15B-E).

Interestingly, the differences in their affinities towards CcdA_45-72_ were not correlated with the extent of rejuvenation of the CcdB mutant:GyrA14 complexes. Both Y8L and Y8V had approximately five-fold lower affinities towards CcdA_45-72_ relative to WT CcdB. However, there were differences in the peaks observed for their CcdB:GyrA14 complexes after being rejuvenated with CcdA_45-72_.Y8V displayed higher amounts of the complex being retained relative to the Y8L mutant and was associated with the absence of a peak corresponding to free GyrA14, in contrast to Y8L. Another interesting observation was that the Y8D mutant which had significantly lower affinity towards CcdA_45-72_ than Y8L and Y8V CcdB, rejuvenated to the same extent as was seen for Y8L and Y8V. In contrast, A69R which showed approximately 900-fold reduced affinity towards CcdA_45-72_, rejuvenated faster, compared to the aforementioned three mutants (Y8V, Y8L and Y8D) though slower than CcdB WT.

In all cases, the inability to distinguish between the possible ternary CcdA_45-72_-CcdB mutant-Gyrase complex and the hetero-tetrameric CcdB mutant-Gyrase complex is due to the small difference in the molecular weights of the two complexes which is beyond the resolution limit of the column. Owing to the small size of the CcdA_45-72_ peptide (∼3 kDa), SDS PAGE of the peak fractions were not performed.

### FRET studies confirm that mutants at CcdA interacting sites form ternary complexes with increased stability

To further probe the formation of probable ternary complexes in solution, FRET studies were carried out. Previously, we have used the GyraA14-E487C mutant and CcdA_50-72_ peptide for FRET studies to probe the formation of a transient ternary complex in solution for WT CcdB. In the same study, we have shown that the labelled and the unlabelled peptide rejuvenated WT CcdB from GyrA14 to the same extent (Aghera et al., 2020). Further, the rejuvenation rate constants of CcdA_50-72_ peptide were observed to be similar to that of the CcdA_37-72_ and CcdA_45-72_ peptides (Aghera et al., 2020). Since the CcdA_50-72_ peptide has a R57C mutation which can be used for labelling with a suitable fluorophore, in the present study we used this peptide for probing the formation of the ternary complex. GyrA14-E487C was labelled with Fluorescein-5-Maleimide donor (GyrA14*) and the CcdA_50-72_ peptide was labelled with 5-TAMRA acceptor (CcdA_50-72_*) to facilitate the detection of probable ternary complexes.

The rejuvenation process was initiated by mixing the GyrA14*:CcdB complex (10 nM each) with 100 nM _50-72_* and monitored by the change in acceptor (TAMRA) fluorescence on CcdA50-72*. All the CcdB mutants used in the study showed the existence of a transient ternary complex of CcdB:GyrA14*:CcdA_50-72_* (Figure 6A). Mutants at sites involved in CcdA interaction showed a much more stable ternary complex than either WT CcdB or CcdB mutants involved in GyrA14 binding (Figure 6A). Interestingly, two of the CcdB mutants S60E and R86D, at residues which do not interact with either CcdA or Gyrase showed the formation of a stable ternary complex, consistent with the YSD and SPR studies (Figure 6A). The dead time of manual mixing in all the experiments was ∼10 s, therefore, only the slow phase corresponding to the decrease in the acceptor fluorescence could be captured (Figure 6A). Expectedly, the acceptor fluorescence does not change when rejuvenation studies were carried out for a few of the CcdB mutants using 100 nM of unlabeled CcdA_50-72_ (Figure S16). In addition, no change in acceptor fluorescence was observed in CcdB:GyrA14* complexes (Figure S16). The rate of rejuvenation of all the CcdB mutants determined using FRET was also calculated (Figure 6B, Table S2). Consistent with the other studies it was observed that the mutants at CcdA interacting residues have much slower rejuvenation rate constants compared to WT or mutants at Gyrase interacting residues. The rejuvenation rate constants from FRET studies correlated very well with those obtained from either YSD or SPR (Figure 6C), thus showing the consistency of our methodologies.

**Figure 6.**
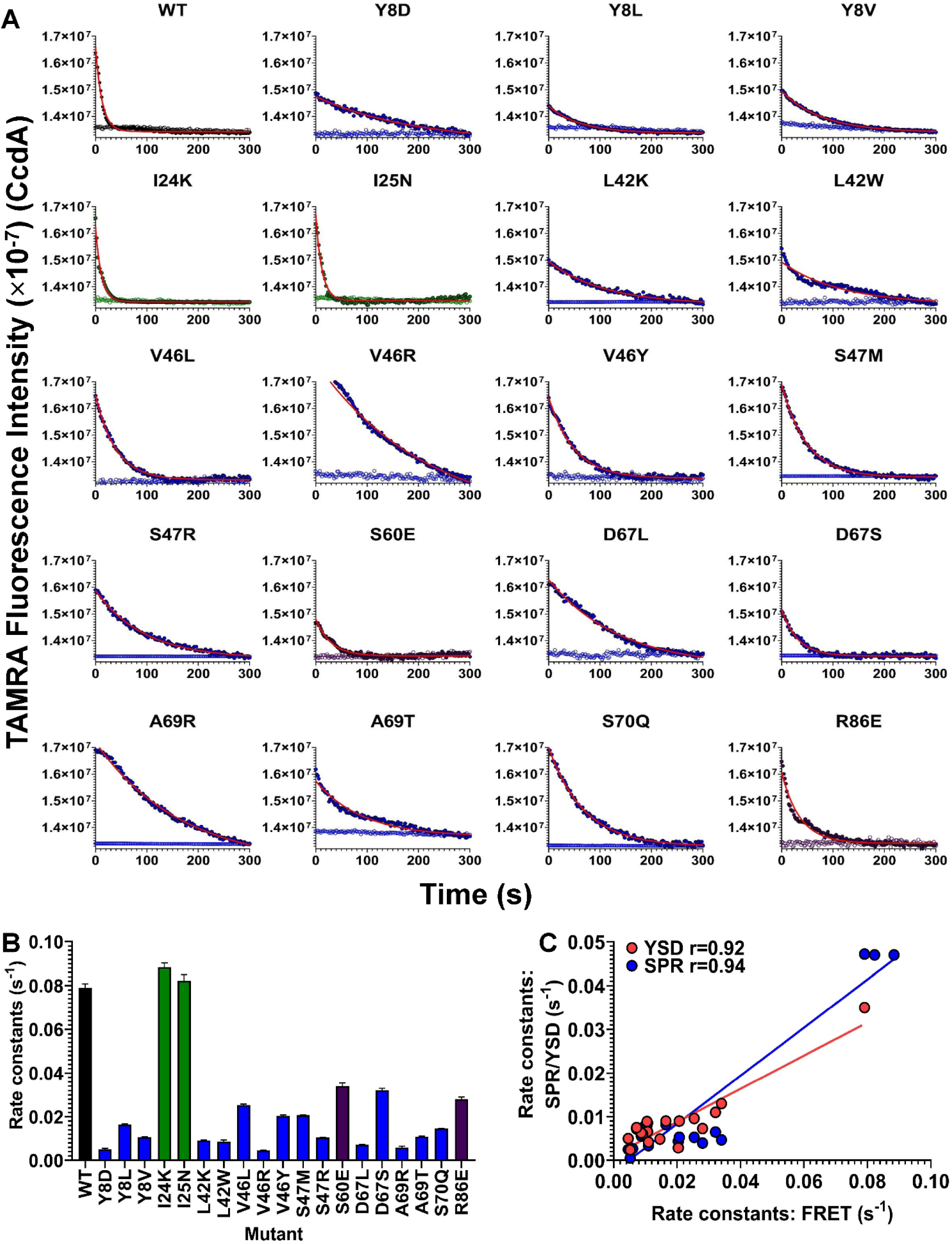
Ternary Complex Captured Using FRET. (A) Overlays show the time-dependent change in fluorescence observed upon mixing 10 nM of a preformed complex of CcdB:GyrA14* with 100 nM of CcdA_50-72_*. The rejuvenation traces of WT CcdB, CcdB mutants at residues involved in Gyrase binding, CcdB mutants at residues involved in CcdA binding and CcdB mutants that do not interact with either Gyrase or CcdA are shown in black, green, blue and magenta solid circles. The open circles in respective colours show the fluorescence of CcdB:CcdA_50-72_*. The fits of the kinetic traces are shown in solid red lines. (B) The rate constants for rejuvenation of the CcdB mutants obtained from the FRET studies. The error bars represent the standard deviation from two independent experiments, each performed in duplicates. (C) The correlation of rejuvenation rate constants obtained from FRET with those obtained from YSD (red) and SPR studies (blue). All experiments are carried out in 1XPBS, pH7.4 at 25 °C. *See also Figure S15-S16*.

## Discussion

Molecular switches, where binding of one or more ligands brings about desired functional changes in a target protein are important components of many biological processes (Schreiber, 2017). In the present study, we have carried out detailed experimental studies of one such molecular switch, CcdA, which is known to restore the function of Gyrase by facilitating dissociation of CcdB from the CcdB:GyrA14:DNA complex (Dao-Thi et al., 2005; De Jonge et al., 2009). The disordered C-terminal domain of CcdA is involved in binding to CcdB, thereby inhibiting CcdB toxicity and autoregulating the transcription of the *ccd* operon (Burger et al., 2017; Drobnak et al., 2013; Madl et al., 2006).

In a recent study, it was demonstrated that binding of CcdA to CcdB:GyrA14 results in the formation of short-lived, transient ternary and quaternary complexes of GyrA14, CcdB and CcdA (Aghera et al., 2020). In the same study, the molecular mechanism involved in the rejuvenation process was probed by a combination of FRET, HX-MS, NMA and PRS studies (Aghera et al., 2020). The binding of the 61-72 segment of CcdA to CcdB induces the vital structural and dynamic changes which are required to facilitate the dissociation of CcdB from GyrA14. The dissociation process is further enhanced by the 50-60 segment of CcdA through additional allosteric effects. Finally, GyrA14 rebinding to CcdB is prevented by the 37-49 segment of CcdA (Aghera et al., 2020).

In the current study, we first investigated the role of different lengths of CcdA on the rejuvenation rate constant of WT CcdB from the CcdB-GyrA14 complex. Consistent with the previous study (Aghera et al., 2020), we observe that CcdA peptides of different lengths (CcdA_37-72_, CcdA_45-72_) have similar *in vitro* effects on the rejuvenation rate constants of WT CcdB as full-length CcdA, except for CcdA_61-72_ which fails to rejuvenate.

The present study utilises YSD coupled to Sanger sequencing to quantitate mutational effects on rejuvenation kinetics, and to isolate stabilised ternary complexes in a high throughput fashion, using the CcdA_45-72_:CcdB:GyrA14 system. While YSD is commonly used to isolate high affinity binder and estimate affinities (!refs), there are few studies which employ this for kinetic measurements. Following two rounds of sorting and enrichment, CcdB mutants having both significantly slower and faster rejuvenation rate constants than the WT were isolated. The mutants were characterised extensively to determine the probable mechanisms affecting the rates of rejuvenation and the rejuvenation kinetics measured by YSD were validated by characterising rejuvenation of corresponding purified proteins by SPR as well as FRET. In general, it was observed that the mutants defective in or having reduced affinities towards CcdA rejuvenated slower than the WT CcdB. Further, a few selected mutants were allowed to rejuvenate for longer time points. It was observed that most mutants at residues involved in CcdA binding (Y8D, Y8L, Y8V, L42K, V46R, M64E, M64R, T66E, A69R) show significant GyrA14 binding signals even after 200 seconds of rejuvenation, as compared to WT CcdB, except for S12G and R13F. In contrast, mutants with reduced affinities towards Gyrase rejuvenated significantly faster than the WT. Weak to strong quantitative correlation was observed between the binding affinities for CcdA_45-72_ and GyrA14 respectively, and the observed rejuvenation rate constants. It was observed that amongst the CcdA interacting regions, mutations at residues 60-71 had the strongest effect on the CcdA binding affinity followed by mutations at the 8-14 loop. Mutations at GyrA14 and CcdA/GyrA14 non-interacting sites did not affect the CcdA binding affinities, relative to WT CcdB. In the CcdA interacting regions, most types of substitutions had a similar fold reduction in CcdA affinity, except in certain cases, where aromatic (S12W, P35W, V46Y, A69W) or charged (V46R, M64R, A69R) substitutions cause a drastic change in CcdA binding as measured by both YSD and SPR. It was observed that the CcdB residues involved in GyrA14 binding are evolutionarily more conserved than the CcdA interacting residues. The surface area buried during complex formation is uncorrelated with mutational effects on rejuvenation rate.

Mutants in different regions of CcdB which are either involved in CcdA or GyrA14 binding or at residues non-interacting with either of them, were also purified and their rejuvenation rate constants were determined by SPR. The rejuvenation rate constants of CcdA interacting-site mutants were strongly affected by mutations, with significant effect observed for certain residues in the 8-15, 30-40, 41-52 and 65-72 residue regions. Amongst these CcdA interacting regions, residues 65-72 and the loop residues from 40-50 have the maximum mutational effect on rejuvenation rate constants as compared to mutations in residues 8-15 and 30-40. This is consistent with our previous computational study using perturbation response scanning (PRS), where it was shown that binding of CcdA residues 61-72 to residues 67-72 of CcdB was a crucial event, as residues in the 67-72 region of CcdB act as effectors and induce major conformational changes that facilitate CcdA binding (Aghera et al., 2020). In order to further delineate the effect of different substitutions on rejuvenation rate constants, the mutations at CcdA interacting positions were also classified into different categories of amino acids, namely, aliphatic, aromatic, charged and polar. For all concentrations of CcdA peptide used, it was observed that different classes of substitutions have similar normalised rejuvenation rate constants.

While, mutations at GyrA14 interacting sites have faster rejuvenation rate constants than WT CcdB, the mutants at CcdA/GyrA14-non-interacting sites did not affect rejuvenation, relative to that for the WT CcdB, except for a few mutants such as S60E and R86E (Table 1). This is of particular interest as these mutations themselves do not have any effect on CcdA or Gyrase binding affinities, however, they may cause structural perturbations in loops involved in CcdA binding which in turn affect the rejuvenation process. In a recent study, we solved the structure of the S60E mutant, (Chattopadhyay et al., 2021). The mutant E60 is involved in a number of new salt-bridge interactions with residues R48, H55 and R62 which may hinder CcdA interaction when bound to Gyrase (Chattopadhyay et al., 2021). In addition, certain mutants at sites that are not involved in interaction with either CcdA or GyrA14 such as K9R, E11P, D44V, R48T, D58N, S60G and R86D have rejuvenation rate constants five to twenty folds faster than the WT. The effect of these substitutions is discussed in Table 1. Most of these mutations are involved in several intramolecular non-covalent interactions in the CcdA bound context (PDB ID: 3G7Z). Mutations at positions K9R and E11P, though not involved directly in CcdA interaction, have an impact on the rejuvenation rate constants because of the importance of the 8-14 CcdA interacting loop in the early events of CcdA binding during the rejuvenation process. K9 is involved in H-bonds with V75 and S76 and therefore the mutation to arginine may cause local distortion of the 8-14 loop. E11P likely results in a kink in the 8-14 loop thus affecting the rejuvenation process. D44 and R48 are involved in an intramolecular salt bridge interaction with each other and D58S is involved in a salt bridge interaction with H55 that stabilizes the 39-50 loop in the CcdA bound form. The D44V, R48T and D58N mutations likely affect the loop conformation, thus impacting the rejuvenation process. The impact of S60G on the rejuvenation rate constants may result from local structural rearrangements caused due to the breaking of the H-bond of S60 with A39 and H55. The R86 position is involved in a series of salt bridge interactions with D89 and H-bonds with V53, l83, D89 and I90 which will all be disrupted by the R86E mutation. This mutation also results in a 6-fold decrease in GyrA14 affinity.

**Table 1.**
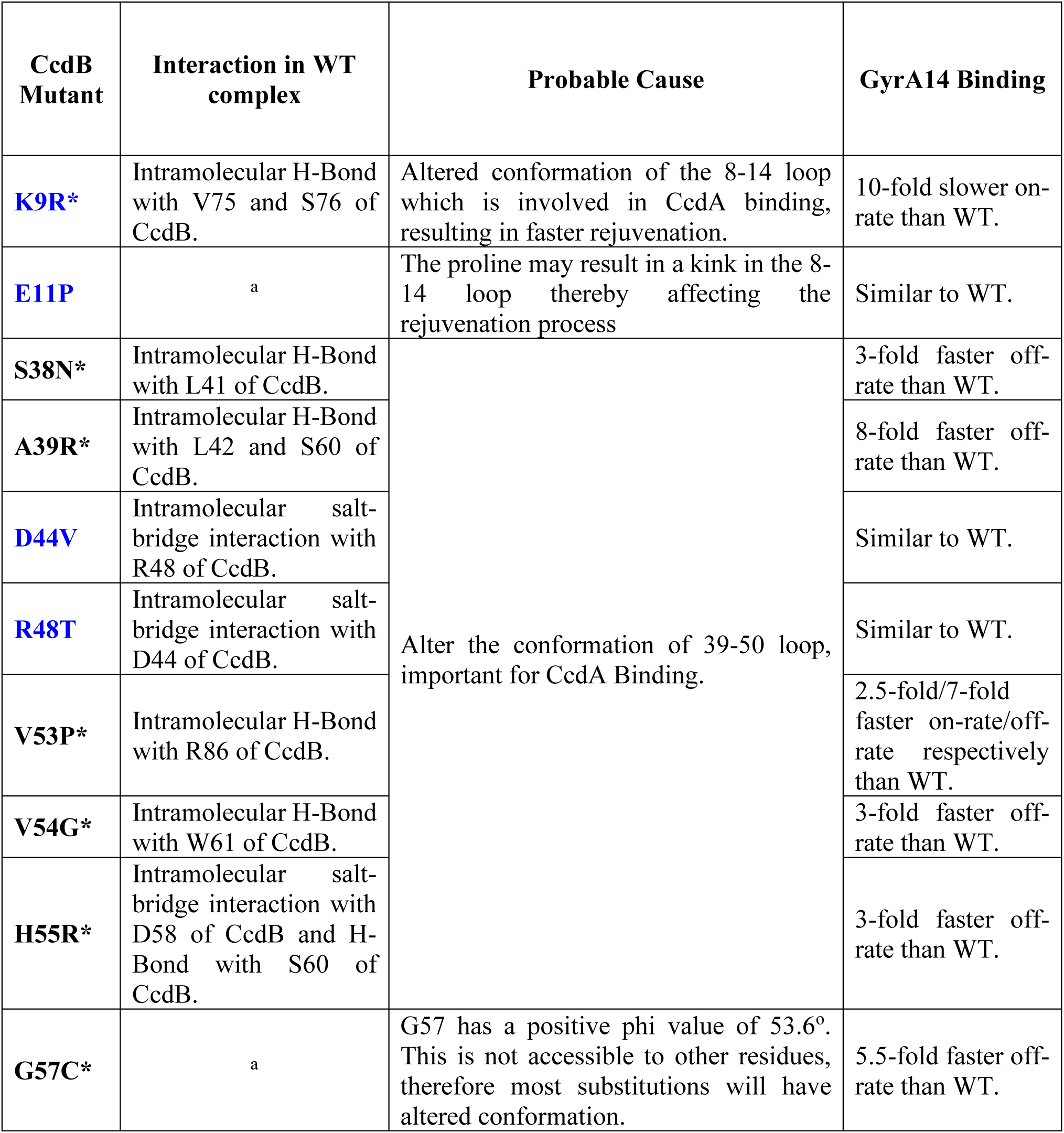
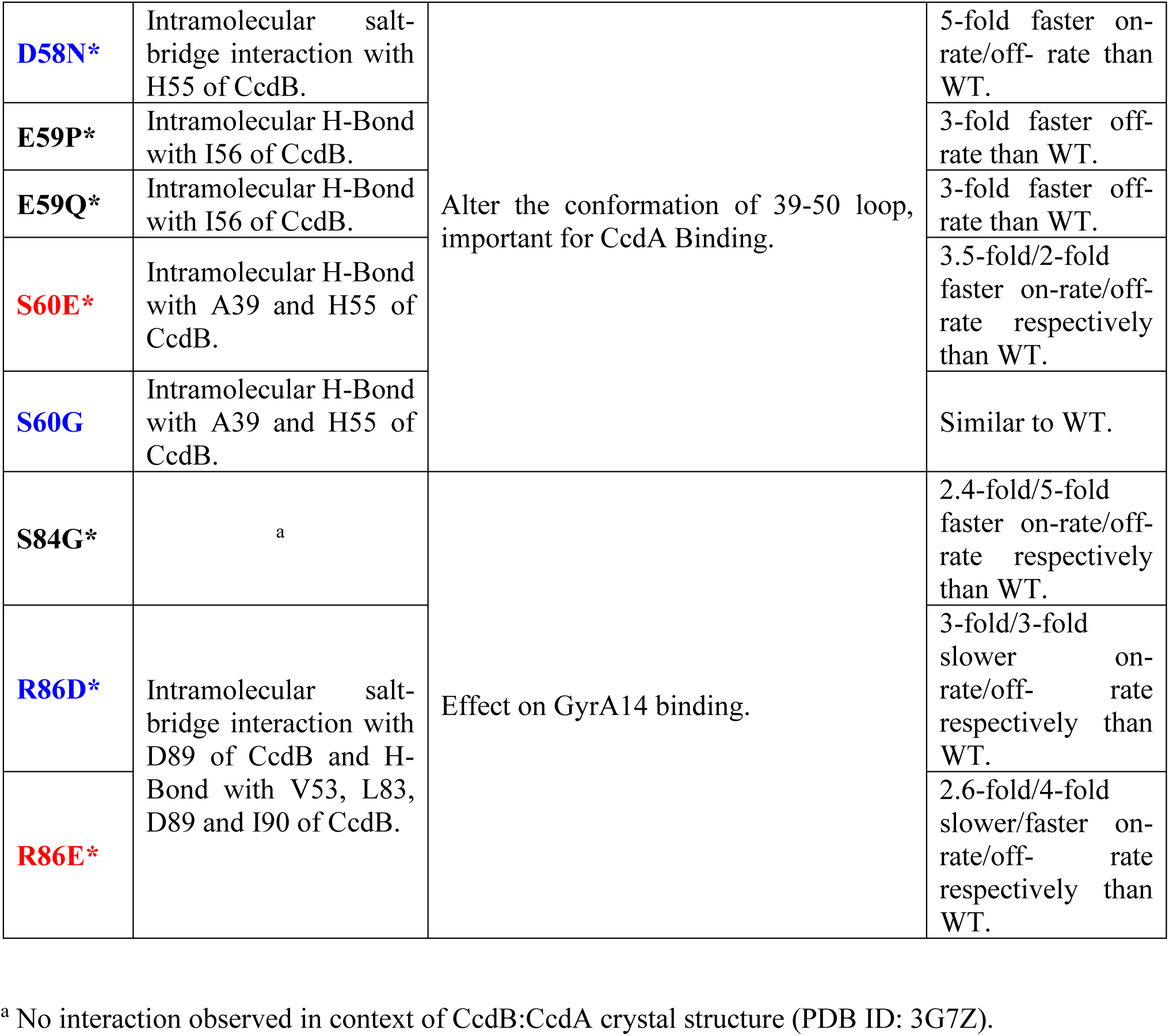
Interaction analysis of CcdB mutants in CcdA bound conformation (PDB ID: 3G7Z), that neither interact with CcdA nor with GyrA14 but show significant effect on measured rejuvenation rate constants. Mutants with significantly slower and faster rejuvenation rate than WT are highlighted in red and blue boxes respectively and mutants with altered binding to GyrA14 are marked with a star (*).

| CcdB Mutant | Interaction in WT complex | Probable Cause | GyrA14 Binding |
| --- | --- | --- | --- |
| **K9R\*** | Intramolecular H-Bond with V75 and S76 of CcdB. | Altered conformation of the 8-14 loop which is involved in CcdA binding, resulting in faster rejuvenation. | 10-fold slower on-rate than WT. |
| **E11P** | <sup>a</sup> | The proline may result in a kink in the 8-14 loop thereby affecting the rejuvenation process | Similar to WT. |
| **S38N\*** | Intramolecular H-Bond with L41 of CcdB. | Alter the conformation of 39-50 loop, important for CcdA Binding. | 3-fold faster off-rate than WT. |
| **A39R\*** | Intramolecular H-Bond with L42 and S60 of CcdB. |  | 8-fold faster off-rate than WT. |
| **D44V** | Intramolecular salt-bridge interaction with R48 of CcdB. |  | Similar to WT. |
| **R48T** | Intramolecular salt-bridge interaction with D44 of CcdB. |  | Similar to WT. |
| **V53P\*** | Intramolecular H-Bond with R86 of CcdB. |  | 2.5-fold/7-fold faster on-rate/off-rate respectively than WT. |
| **V54G\*** | Intramolecular H-Bond with W61 of CcdB. |  | 3-fold faster off-rate than WT. |
| **H55R\*** | Intramolecular salt-bridge interaction with D58 of CcdB and H-Bond with S60 of CcdB. |  | 3-fold faster off-rate than WT. |
| **G57C\*** | <sup>a</sup> | G57 has a positive phi value of 53.6°. This is not accessible to other residues, therefore most substitutions will have altered conformation. | 5.5-fold faster off-rate than WT. |
| **D58N\*** | Intramolecular salt-bridge interaction with H55 of CcdB. | Alter the conformation of 39-50 loop, important for CcdA Binding. | 5-fold faster on-rate/off-rate than WT. |
| **E59P\*** | Intramolecular H-Bond with I56 of CcdB. |  | 3-fold faster off-rate than WT. |
| **E59Q\*** | Intramolecular H-Bond with I56 of CcdB. |  | 3-fold faster off-rate than WT. |
| **S60E\*** | Intramolecular H-Bond with A39 and H55 of CcdB. |  | 3.5-fold/2-fold faster on-rate/off-rate respectively than WT. |
| **S60G** | Intramolecular H-Bond with A39 and H55 of CcdB. |  | Similar to WT. |
| **S84G\*** | <sup>a</sup> | Effect on GyrA14 binding. | 2.4-fold/5-fold faster on-rate/off-rate respectively than WT. |
| **R86D\*** | Intramolecular salt-bridge interaction with D89 of CcdB and H-Bond with V53, L83, D89 and I90 of CcdB. |  | 3-fold/3-fold slower on-rate/off-rate respectively than WT. |
| **R86E\*** |  |  | 2.6-fold/4-fold slower/faster on-rate/off-rate respectively than WT. |

<sup>a</sup> No interaction observed in context of CcdB:CcdA crystal structure (PDB ID: 3G7Z).

Interestingly some of the other mutations outside the GyrA14 binding site also have an impact on GyrA14 affinity, namely S38N, A39R, V53P, V54G, G57C, E59P, E59Q and S84G. Most of these positions are again involved in a number of non-covalent interactions with R86E, which is in turn involved in a series of interactions with residues involved in Gyrase binding, thereby indirectly causing an effect in the binding kinetics. G57 has a positive phi value of 54 that is only accessible to Glycine. Therefore, substitutions to other amino acids at this position will result in altered loop conformation.

At all CcdA interacting positions aromatic and charged substitutions showed the largest effects on rejuvenation rate constants. This was consistent with the YSD results, where substitutions such as Y8D, Y8E, S12W, P35W, V46R, V46Y, M64E, M64R, T66E, A69R and A69W strongly affected rejuvenation rate constants. Regions of CcdB, such as the 8-15 loop and 65-72 region, which interacts with CcdA_61-72_ are accessible in the Gyrase bound form and facilitate the early binding of CcdA, which in turn induces conformational change in CcdB. As residues 61-72 of CcdA are also rich in charged (D67, E68, R70, D71) and aromatic amino acids (F65, W72), substitution to the same categories of amino acids in the CcdA interacting regions of CcdB may cause charge-charge repulsion or steric hindrance due to the introduction of bulky aromatic amino acids, thus slowing down the rejuvenation process.

The formation of probable ternary complexes for a few of the CcdB mutants with slower rejuvenation rate constants was also probed by FRET. Mutants at critical regions of CcdB which facilitate either CcdA or GyrA14 binding and rejuvenation, and also showed the strongest effect on the rejuvenation rate constants in both SPR and YSD studies, were chosen for this study. It was observed that CcdB mutants with slower rejuvenation rate constants, that were at positions largely involved in CcdA binding, formed a stable ternary complex relative to WT CcdB.

In summary, we describe novel methodology that can be used to measure rejuvenation kinetics in a high throughput fashion. The results obtained from YSD also correlated well with SPR and FRET experiments, validating this novel use of YSD for kinetic measurements. The methodology was used to provide insights into the rejuvenation process and identify mutations that stabilized otherwise highly transient species. In this proof of principle study, down selected mutants were individually cloned and characterized, but in future, mutant pools isolated by FACS can be analysed by deep sequencing to further enhance throughput (Adams et al., 2016; Ahmed et al., 2022b, 2022a; Noderer et al., 2014; Reich et al., 2015; Starr et al., 2020).

## Materials and Methods

### Plasmids and host strains

The *ccdB* gene was cloned under control of the P_BAD_ Promoter in pBAD24 vector (Bajaj et al., 2004). *Escherichia coli* host strain *Top10Gyrase*, resistant to the action of CcdB toxin (Bernard and Couturier, 1992; Tripathi et al., 2016), was used for expression and purification of the CcdB mutant proteins. GyrA14 gene was cloned under control of the T7 promoter in pET-15b vector. *Escherichia coli* host strain BL21 (DE3) pLysE, a T7 RNA pol expression strain that is deficient in Lon and OmpT proteases, was used for expressing GyrA14 protein. The *Saccharomyces cerevisiae* strain EBY100 was used for yeast surface display to monitor the binding and expression of the displayed proteins cloned in the yeast surface display vector pPNLS. (Chao et al., 2006).

### Cloning of WT and ccdB mutants

For all the single mutants, the *ccdB* gene was amplified in two fragments with the desired point mutations. The fragments had overlapping regions (introduced during PCR) of 25-28 nucleotides, which were then recombined with pET-15b vector for protein purification using Gibson assembly (Gibson et al., 2009) or *in vivo* recombined with pPNLS vector for YSD as described earlier (Chandra et al., 2021). Amplification was done using Phusion Polymerase from NEB as per the manufacturer’s protocol.

### Expression and purification of WT and CcdB mutants

The WT and the CcdB mutants were expressed from the P_BAD_ promoter in the pBAD24 vector in the CcdB resistant Top10Gyrase strain of *E. coli*. The purification of the CcdB mutants were carried out as described previously (Chattopadhyay and Varadarajan, 2019). 500 mL of Luria bertani medium (HiMedia) was inoculated with 1% of the primary inoculum, grown at 37 °C till OD_600_=0.6, followed by induction with 0.2% (w/v) arabinose and grown at 37 °C for 5 hours. The harvested cells were re-suspended in HEG re-suspension buffer pH 7.4 (10 mM HEPES, 50 mM EDTA, 10% glycerol containing 10 mM PMSF) and lysed by sonication. The supernatant was incubated with CcdA peptide (residues 46-72) coupled Affi-gel15 (Biorad) overnight at 4 °C. The unbound fraction was removed, washed with five column volumes of coupling buffer pH 8.3 (0.05 M Sodium Bicarbonate, 0.5 M Sodium Chloride) and eluted with 0.2 M Glycine, pH 2.5 into a tube containing an equal volume of 400 mM HEPES, pH 8.4, 4 °C (Chattopadhyay and Varadarajan, 2019; Tripathi et al., 2016). The eluted fractions were subjected to 15% Tricine SDS-PAGE and the protein concentration was determined. Fractions containing pure protein were pooled and stored at −80 °C. For SPR, and FRET experiments the CcdB proteins were buffer exchanged in 1XPBS, pH7.4.

### Biotinylating full-length CcdA and CcdA peptides of different lengths

The full-length CcdA and CcdA peptides of different lengths (Table S1) were biotinylated with EZ-link^TM^ NHS-Biotin (ThermoFisher Scientific) as per the manufacturer’s protocol. Briefly, biotinylation was carried out using a 2-3 fold molar-excess of biotin, followed by desalting with a PD MiniTrap G-10 column (GE Healthcare) to remove the excess unreacted biotin. Biotinylation was further confirmed by mass spectrometry.

### Sorting and enrichment of the CcdB library at different time points

An SSM library of ccdB generated previously in the pBAD24 vector (Adkar et al., 2012; Tripathi et al., 2016) was PCR amplified with primers that also had extensions homologous to the pPNLS vector. The amplified library was then gel purified and recombined with pPNLS vector *in vivo* using the *Saccharomyces cerevisiae* EBY 100 strain as described earlier (Ahmed et al., 2022a). Yeast cells containing libraries were grown and induced for protein expression as explained previously (Ahmed et al., 2022a). Briefly, yeast cells expressing the CcdB library were incubated with the cognate partner, 100 nM GyrA14 for 1 hour at 4 °C with shaking. The cells were then washed twice with 200 µL PBS (with 0.5% BSA). This was followed by addition of 10 nM or 100 nM biotinylated CcdA_45-72_ peptide to isolate CcdB mutants with rejuvenation rate constants faster or slower than the WT CcdB respectively. The reaction was quenched by adding 10 fold molar excess of purified CcdB WT protein over CcdA, after 5 s for isolating mutants which are more sensitive to rejuvenation than WT, and after 30 s to isolate mutants which are more resistant to rejuvenation than WT. GyrA14 binding was probed by mouse anti-FLAG primary antibodies followed by secondary rabbit anti-mouse antibodies conjugated to Alexa Fluor 633. CcdA binding was probed by Streptavidin-PE. For enriching mutants with faster rejuvenation rate constants, after two rounds of sorting, the populations having higher bound CcdA signal than WT were sorted. For enriching mutants with slower rejuvenation rate constants, after two rounds of sorting, the populations having higher bound GyrA14 signal than WT were sorted. After sorting, the sorted cells were grown and plasmid pools were isolated, purified and transformed in *E.coli Top 10Gyrase* cells. The colonies obtained were Sanger sequenced for mutant identification.

### Analysis of expression, GyrA14 and CcdA binding and rejuvenation of the identified CcdB mutants by YSD

The individual CcdB mutants identified from Sanger sequencing were transformed into *Saccharomyces cerevisiae* EBY100 cells as explained earlier (Ahmed et al., 2022a). The transformed CcdB mutants were grown in SDCAA media (glucose 20 g/L, yeast nitrogen base 6.7 g/L, casamino acid 5 g/L, citrate 4.3 g/L, sodium citrate dihydrate 14.3 g/L) for sixteen hours at 30 °C, 250 rpm. Secondary inoculation was done by inoculating cells from the overnight culture till a final OD_600_ of 0.2 was reached in SDCAA media. Following secondary inoculation, cells were grown for six hours and reinoculated in induction media (SGCAA media; galactose 20 g/L, glucose 2 g/L, yeast nitrogen base 6.7 g/L, casamino acid 5 g/L, citrate 4.3 g/L, sodium citrate dihydrate 14.3g/L) to attain a final OD_600_ of 0.5. Induction was done for 16 hours at 30 °C, 250 rpm. Ten million cells were used for FACS sample preparation. The amount of expression of proteins on the yeast cell surface was estimated by incubating the induced cells in 20 µL FACS buffer (1X PBS and 0.5% BSA), containing chicken anti-HA antibodies from Bethyl labs (1˸600 dilution), for 30 minutes at 4 °C. This was followed by washing the cells twice with FACS buffer. Washed cells were incubated with 20 µL FACS buffer containing goat anti-chicken antibodies conjugated to Alexa Fluor 488 (1:300 dilution), for 20 minutes at 4 °C. Fluorescence of yeast cells was measured by flow cytometric analysis. The GyrA14 binding activity on the yeast cell surface was estimated by incubating the induced CcdB mutants in 20 µL FACS buffer containing 100 nM FLAG tagged GyrA14, followed by washing with FACS buffer and incubation with 20 µL mouse anti-FLAG antibodies, at a dilution ratio of 1˸300. This was followed by washing the cells twice with FACS buffer, and 20 µL rabbit anti-mouse antibodies conjugated to Alexa Fluor 633 (1:1600 dilution) prepared in FACS buffer was added. The amount of bound CcdA_45-72_ peptide on the yeast cell surface was estimated by incubating the induced CcdB mutants in 25 µL FACS buffer containing 1 nM biotinylated CcdA, followed by washing with FACS buffer and incubation with 20 µL Streptavidin-PE antibody, at a dilution ratio of 1˸1000. This was followed by washing the cells twice with FACS buffer. The rejuvenation rate constants of CcdB mutants were measured by incubating them first with GyrA14, followed by addition of 10 nM biotinylated CcdA_45-72_ peptide for 5 s or 100 nM biotinylated CcdA_45-72_ peptide for 30 s for isolating mutants with faster or slower rejuvenation rate constants than WT respectively. The reaction at each time point was quenched by adding 10 fold molar excess of purified CcdB WT protein over CcdA. The flowcytometry was done using a BD Aria III and the data were analysed as discussed below.

### Measurement of rejuvenation rate constants of CcdB mutants from YSD

The CcdB mutants were grown and induced for protein expression as explained above. Briefly, the CcdB mutants were incubated with the cognate partner, 100 nM GyrA14, for 1 hour at 4 °C with shaking. The cells were then washed twice with 100 µL PBS (0.5% BSA). This was followed by addition of 10 nM or 100 nM biotinylated CcdA_45-72_ peptide for CcdB mutants having faster or slower rejuvenation rate constants than WT respectively. The reaction was quenched by adding 10 fold molar excess of purified CcdB WT protein over CcdA_45-72_ after 10, 20, 30, 40 and 50 s. Rejuvenation rate constants were calculated based on the decrease in the Mean Fluorescence Intensity (MFI) of GyrA14 binding which was probed as explained in Section 5.2.6. The MFI of GyrA14 Binding at each time point was normalised with the MFI of GyrA14 Binding observed at 10 s for relative comparisons across mutants. The data was then fit using the equation Y=(Y_0_)*exp(-k*X) in GraphPad Prism version 8.0, where Y is the MFI of GyrA14 Binding at each time point X in s, Y_0_ is the maximum MFI of GyrA14 Binding at time 10 s and k is the rejuvenation rate constant in s^-1^. During analysis, we observed that within 10 s of rejuvenation there was a large decrease in GyrA14 binding, after which the reduction was significantly slower. Given that the dead time of manual mixing is a few seconds, we excluded the data of GyrA14 binding for the first 10 seconds.

### Measurement of CcdA_45-72_ peptide affinity of CcdB mutant proteins by YSD

The affinities of CcdB mutants for CcdA_45-72_ were measured using YSD as described earlier (Chandra et al., 2021). The CcdB mutants were expressed on the yeast cell surface and the levels of expression were measured as described previously (Ahmed et al., 2022a). Binding was measured by incubating yeast cells expressing these mutants with different concentrations of their cognate partner CcdA_45-72_ (1 fM to1 µM). Binding was measured on a BD Accuri FACS machine and the binding MFI values were fit against the concentration of ligand to estimate the dissociation constant, K_D_, using the equation Y=B_max_*X/(K_D_ + X) in GraphPad Prism version 8.0, where Y is the MFI of CcdA_45-72_ Binding at each CcdA_45-72_ concentration X in nM, B_max_ is the maximum MFI of CcdA_45-72_ Binding and K_D_ is the equilibrium dissociation constant in nM.

### Measurement of GyrA14 binding affinities and rejuvenation rate constants of the CcdB mutant proteins by Surface Plasmon Resonance (SPR)

All the SPR experiments were performed with a Biacore 3000 (Biacore, Uppsala, Sweden) optical biosensor at 25 °C as described previously (Chattopadhyay et al., 2021). GyrA14 was immobilized at a 30 µL/min flow rate for 180s. 3000 response units of GyrA14 was covalently linked by standard amine coupling to the surface of a research-grade CM5 chip. A sensor surface (without GyrA14) that had been activated and deactivated served as a negative control for each binding interaction. For affinity measurements with GyrA14, different concentrations of the CcdB mutants were passed across each sensor surface in a running buffer of PBS (pH 7.4) containing 0.005% Tween surfactant. Protein concentrations ranged from 1 nM to 2 µM. Both association and dissociation were measured at a flow rate of 30 µL/min. For the rejuvenation experiments, the SPR was done in co-inject mode with 50 nM of CcdB WT or mutant proteins, where the association was allowed for 100 s, followed by immediate dissociation with different concentrations of the CcdA_45-72_ peptide. In all cases, the sensor surface was regenerated between binding reactions by one to two washes with 4 M MgCl_2_ for 30 s at a flow rate of 30 µL/min. Each binding curve was corrected for nonspecific binding by subtraction of the signal obtained from the negative control flow cell. The kinetic parameters were obtained by fitting the data to a simple 1:1 Langmuir interaction model by using BIA EVALUATION 3.1 software. The rejuvenation rate constants, k_d_, obtained at each concentration (10, 25, 100 and 200 nM) were normalised with respect to the rejuvenation rate constant of WT obtained at the same concentration as described below:

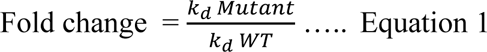

### ΔASA and Conservation score calculations

Crystal structures were used to identify CcdB residues interacting with CcdA (PDB ID: 3G7Z) (De Jonge et al., 2009) and Gyrase (PDB ID: 1X75) (Dao-Thi et al., 2005). ΔASA (CcdA) calculated using NACCESS (Hubbard & Thornton, 1993), the difference between the solvent accessible surface areas of a CcdB residue in its free (PDB ID: 3VUB) (Loris et al., 1999) and CcdA-bound form (3G7Z), was used for identifying CcdA-interacting residues. Similarly ΔASA (GyrA), the difference between the solvent accessible surface areas of a CcdB residue in its free (3VUB) and GyrA14 bound form (1X75), was used for identifying GyrA14 interacting residues. The evolutionary conservation score was calculated from the online ConSurf server which uses a multiple sequence alignment of homologous sequences to build a phylogenetic tree (Ashkenazy et al., 2016). The minimum and maximum percentage identity cut-off used was 30% and 95% respectively for the multiple sequence alignment of the toxin CcdB.

### SEC-MALS (Size exclusion chromatography-multi angle light scattering) to probe the oligomeric states of the complexes

Free CcdB, free GyrA14, CcdB:GyrA14, CcdB:CcdA_45-72_ and CcdB:GyrA14 rejuvenated with CcdA_45-72_ were separated on a Superdex-75 analytical gel filtration column (GE Healthcare) equilibrated in 1XPBS, pH 7.4 and with in-line UV (SHIMADZU), MALS (mini DAWN TREOS, Wyatt Technology Corporation) and refractive index (RI) detectors (WATERS24614) for molecular weight, aggregation and oligomerization analysis. For each run, 70 µg of the individual protein or protein complex was injected and eluted at a flow rate of 0.4 mL/min. UV, MALS and RI data were collected and analysed using ASTRA^TM^ software (Wyatt Technology) as described previously (Sharma et al., 2020).

### Purification and Labeling of GyrA14-E487C and CcdA_50-72_ peptide

The cysteine GyrA14-E487C mutant was expressed from the T7 promoter in the pET15b vector in the BL21DE3* *E. coli*. The purification of the GyrA14-E487C mutant were carried out as described previously (Aghera et al., 2020). The culture was induced at 20 °C overnight by adding 1 mM IPTG at OD_600_=0.8. The harvested cells were re-suspended in lysis buffer (1XPBS, 0.5 mM EDTA, 0.5 M Sucrose, 1 mM DTT, 10 mM PMSF, 10% glycerol, pH 7.4) and lysed by sonication. The supernatant was incubated with Ni-NTA beads for 2hours at 4 °C. The unbound fraction was removed, washed with twenty-five column volumes of wash buffer (1XPBS, 50 mM imidazole, 1 mM DTT, 10% glycerol, pH 7.4) and eluted with 1mL of elution buffer (1XPBS, 10% glycerol, 1 mM DTT and a gradient of imidazole (100 mM-900 mM, pH 7.4). The eluted fractions were buffer exchanged in 1XPBS, pH7.4 and stored at −80 °C.

The GyrA14-E487C and the CcdA_50-72_ peptide (Table S1) were labeled with Fluorescein-5-Maleimide and 5-TAMRA (Tetramethylrhodamine-5-Maleimide) from ThermoFisher Scientific as per the manufacturer’s protocol as described previously (Aghera et al., 2020). Briefly, labelling was carried out using a 5 fold molar-excess of the fluorophores, followed by desalting with a PD MiniTrap G-10 column (GE Healthcare) to remove the excess unreacted label. Labelling was further confirmed by mass spectrometry.

### Rejuvenation Kinetic Fluorescence Studies using FRET

The fluorescence measurements were carried out as described previously (Aghera et al., 2020) using a Fluoromax-3 spectrofluorometer (Horiba). To perform the rejuvenation studies, 10 nM of fluorescein labeled GyrA14-E487C (GyrA14*) was mixed with 10 nM CcdB and incubated for 30 minutes at room temperature to form the CcdB-GyrA14* complex. The rejuvenation of GyrA14* was initiated by mixing it with TAMRA labelled CcdA_50-72_-R57C (CcdA_50-72_*) at a final concentration of 100 nM. All the studies were carried out in 1XPBS, pH7.4 at 25 °C. The kinetics was measured by monitoring the fluorescence of the acceptor fluorophore on CcdA_50-72_* using excitation at 490 nm, 1 nm bandwidth and emission at 570 nm, 5 nm bandwidth using an integration time of 1 second with a data pitch of 1 nm for 300 seconds. The data for the rejuvenation rate constants obtained by FRET, was analyzed using Sigmaplot^TM^ for Windows^TM^ scientific graphing software and plots were fitted to a 3 parameter equation for exponential decay for refolding (y=y0+a*exp(-k*x)), where k is the rejuvenation rate constant (s^-1^) and x is the time of decay in seconds. We also monitored the fluorescence as a function of time of the CcdB:GyrA14* and CcdB:GyrA14* rejuvenated with unlabeled CcdA_50-72_ as controls.

## Acknowledgements

This work was funded by grants to RV from the Department of Biotechnology, grant number-BT/COE/34/SP15219/2015, DT.20/11/2015) and a JC Bose Fellowship from the Department of Science and Technology. We also acknowledge funding for infrastructural support from the following programs of the Government of India: DST FIST, UGC Centre for Advanced study, Ministry of Human Resource Development (MHRD), and the DBT IISc Partnership Program. The funders had no role in study design, data collection and interpretation, or the decision to submit the work for publication. G.C. acknowledges Ministry of Human Resource Development (MHRD) for his fellowship. S.A. is thankful to Department of Biotechnology (BT/IN/EU-INF/15/RV/19-20) for his research fellowship. Dr. Pankaj Jain is acknowledged for providing the CcdB library. We also thank all the members of RV lab for their valuable suggestions.

## Author Contributions

R.V., G.C. and S.A. designed the experiments. G.C. cloned the CcdB mutants in pPNLS vector. S.A. cloned the CcdB mutants in pBAD24 vector and purified the CcdB mutants. G.C. and S.A. performed the analysis of CcdB mutants on YSD and the library sorting. G.C. performed the SPR binding study with GyrA14, rejuvenation studies on SPR and SEC-MALS. G.C. purified and labelled the Gyrase-E487C and performed the FRET experiments. G.C. and S.A. determined the rejuvenation rate constants and affinity with CcdA peptide of the CcdB mutants on YSD. N.S.S. helped with the SPR experiments. A.A. assisted with the FACS experiments. G.C. analyzed the data for all the experiments. R.V and G.C. wrote most of the manuscript with critical inputs and review from all other authors.

## Data and Materials availability

This study did not generate any unique dataset or code. The data relevant to the figures in the paper have been made available within the article and in the supplementary information section. Further information and requests for resources and reagents should be directed to and will be fulfilled by the Lead Contact, Prof. Raghavan Varadarajan. Lead contact for the material availability: Prof. Raghavan Varadarajan. All unique/stable reagents generated in this study are available from the Lead Contact without restriction.

## Statistical analysis

All the experiments are carried out in biological replicates (n=2), and the listed errors are the standard error derived from the values obtained for individual replicates. The SPR experiments have been performed once at each concentration with four different concentrations of the protein and the listed error is the standard error derived from the values at multiple concentrations. The P values for comparing the rejuvenation parameters, were analysed with a two-tailed Mann Whitney test using the GraphPad Prism software 8.0.0 (P value indicated with ****,***,**,* indicates < 0.0001, <0.0005, <0.005, <0.1 respectively).

## Supplementary Figures

**Figure S1.**
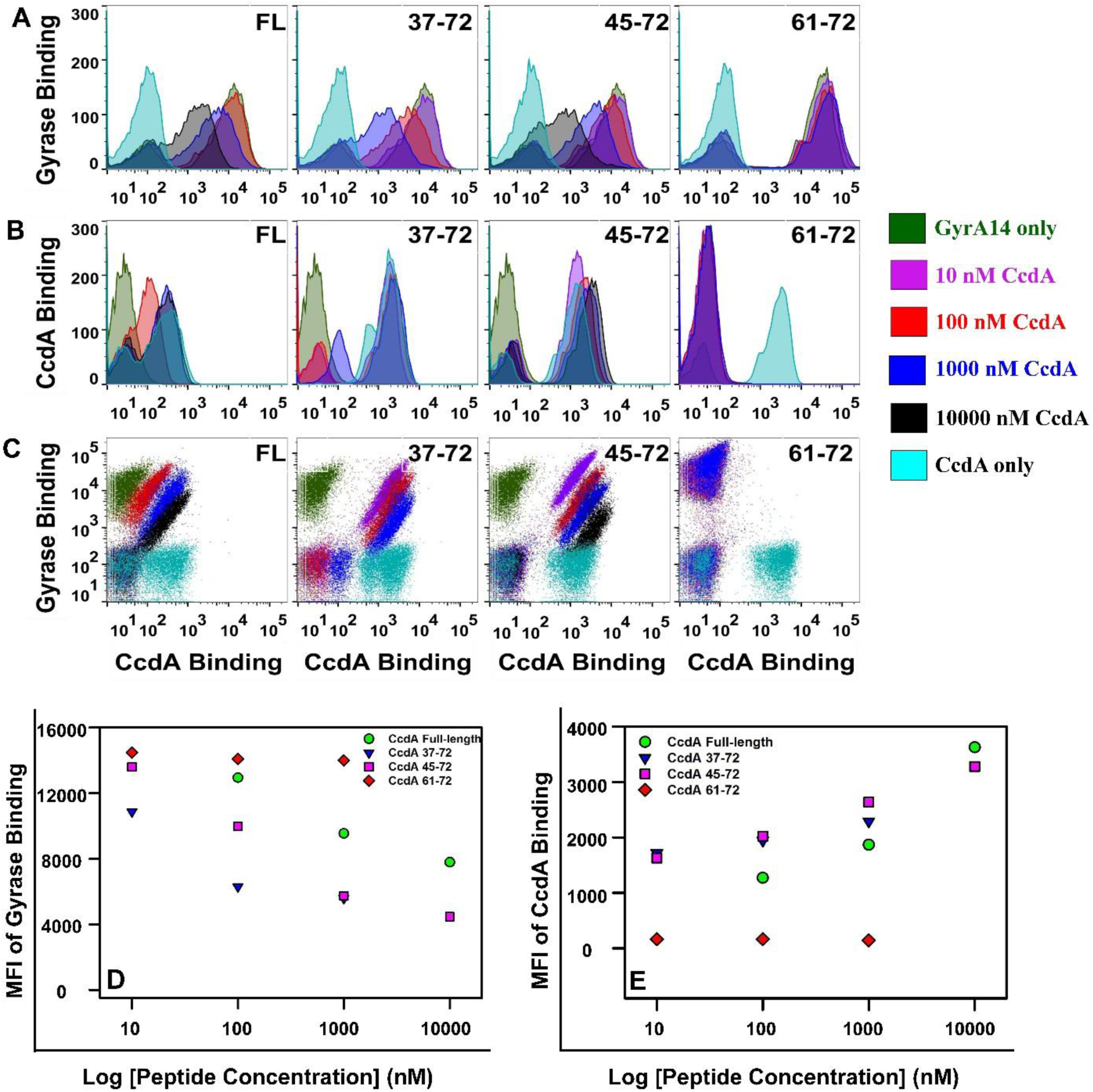
Effect of the length of CcdA on the rejuvenation process. Related to Figure 1 and 2. The rejuvenation process of the CcdB WT:GyrA14 complex on the yeast surface was monitored by using different concentrations of the full-length (FL) CcdA, CcdA_37-72_, CcdA_45-72_ and CcdA_61-72_. Comparison of the GyrA14 binding (A), CcdA binding (B) and double plots (C) showing both GyrA14 and CcdA binding of WT CcdB with different lengths of CcdA on the yeast cell surface. In all cases WT CcdB bound to 100 nM of GyrA14 is rejuvenated with different concentrations of CcdA derivative. The reaction is quenched by the addition of ten fold molar excess of purified WT CcdB over CcdA derivative after 30 s. Histograms showing rejuvenation mediated by 0 nM CcdA (green), 10 nM CcdA (purple), 100 nM (red), 1000 nM CcdA (blue), 10000 nM CcdA (black) are overlaid with the histogram showing binding with 1 nM CcdA in the absence of GyrA14 (cyan). Changes in the MFI values of GyrA14 binding (D) and MFI values of CcdA binding (E) by full-length CcdA and CcdA peptides of different lengths at 30 seconds plotted against varying CcdA concentrations are shown. Owing to the degradation prone nature of the antitoxins, the full-length CcdA shows a lower CcdA binding signal compared to the CcdA peptides. CcdA_61-72_ fails to rejuvenate the CcdB:GyrA14 complex. The effects on rejuvenation of CcdB:GyrA14 by full-length CcdA as well as CcdA_37-72_ and CcdA_45-72_ peptides are similar.

**Figure S2.**
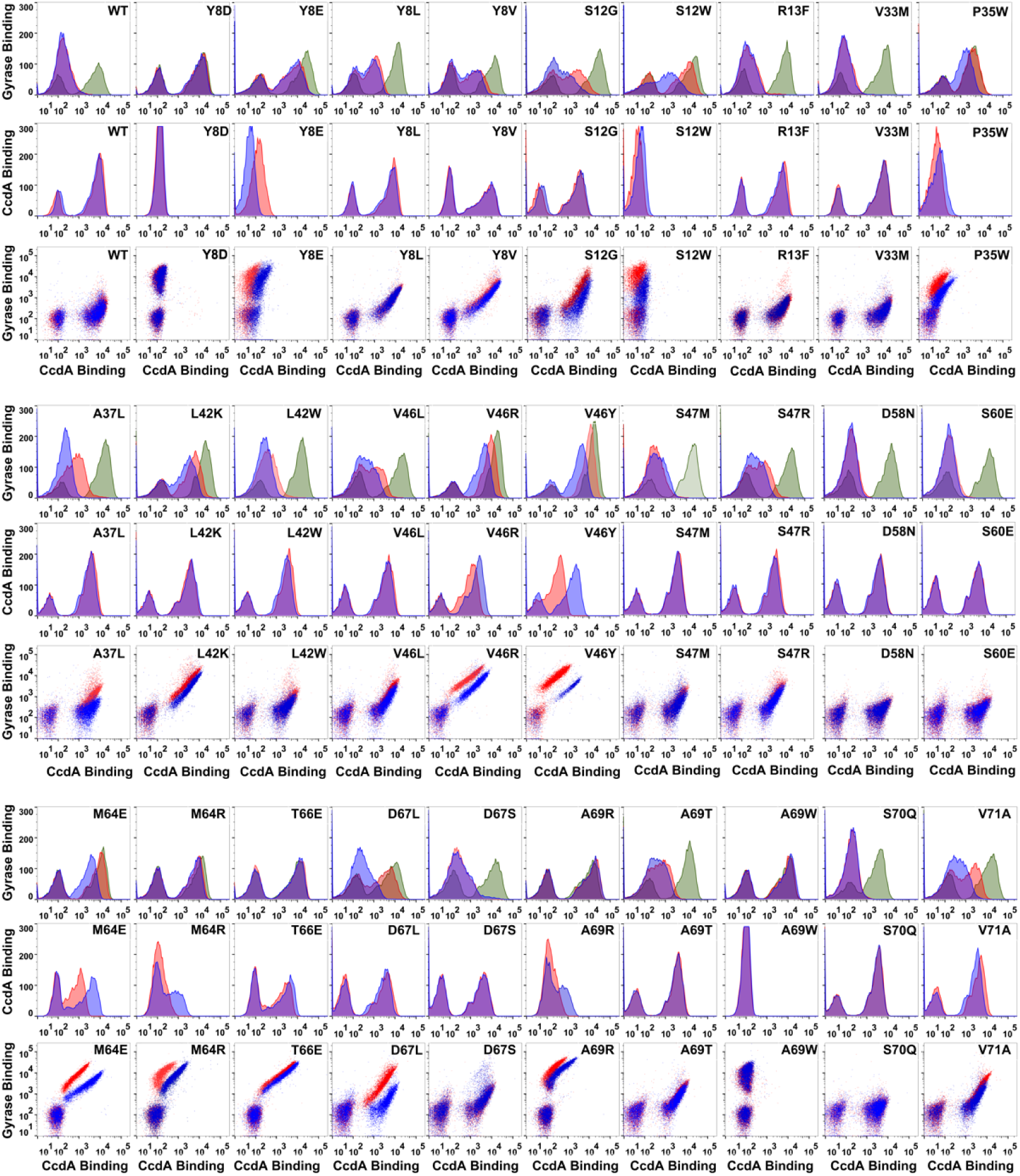
FACS analysis of rejuvenation of different CcdB mutants with slower rejuvenation rate constants than WT CcdB on the yeast cell surface. Related to Figure 2. 1D histograms and double plots showing both GyrA14 and CcdA_45-72_ binding of WT and different CcdB mutants displayed on the yeast cell surface. The GyrA14 binding histogram in the absence of CcdA_45-72_ is shown in green. Histogram for rejuvenation experiments with 100 nM CcdA_45-72_ for 30 s (red) is overlaid with histograms of rejuvenation experiments carried out with 1000 nM CcdA_45-72_ for 30 s (blue). The CcdB mutants, Y8D, S12W, P35W, M64R, A69R and A69W, have the strongest effect on the rejuvenation rate constants. All the mutations identified are located at residues involved in CcdA-binding, except for S60E.

**Figure S3.**
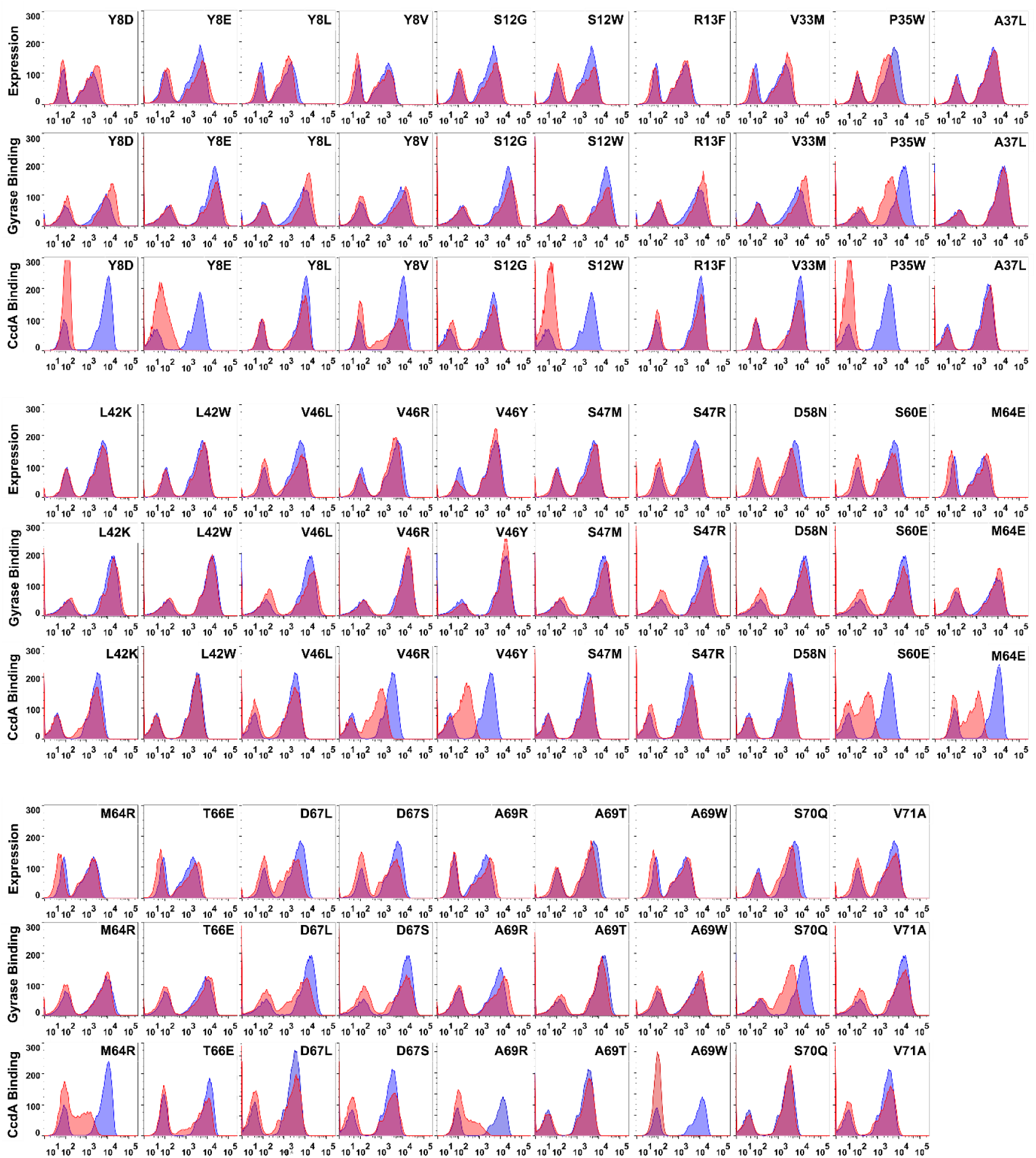
Analysis of yeast cell surface expression, GyrA14 (100 nM) and CcdA_45-72_ (100 nM) binding of different CcdB mutants with slower rejuvenation rate constants than WT. Related to Figure 2. Expression panels represent expression in the absence of GyrA14 and CcdA_45-72_. GyrA14 binding represents GyrA14 binding in the absence of CcdA_45-72_, and the CcdA_45-72_ binding panels represent CcdA_45-72_ binding in the absence of GyrA14. WT CcdB histogram (blue) is overlaid with the histograms obtained for the mutants (red) in each of the plots. A difference in the WT signal in different plots is observed as the measurements were carried out at different times and using two different instruments.

**Figure S4.**
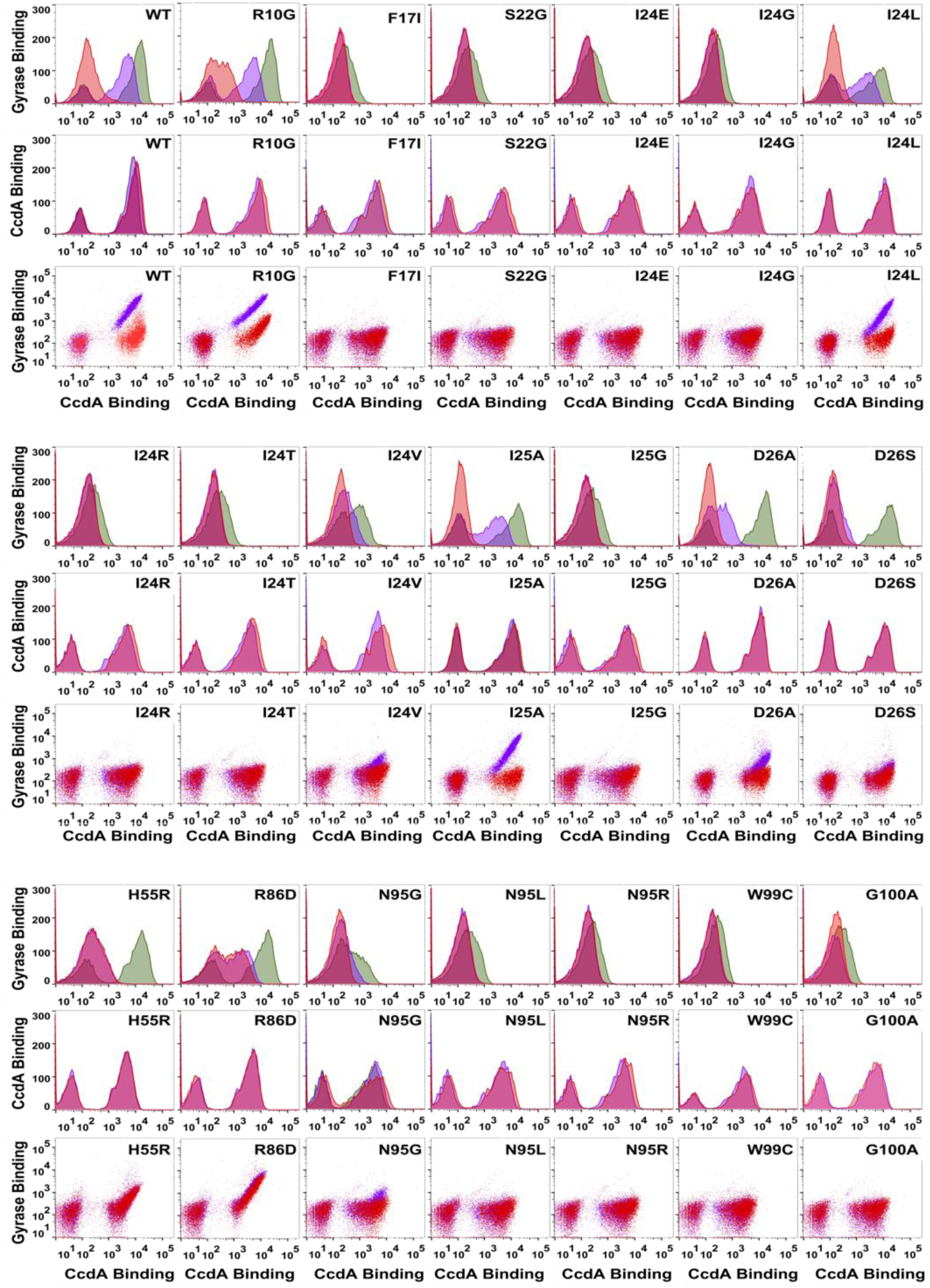
FACS analysis of rejuvenation of different CcdB mutants with faster rejuvenation rate constants than WT CcdB on the yeast cell surface. Related to Figure 2. The GyrA14 binding histogram in the absence of CcdA_45-72_ is shown in green. Histograms showing rejuvenation carried out with 10 nM CcdA_45-72_ for 5 s (purple) are overlaid with histograms showing rejuvenation carried out with 100 nM CcdA_45-72_ for 5 s (red). The CcdB mutants, namely, F17I, S22G, I24E, I24G, I24R, I24T, I25G, N95L, N95R, W99C and G100A have the strongest effects on the rejuvenation rate constants. All the mutants identified are located at residues involved in Gyrase-binding, except for F17I.

**Figure S5.**
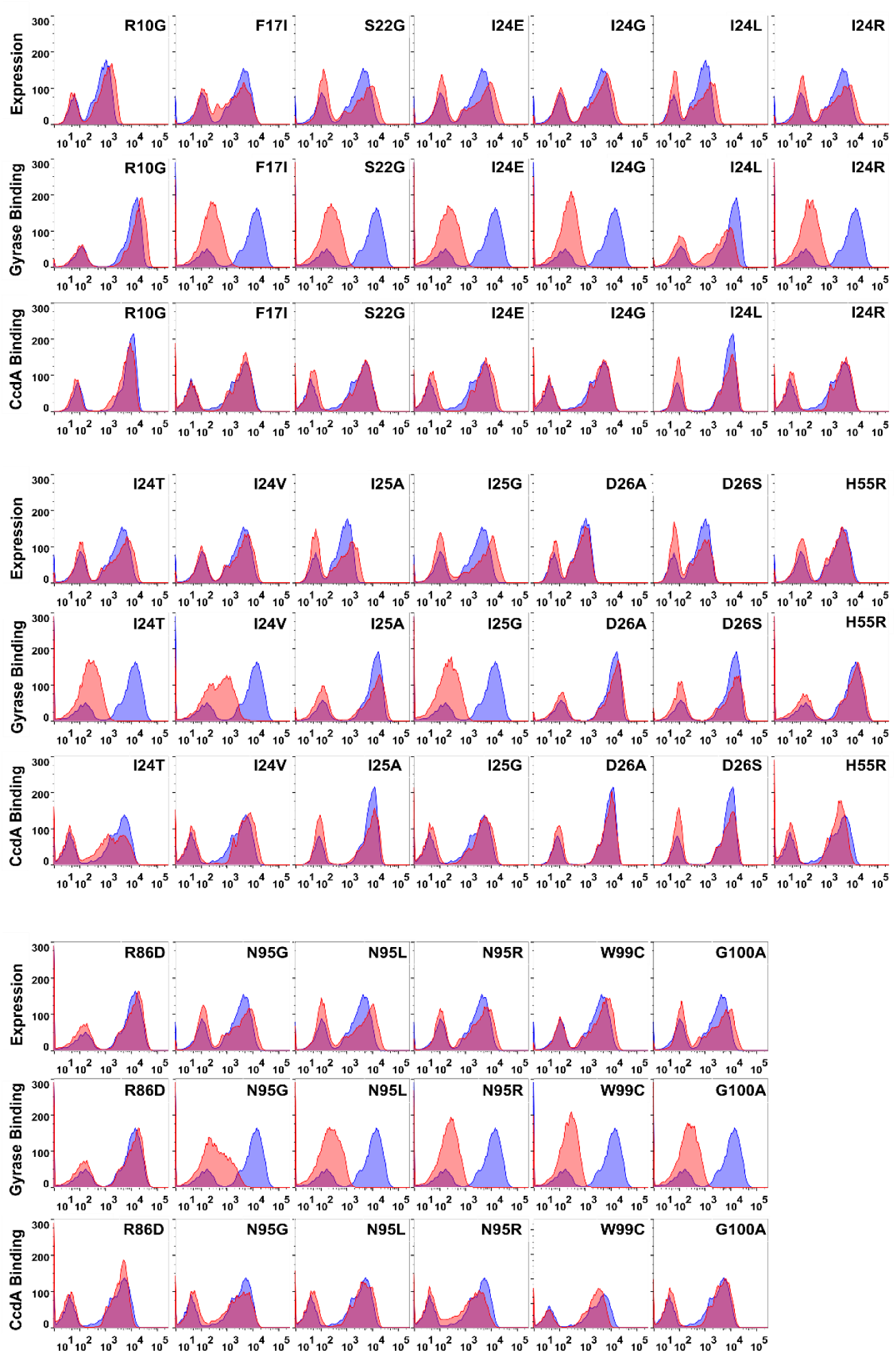
Analysis of yeast cell surface expression, GyrA14 (100 nM) and CcdA_45-72_ (100 nM) binding of different CcdB mutants with faster rejuvenation rate constants than WT. Related to Figure 2. Expression panels represent expression in the absence of GyrA14 and CcdA_45-72_. GyrA14 binding represents GyrA14 binding in the absence of CcdA_45-72_ and the CcdA_45-72_ binding panels represent CcdA_45-72_ binding in the absence of GyrA14. WT CcdB histogram (blue) is overlaid with histograms observed for the mutants (red) in each of the plots. A difference in the WT signal in different plots is observed as the measurements were carried out at different times and using two different instruments.

**Figure S6.**
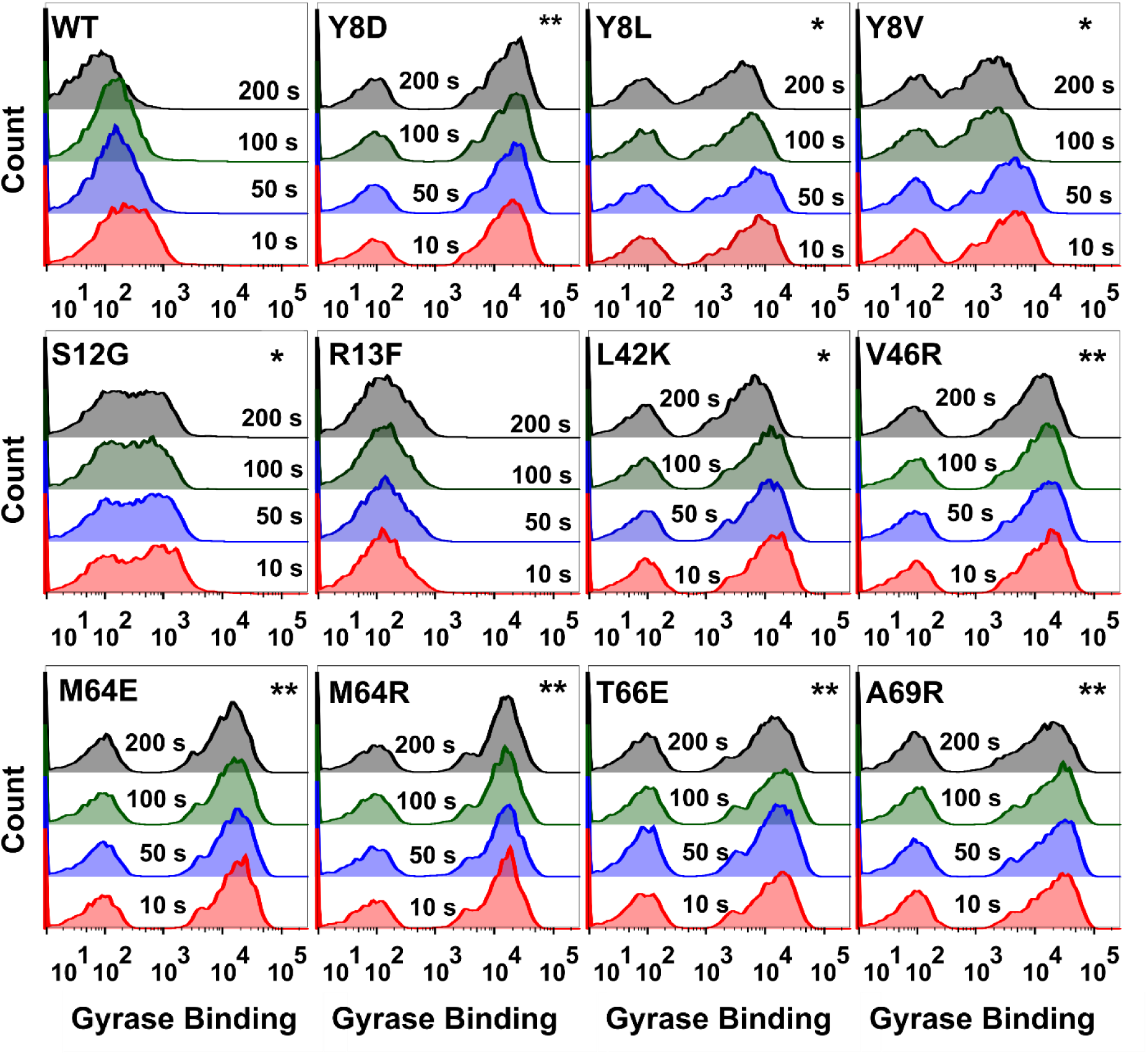
Analysis of different CcdB mutants with slower rejuvenation rate constants than the WT CcdB on the yeast cell surface. Related to Figure 2. Comparison of the binding of WT or different CcdB mutants on the yeast cell surface to GyrA14 are shown. Histograms of GyrA14 binding signals after rejuvenation carried out with 100 nM CcdA_45-72_ for 10 s (red) are overlaid with histograms showing GyrA14 binding signals after rejuvenation carried out with 100 nM CcdA_45-72_ for 50 s (blue), 100 s (green) and 200 s (black). Several CcdB mutants, namely, Y8D, Y8L, Y8V, L42K, V46R, M64E, M64R, T66E, and A69R showed significant GyrA14 binding even after 200 s. While S12G showed a reduction in GyrA14 binding, the rate was still significantly slower than WT. R13F behaved similar to WT and was used as a control.* indicates mutants with decreased CcdA_45-72_ binding and ** with highly decreased or no CcdA_45-72_ binding as in Figure S3.

**Figure S7.**
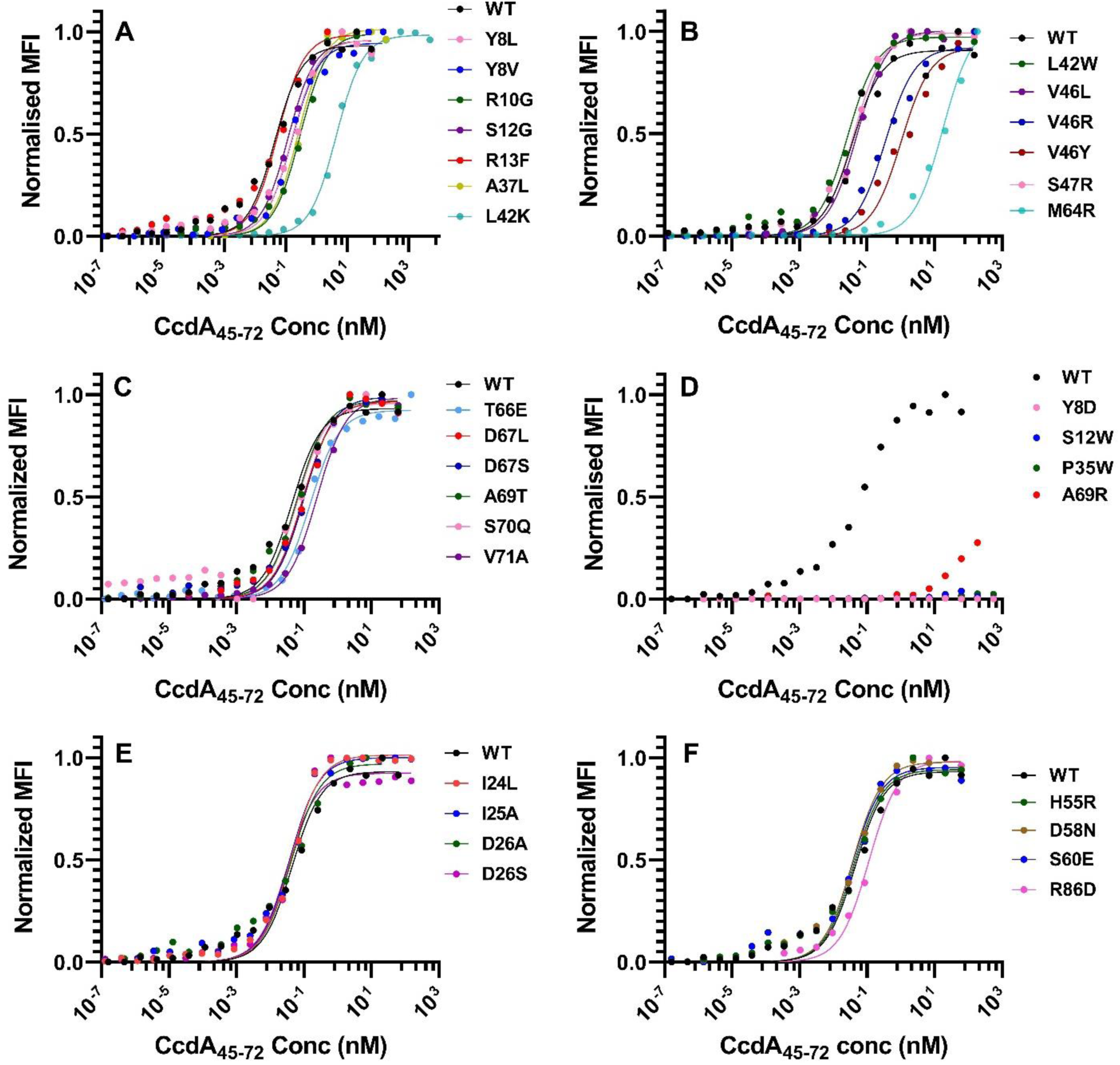
Binding of CcdA_45-72_ peptide to CcdB mutants measured by YSD. Related to Figure 3. (A-C) Binding to CcdA_45-72_ peptide of CcdB mutants at the CcdA binding interface. (D) CcdB mutants at the CcdA binding interface which bind poorly to CcdA. (E) Binding to CcdA_45-72_ peptide of CcdB mutants at the GyrA14 binding interface. (F) Binding to CcdA_45-72_ peptide of CcdB mutants which are not involved in binding with either CcdA or GyrA14. Mutants were displayed on the yeast cell surface and titrated with serial dilutions of the CcdA peptide 45-72. The binding was measured on a BD Aria III flow cytometer and the data was fit using GraphPad Prism 8.0 (also listed in Table S2).

**Figure S8.**
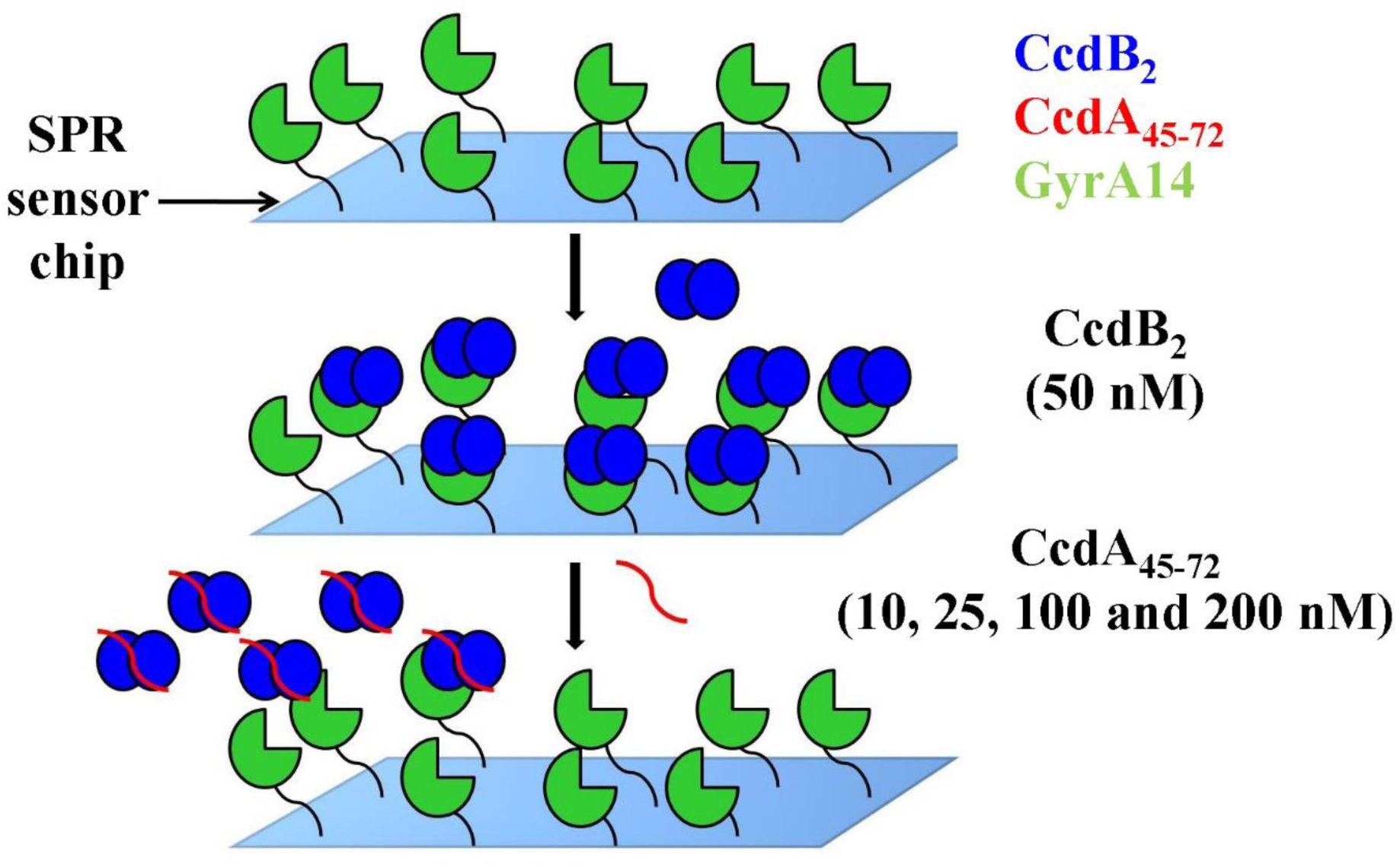
Schematic of Surface Plasmon Resonance (SPR) experiment used to measure rejuvenation rate constant. Related to Figure 4. (A) GyrA14 (green) is immobilized on a CM5 chip (cyan) by amine coupling. 50 nM of the CcdB mutants (blue) are passed over the immobilized GyrA14 for 100 s followed by dissociation mediated by different concentrations of the CcdA_45-72_ peptide (red).

**Figure S9.**
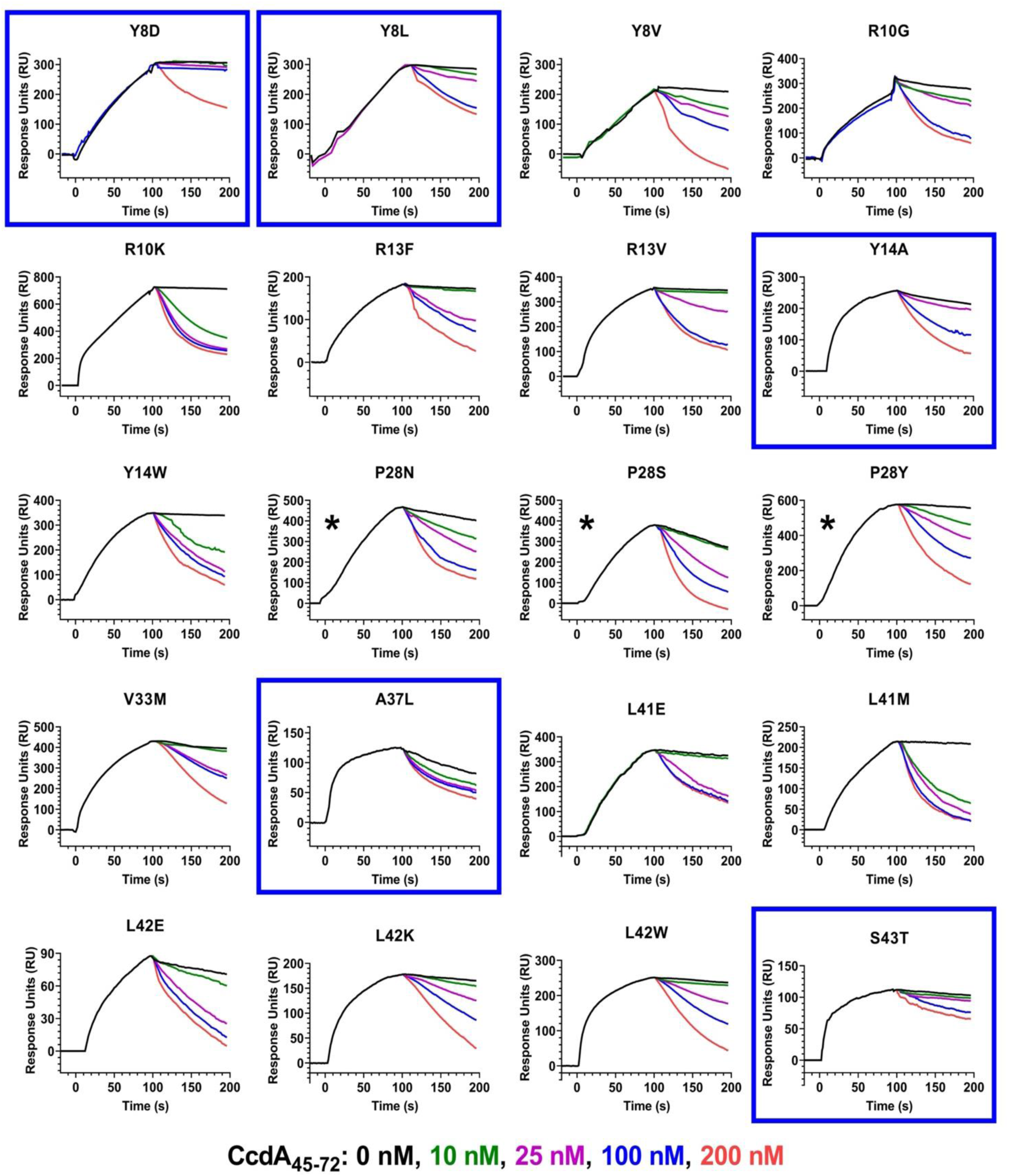
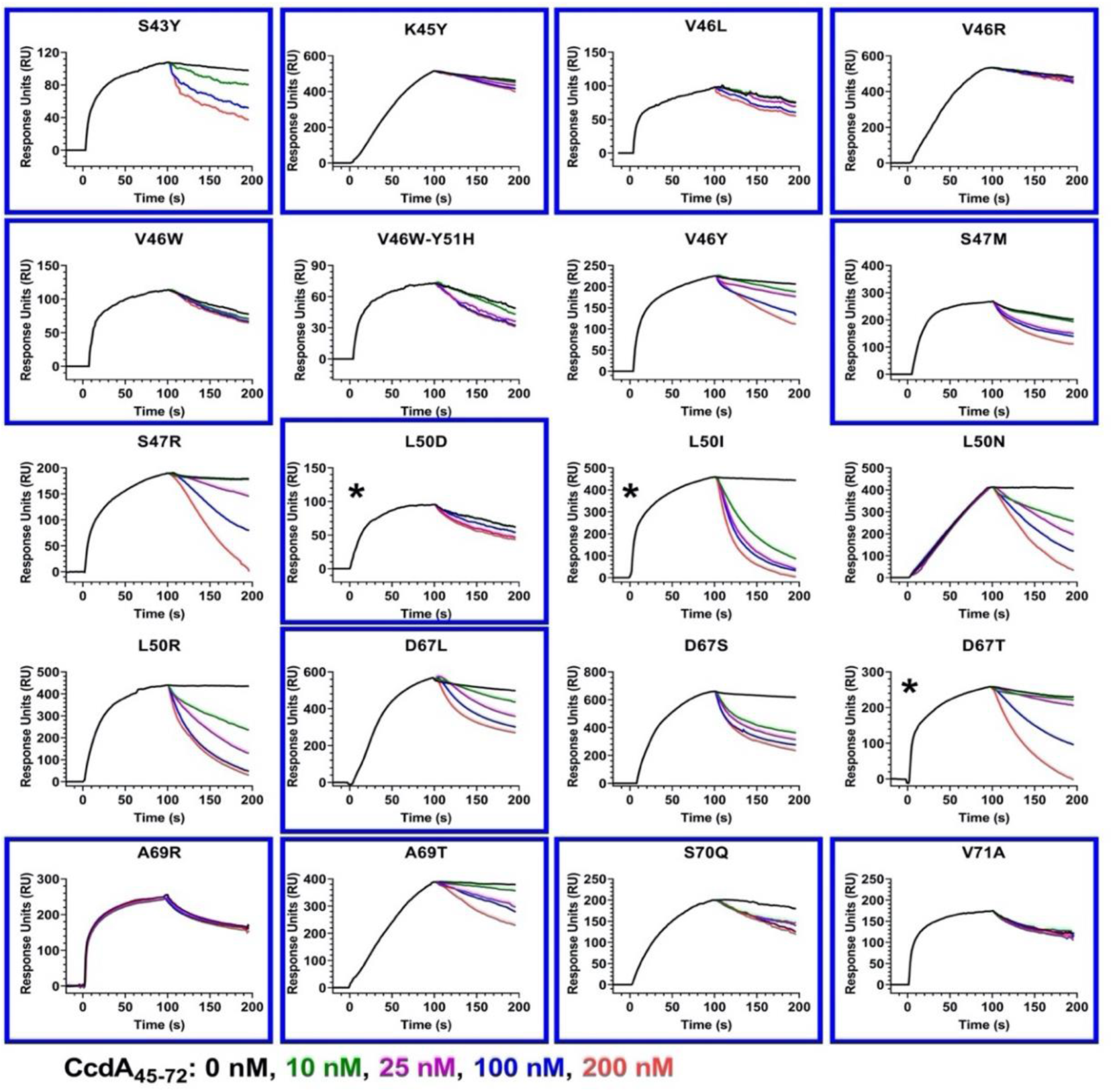
Dissociation of CcdA interacting-site CcdB mutants from GyrA14, in the presence and absence of CcdA_45-72_. Related to Figure 4 and 5. Overlays show the binding kinetics of 50 nM of these CcdB mutants which are passed over GyrA14 immobilized on a CM5 chip for 100 s, followed by dissociation mediated by different concentrations of CcdA_45-72_. CcdA_45-72_ peptide concentrations increase from top to bottom, in the order 0 nM (black), 10 nM (green), 25 nM (purple), 100 nM (blue) and 200 nM (red). The ligand GyrA14 was immobilized on the CM5 chip by standard amine coupling. The rejuvenation of WT CcdB is reported previously (Aghera et al., 2020). Mutants with significantly slower rejuvenation than WT are highlighted in blue boxes and mutants with altered binding to GyrA14 are marked with a star (*). All studies are carried out in 1XPBS running buffer, pH7.4 at 25 °C . The rejuvenation of WT CcdB is similar to I24L data shown in (Figure S10) and was carried out previously (Aghera et al., 2020).

**Figure S10.**
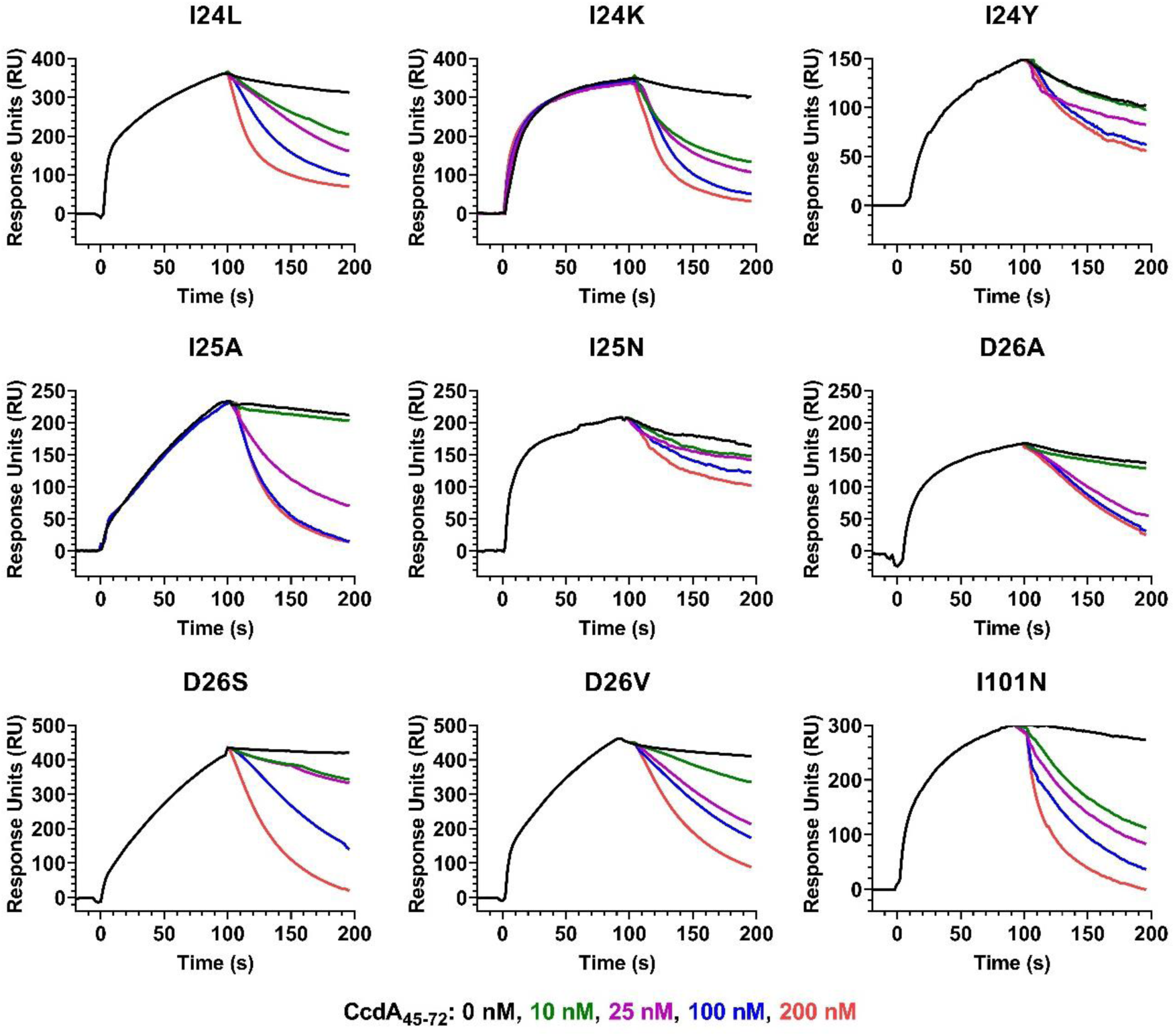
Dissociation of GyrA-14 interacting-site CcdB mutants from GyrA14, in the presence and absence of CcdA_45-72_. Related to Figure 4 and 5. Overlays show the binding kinetics of 50 nM of these CcdB mutants which are passed over GyrA14 immobilized on a CM5 chip for 100 s, followed by dissociation mediated by different concentrations of CcdA_45-72_. CcdA_45-72_ peptide concentrations increase from top to bottom, in the order 0 nM (black), 10 nM (green), 25 nM (purple), 100 nM (blue) and 200 nM (red). The ligand GyrA14 was immobilized on the CM5 chip by standard amine coupling. All studies are carried out in 1XPBS running buffer, pH7.4 at 25 °C. The rejuvenation of WT CcdB is similar to I24L data and was carried out previously (Aghera et al., 2020).

**Figure S11.**
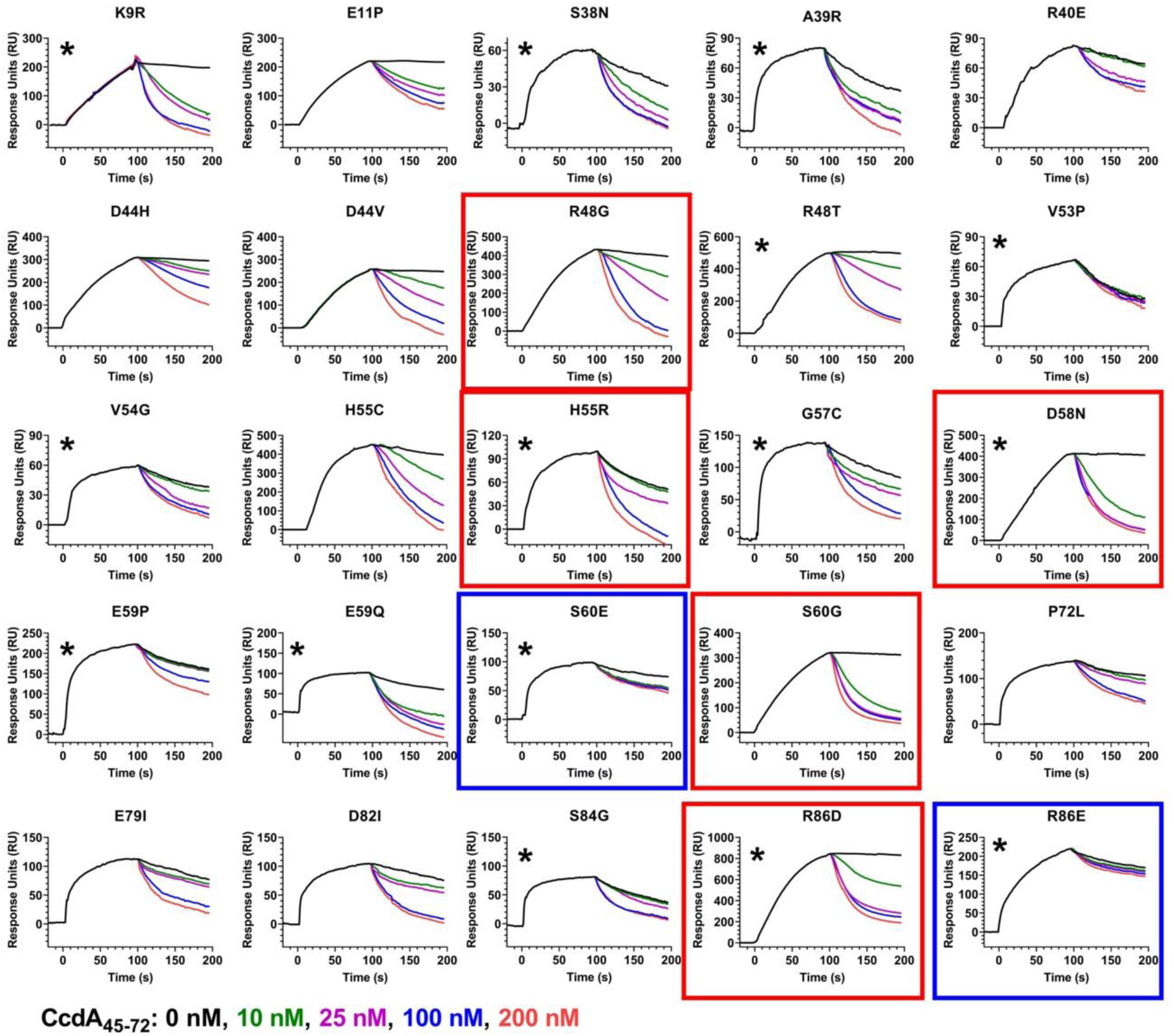
Dissociation of CcdB mutants at residues outside the GyrA14 and CcdA binding site from GyrA14 in the presence and absence of CcdA_45-72_. Related to Figure 4 and 5. Overlays show the binding kinetics of 50 nM of these CcdB mutants which are passed over GyrA14 immobilized on a CM5 chip for 100 s, followed by dissociation mediated by different concentrations of CcdA_45-72_. CcdA_45-72_ peptide concentrations increase from top to bottom, in the order 0 nM (black), 10 nM (green), 25 nM (purple), 100 nM (blue) and 200 nM (red). The ligand GyrA14 was immobilized on the CM5 chip by standard amine coupling. Mutants with significantly slower and faster rejuvenation than WT are highlighted in blue and red boxes respectively and mutants with altered binding to GyrA14 are marked with a star (*). All studies are carried out in 1XPBS running buffer, pH7.4 at 25 °C. The rejuvenation of WT CcdB is similar to I24L data shown in (Figure S10) and was carried out previously (Aghera et al., 2020).

**Figure S12.**
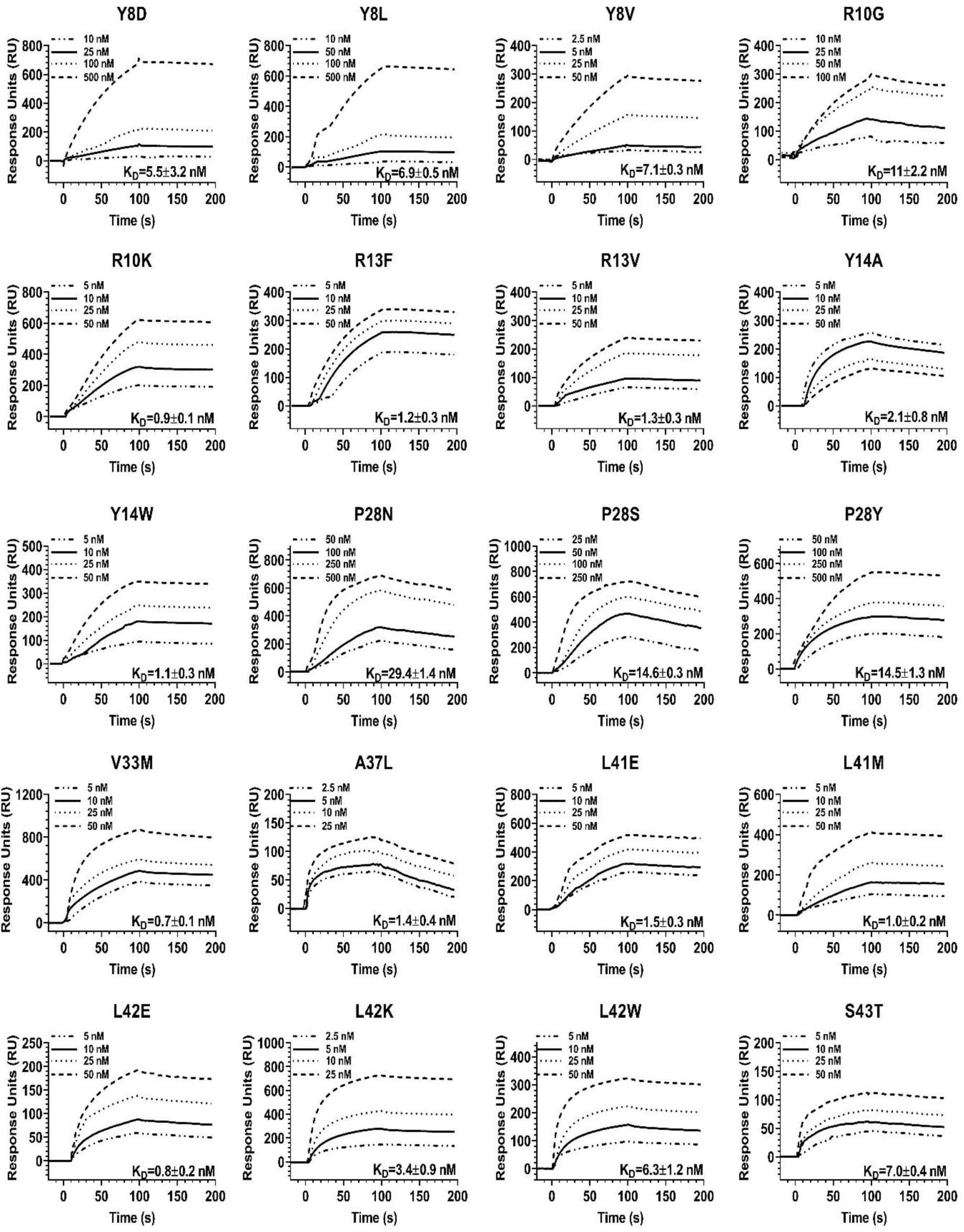
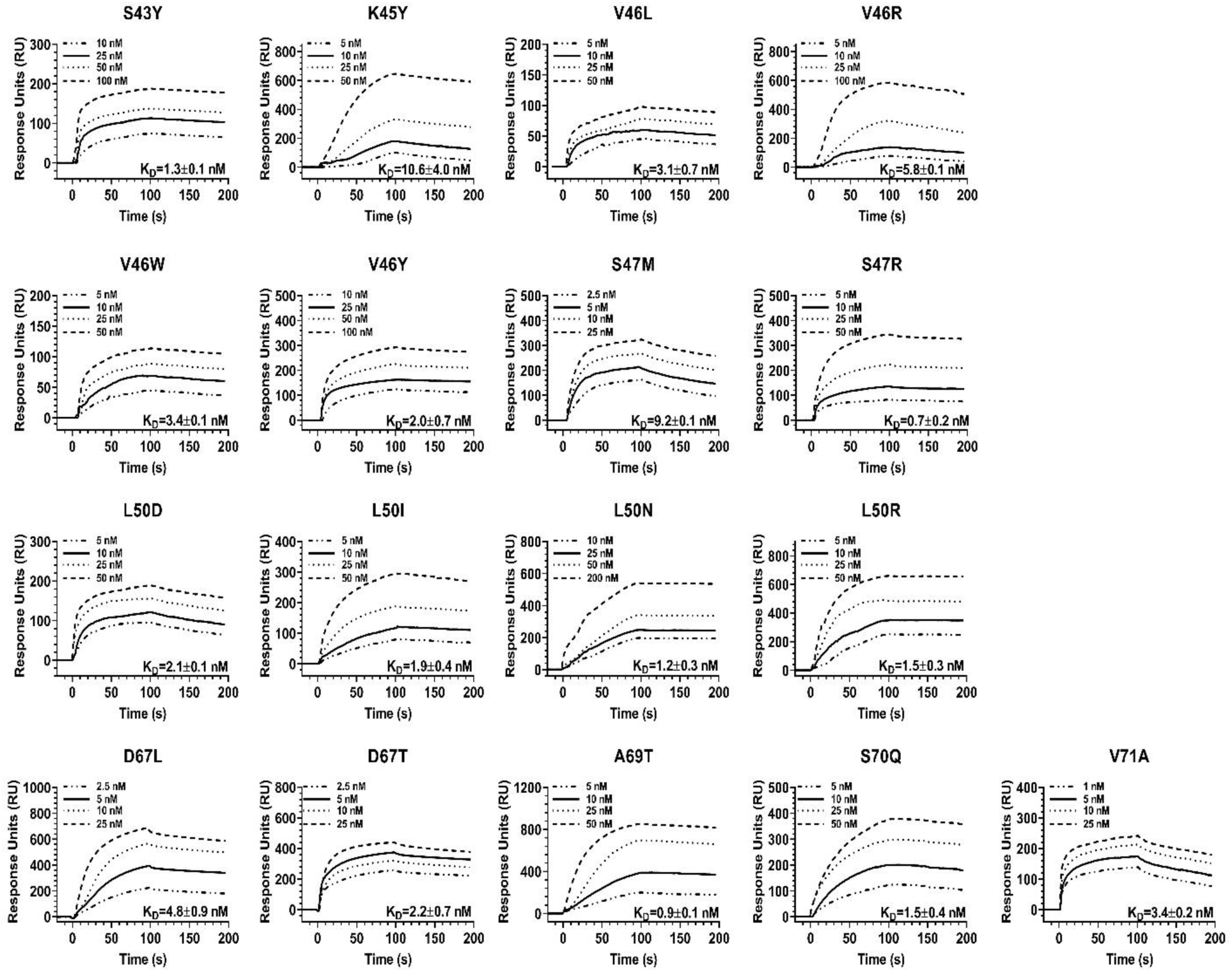
Binding of GyrA14 to CcdA interacting-site CcdB mutants. Related to Figure 4 and 5. Overlays show the binding kinetics with analyte concentrations increasing from bottom to top inside each plot for Y8D (10, 25, 100, 500 nM), Y8L (10, 50, 100, 500 nM), Y8V (2.5, 5, 25, 50 nM), R10G (10, 25, 50, 100 nM), R10K (5, 10, 25, 50 nM), R13F (5, 10, 25, 50 nM), (I) R13V (5, 10, 25, 50 nM), Y14A (10, 25, 50, 100 nM), Y14W (5, 10, 25, 50 nM), P28N (50, 100, 250, 500 nM), P28S (25, 50, 100, 250 nM), P28Y (50, 100, 250, 500 nM), V33M (5, 10, 25, 50 nM), A37L (2.5, 5, 10, 25 nM), L41E (5, 10, 25, 50 nM), L41M (5, 10, 25, 50 nM), L42E (5, 10, 25, 50 nM), L42K (2.5, 5, 10, 25 nM), L42W (5, 10, 25, 50 nM), S43T (5, 10, 25, 50 nM), S43Y (10, 25, 50, 100 nM), K45Y (5, 10, 25, 50 nM), V46L (5, 10, 25, 50 nM), V46R (5, 10, 25, 50 nM), V46W (5, 10, 25, 50 nM), V46Y (10, 25, 50, 100 nM), S47M (2.5, 5, 10, 25 nM), S47R (5, 10, 25, 50 nM), L50D (5, 10, 25, 50 nM), L50I (5, 10, 25, 50 nM), L50N (10, 25, 50, 200 nM), L50R (5, 10, 25, 50 nM), D67L (2.5, 5, 10, 25 nM), D69T (2.5, 5, 10, 25 nM), A69T (5, 10, 25, 50 nM), S70Q (5, 10, 25, 50 nM) and V71A (1, 5, 10, 25 nM). The ligand GyrA14 was immobilized on the CM5 chip by standard amine coupling. Binding was measured by passing varying concentrations of the analyte (CcdB proteins) over the ligand (GyrA14) immobilised chip and the data was fitted to a 1:1 Langmuir interaction model to obtain the kinetic parameters (also listed in Table S7). All studies are carried out in 1XPBS running buffer, pH7.4 at 25 °C. WT CcdB binds GyrA14 with a K_D_ of 1.4 nM (Chattopadhyay et al., 2021).

**Figure S13.**
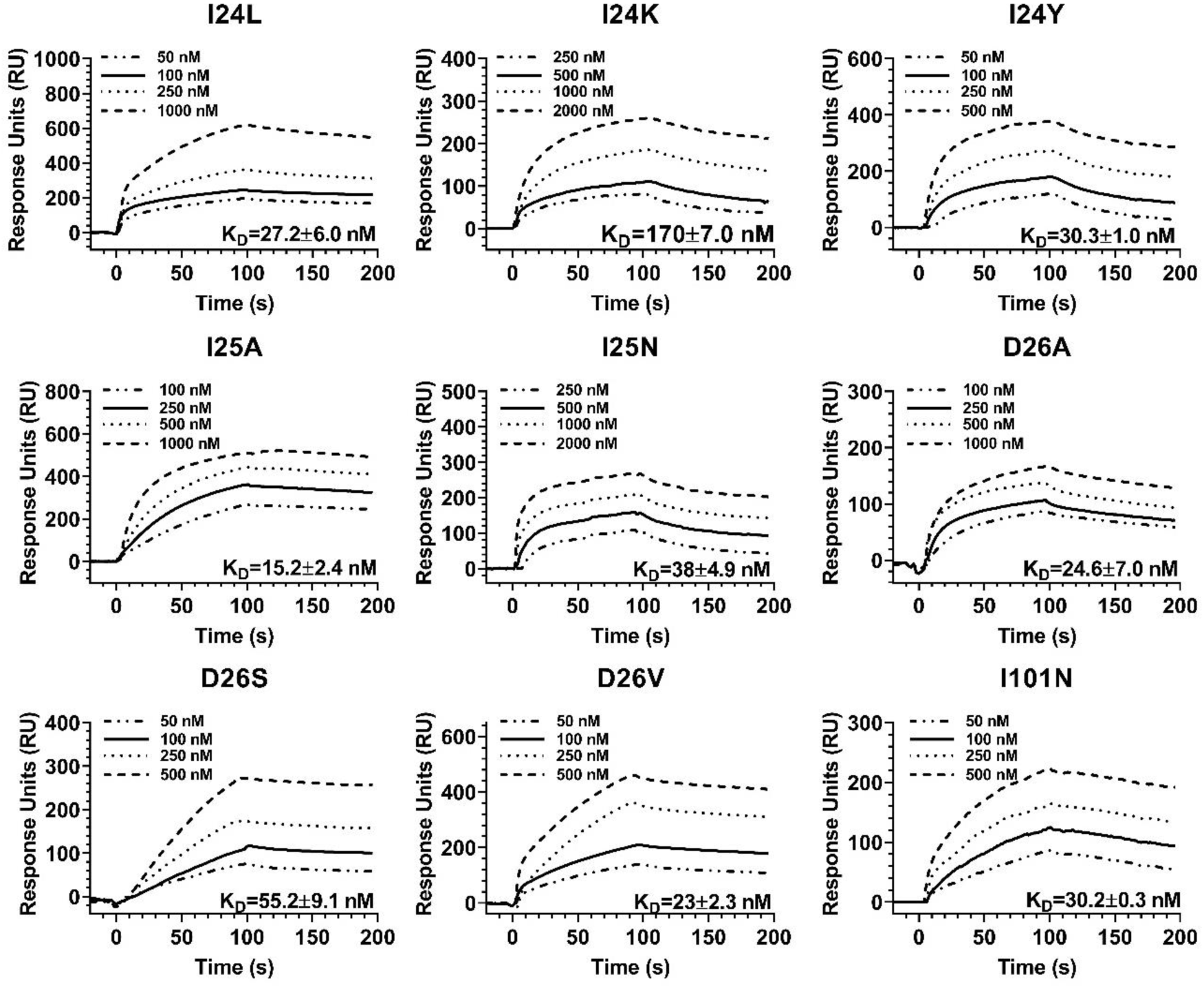
Binding of GyrA14 to GyrA14 interacting-site CcdB mutants. Related to Figure 4 and 5. Overlays show the binding kinetics with analyte concentrations increasing from bottom to top inside each plot for I24L (50, 100, 250, 1000 nM), I24K (250, 500, 1000, 2000 nM), I24Y (50, 100, 250, 500 nM), I25A (100, 250, 500, 1000 nM), I25N (250, 500, 1000, 2000 nM), D26A (100, 250, 500, 1000 nM), D26S (50, 100, 250, 500 nM), D26V (50, 100, 250, 500 nM) and I101N (50, 100, 250, 500 nM). The ligand GyrA14 was immobilized on the CM5 chip by standard amine coupling. Binding was measured by passing varying concentrations of the analyte (CcdB proteins) over the ligand (GyrA14) immobilised chip and the data was fitted to a 1:1 Langmuir interaction model to obtain the kinetic parameters (also listed in Table S8). All studies are carried out in 1XPBS running buffer, pH7.4 at 25 °C. WT CcdB binds GyrA14 with a KD of 1.4 nM (Chattopadhyay et al., 2021).

**Figure S14.**
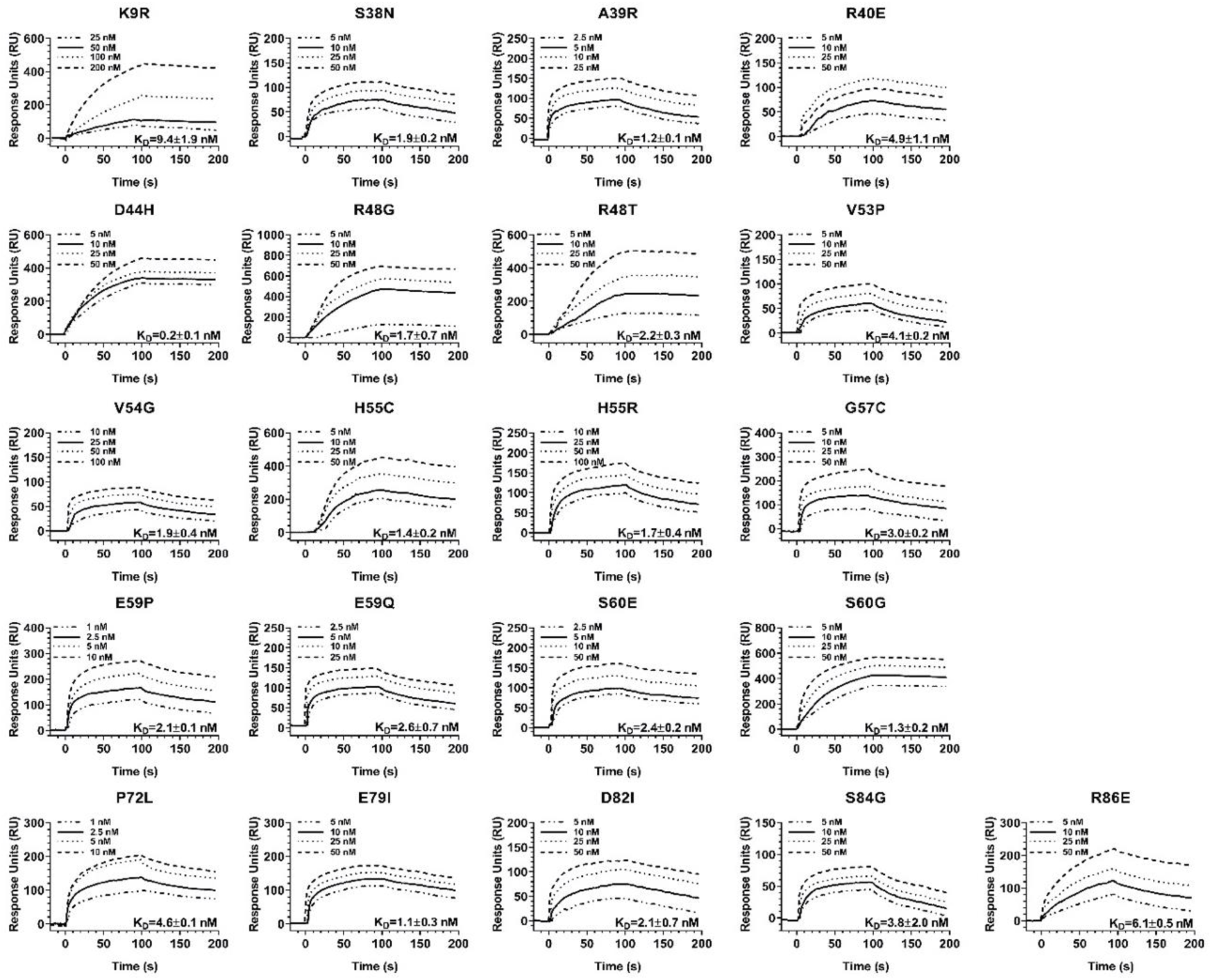
Binding of GyrA14 to CcdB mutants at sites that neither interact with CcdA or GyrA14. Related to Figure 4 and 5. Overlays show the binding kinetics with analyte concentrations increasing from bottom to top inside each plot for K9R (25, 50, 100, 200 nM), S38N (5, 10, 25, 50 nM), A39R (2.5, 5, 10, 25 nM), R40E (5, 10, 25, 50 nM), D44H (5, 10, 25, 50 nM), R48G (5, 10, 25, 50 nM), R48T (5, 10, 25, 50 nM), V53P (5, 10, 25, 50 nM), V54G (10, 25, 50, 100 nM), H55C (5, 10, 25, 50 nM), H55R (10, 25, 50, 100 nM), G57C (5, 10, 25, 50 nM), E59P (1, 2.5, 5, 10 nM), E59Q (2.5, 5, 10, 25 nM), S60E (2.5, 5, 10, 50 nM), S60G (5, 10, 25, 50 nM), P72L (1, 2.5, 5, 10 nM), E79I (5, 10, 25, 50 nM), D82I (5, 10, 25, 50 nM), S84G (5, 10, 25, 50 nM) and R86E (1, 2.5, 5, 10 nM). The ligand GyrA14 was immobilized on the CM5 chip by standard amine coupling. Binding was measured by passing varying concentrations of the analyte (CcdB proteins) over the ligand (GyrA14) immobilised chip and the data was fitted to a 1:1 Langmuir interaction model to obtain the kinetic parameters (also listed in Table S9). All studies are carried out in 1XPBS running buffer, pH7.4 at 25 °C. WT CcdB binds GyrA14 with a KD of 1.4 nM (Chattopadhyay et al., 2021).

**Figure S15.**
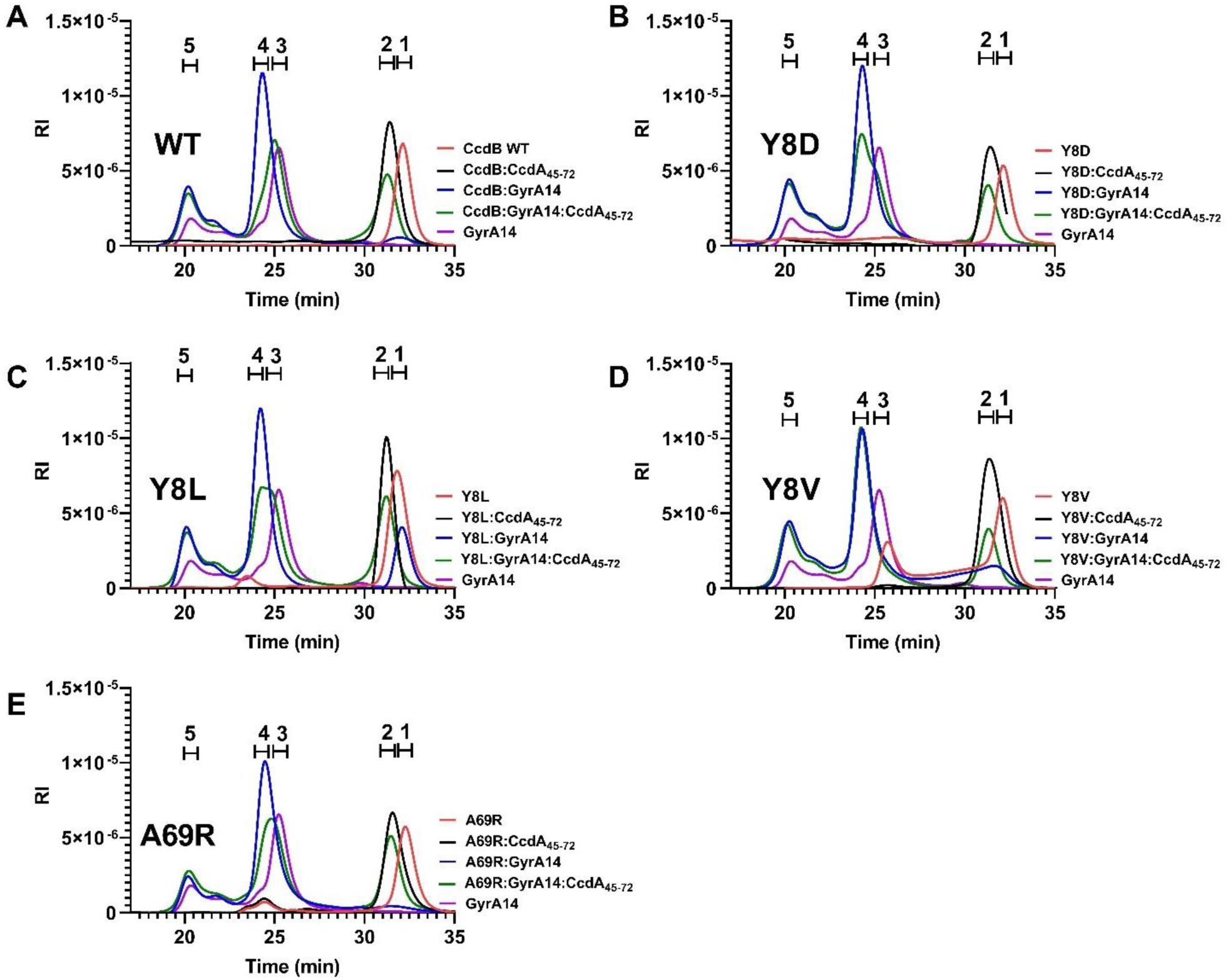
Analysis of purified WT and mutant CcdB proteins and complexes by SEC MALS. Related to Figure 6. Traces for CcdB-WT and mutants (red), GyrA14 (magenta), WT CcdB:CcdA_45-72_ complex (black), WT CcdB:GyrA14 complex (blue) and CcdA_45-72_:CcdB WT:GyrA14 complex (green) are coloured differently. The normalised refractive indices (RI) of all the traces are plotted as a function of time. The complexes are formed *in vitro* by mixing 10 µM of each of the proteins and incubating them for 30 minutes before injection. For the rejuvenation experiments, the CcdB:GyrA14 was incubated for 30 minutes, after which CcdA_45-72_ peptide was added and the sample was immediately injected. All studies are carried out in 1XPBS, pH7.4 at 25 °C. The molar mass and mass fraction of each of the peaks are listed in Table S10.

**Figure S16.**
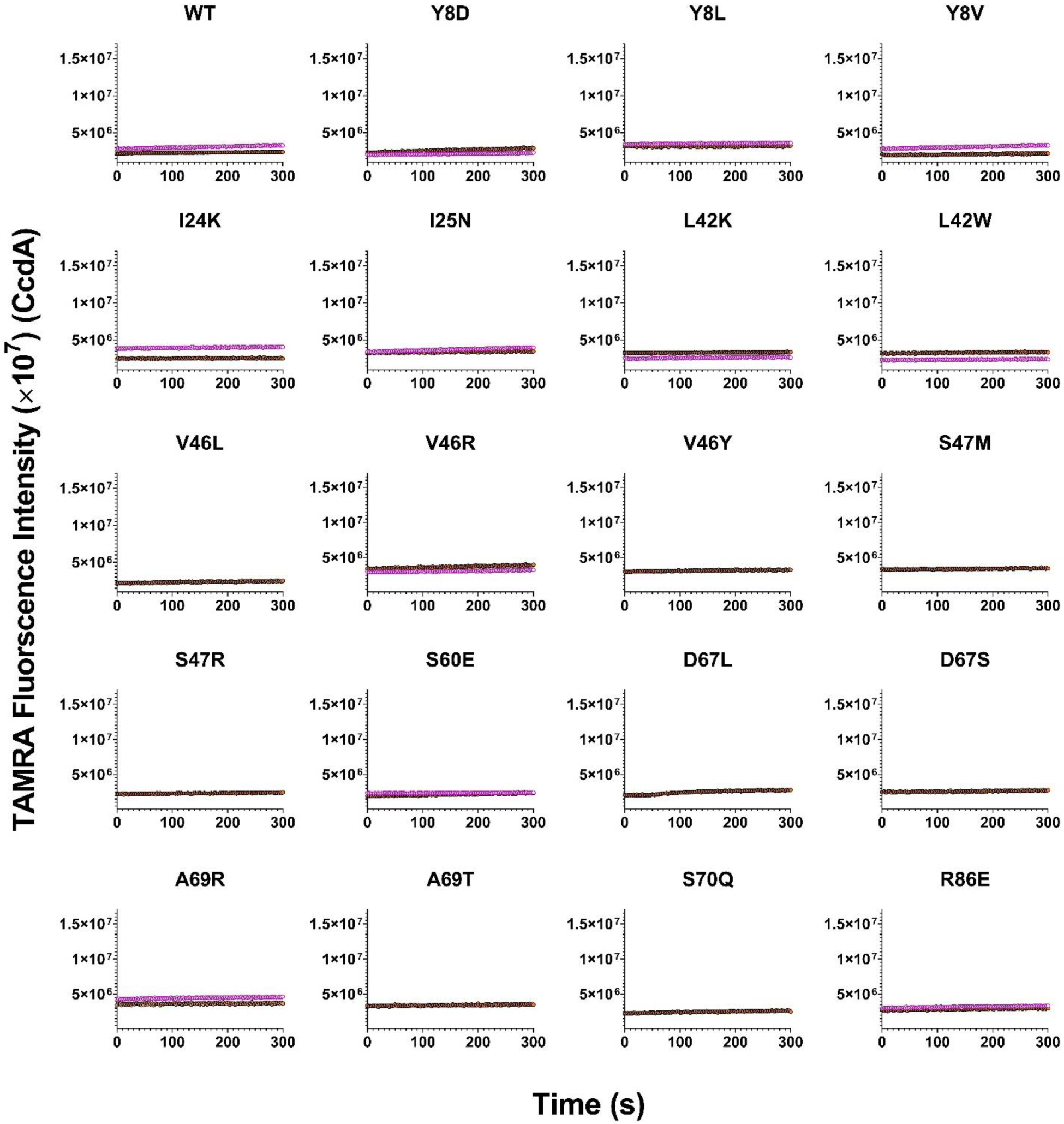
Rejuvenation studies using unlabeled CcdA_50-72_ peptide. Related to Figure 6. Overlays show the lack of time-dependent change in fluorescence upon mixing of 10 nM of a preformed complex CcdB:GyrA14* with 100 nM of unlabeled CcdA_50-72_ (magenta open circles). The fluorescence traces of 10 nM of CcdB:GyrA14* are shown in orange solid circles.

## Supplementary Tables

**Table S1.** Related to Figure 1, 2, 6 and S1. Details of the various CcdA peptides used in the study.

| **CcdA** | **Sequence** | **Length** | **Molecular weight (Da)** |
| --- | --- | --- | --- |
| **Full-length** | MKQRITVTVDSDSYQLLKAYDVNIS<br>GLVSTTMQNEARRLRAERWKAENQ<br>EGMAEVARFIEMNGSFADENRDW | 72 | 8372 |
| **37-72** | * **CRRLRAERWKAENQ** EGMAEVARF<br>IEMNGSFADENRDW | 37 | 4473 |
| **45-72** | * **KGSVENQEGMVEVARFIEMNGSFA**<br>DENRDW | 30 | 3445 |
| **50-72** | EGMAEV**A**C#FIEMNGSFADENRDW | 23 | 2621 |
| **61-72** | * **KGSMNGSFADENRDW** | 15 | 1714 |

\*: In bold are the additional N-terminal Cysteine and Lysine residues present in the CcdA peptides synthesised from GenScript to facilitate conjugation to an appropriate matrix.
#: In bold is the additional Cysteine residue present in the CcdA<sub>50-72</sub> peptide synthesised from GenScript to facilitate FRET studies.

**Table S2.** Related to Figure 3, 4 and 6. Comparison of apparent dissociation rate constants (k_d_) of different CcdB mutants from GyrA14 with 10 (*) and 100 nM of CcdA_45-72_ peptide by YSD,SPR and FRET, and the CcdA-binding affinities of CcdB mutants measured by YSD^1^.

| Mutants | $k_d$ (s<sup>-1</sup>) | | | $K_D$ (pM) |
| --- | --- | --- | --- | --- |
|  | YSD | SPR | FRET |  |
| WT | 35.0e-3 | 47.2e-3 | 79.1e-3 | 44 |
| WT* | 8.6e-3 | 6.2e-3 | N.D. |  |
| Y8D | 2.4e-3 | 0.5e-3 | 5.2e-3 | N.B. |
| Y8L | 9.0e-3 | 8.2e-3 | 17e-3 | 160 |
| Y8V | 6.6e-3 | 8.6e-3 | 11e-3 | 158 |
| R10G* | 6.2e-3 | 2.3e-3 | N.D. | 290 |
| S12G | 6.7e-3 | 8.2e-3 | N.D. | 118 |
| S12W | N.D. | N.D. | N.D. | N.B. |
| R13F | 13.0e-3 | 9.7e-3 | N.D. | 54 |
| I24L* | 8.5e-3 | 12.0e-3 | N.D. | 40 |
| I24K | N.D. | 47.1e-3 | 88e-3 | N.D. |
| I25A* | 8.8e-3 | 13.1e-3 | N.D. | 37 |
| I25N | N.D. | 47.0e-3 | 82e-3 | N.D. |
| D26A* | 7.1e-3 | 9.0e-3 | N.D. | 40 |
| D26S* | 7.8e-3 | 13.7e-3 | N.D. | 32 |
| V33M | 5.9e-3 | 6.5e-3 | N.D. | 5 |
| P35W | N.D. | N.D. | N.D. | N.B. |
| A37L | 13.0e-3 | 8.0e-3 | N.D. | 52 |
| L42K | 6.0e-3 | 7.5e-3 | 9.2e-3 | 133 |
| L42W | 6.3e-3 | 5.6e-3 | 8.7e-3 | 27 |
| V46L | 9.6e-3 | 5.3e-3 | 25e-3 | 52 |
| V46R | 5.0e-3 | 2.5e-3 | 4.6e-3 | 348 |
| V46Y | 2.9e-3 | 4.4e-3 | 20e-3 | 1027 |
| S47M | 9.0e-3 | 5.3e-3 | 21e-3 | N.D. |
| S47R | 8.9e-3 | 8.0e-3 | 11e-3 | 40 |
| H55R* | 5.7e-3 | 6.2e-3 | N.D. | 40 |
| D58N | 37.0e-3 | 45.0e-3 | N.D. | 37 |
| S60E | 13.0e-3 | 4.7e-3 | 34e-3 | 36 |
| M64R | N.D. | N.D. | N.D. | 17670 |
| T66E | N.D. | N.D. | N.D. | 151 |
| D67L | 7.5e-3 | 7.2e-3 | 7.2e-3 | 90 |
| D67S | 11.0e-3 | 6.5e-3 | 32e-3 | 101 |
| A69R | N.D. | 2.7e-3 | 6e-3 | 39840 |
| A69T | 4.2e-3 | 3.4e-3 | 11e-3 | 61 |
| S70Q | 4.9e-3 | 4.9e-3 | 15e-3 | 64 |
| V71A | 4.0e-3 | 3.9e-3 | N.D. | 240 |
| R86D* | 7.3e-3 | 7.6e-3 | N.D. | 112 |
| R86E* | N.D. | 3.7e-3 | 28e-3 | N.D. |

<sup>1</sup>Mutants at the CcdA-interacting and non-interacting residues are shown in green and orange colour respectively.
\*Rejuvenation with 10 nM CcdA<sub>45-72</sub>.
N.B.= No Binding, N.D.= Not Determined

**Table S3.** Related to Figure 4 and 5. The apparent dissociation rate constants (k_d_) for CcdB mutants at CcdA interacting residues obtained at each CcdA_45-72_ concentration have been normalised with respect to the corresponding WT dissociation rate constants obtained at the same concentrations. The fold change (k_d_ (Mutant)/k_d_(WT)) with 10, 25, 100 and 200 nM of CcdA_45-72_ peptide measured by SPR^1^ are shown. All the studies are carried out in 1XPBS, pH7.4 at 25 °C. Mutants with significantly slower rejuvenation than WT are highlighted in blue and mutants with altered binding to GyrA14 are marked with an asterisk (*).

| Mutants | $k_d(\text{mutant})/k_d(\text{WT})$ | | | |
| --- | --- | --- | --- | --- |
|  | 10 nM<br>CcdA<sub>45-72</sub> | 25 nM<br>CcdA<sub>45-72</sub> | 100 nM<br>CcdA<sub>45-72</sub> | 200 nM<br>CcdA<sub>45-72</sub> |
| WT<sup>a</sup> | 1.00 | 1.00 | 1.00 | 1.00 |
| Y8D | 0.01 | 0.04 | 0.03 | N.D. |
| Y8L | 0.15 | 0.17 | 0.54 | 0.58 |
| Y8V | 0.71 | 0.55 | 0.18 | 0.71 |
| R10G | 0.37 | 0.45 | 0.27 | 0.50 |
| R10K | 1.42 | 2.39 | 0.72 | 0.86 |
| S12G<sup>a</sup> | N.D. | N.D. | 0.17 | 0.22 |
| R13F | 0.13 | 0.77 | 0.21 | 0.32 |
| R13V | 0.18 | 0.52 | 0.48 | 0.54 |
| Y14A | N.D. | 0.21 | 0.08 | 0.13 |
| Y14W | 0.84 | 1.35 | 0.40 | 0.51 |
| P28N* | 0.90 | 0.95 | 0.53 | 0.71 |
| P28S* | 1.03 | 1.14 | 0.78 | 0.76 |
| P28Y* | 0.47 | 0.58 | 0.24 | 0.38 |
| V33M | 0.19 | 0.61 | 0.14 | 0.26 |
| A37L | 0.53 | 0.45 | 0.17 | 0.15 |
| L41E | 0.23 | 0.78 | 0.32 | 0.43 |
| L41M | 0.69 | 0.63 | 0.19 | 0.30 |
| L42E | 0.97 | 1.99 | 0.54 | 0.49 |
| L42K | 0.27 | 0.40 | 0.16 | 0.29 |
| L42W | 0.13 | 0.32 | 0.12 | 0.15 |
| S43T | 0.59 | 0.40 | 0.08 | 0.08 |
| S43Y | 0.19 | 0.21 | 0.07 | 0.07 |
| K45Y | 0.21 | 0.21 | 0.05 | 0.05 |
| V46L | 0.58 | 0.49 | 0.11 | 0.10 |
| V46R | 0.18 | 0.16 | 0.05 | 0.06 |
| V46W | 0.76 | 0.53 | 0.11 | 0.09 |
| V46W-Y51H | 0.11 | 0.12 | 0.15 | 0.15 |
| V46Y | 0.32 | 0.24 | 0.09 | 0.13 |
| **S47M** | 0.39 | 0.46 | 0.11 | 0.12 |
| **S47R** | 0.10 | 0.34 | 0.17 | 0.18 |
| **L50D\*** | 0.65 | 0.55 | 0.12 | 0.11 |
| **L50I\*** | 1.50 | 1.87 | 0.85 | 0.89 |
| **L50N** | 0.55 | 0.65 | 0.41 | 0.44 |
| **L50R** | 1.00 | 1.55 | 0.70 | 0.49 |
| **D67L** | 0.47 | 0.62 | 0.15 | 0.14 |
| **D67S** | 0.82 | 0.66 | 0.14 | 0.13 |
| **D67T\*** | 0.15 | 0.14 | 0.07 | 0.09 |
| **A69R** | 0.39 | 0.27 | 0.06 | 0.05 |
| **A69T** | 0.15 | 0.28 | 0.07 | 0.11 |
| **S70Q** | 0.50 | 0.39 | 0.10 | 0.09 |
| **V71A** | 0.50 | 0.35 | 0.08 | 0.08 |

N.D.= Not Determined
<sup>a</sup>Parameters from previous study (Aghera et al., 2020).

**Table S4.** Related to Figure 4 and 5. The apparent dissociation rate constants (k_d_) for CcdB mutants at GyrA-14 interacting sites, obtained at each CcdA_45-72_ concentration have been normalised with respect to the WT rejuvenation rate constants obtained at the same concentrations. The fold change (k_d_Mutant/k_d_WT) with 10, 25, 100 and 200 nM of CcdA_45-72_ peptide measured by SPR^1^ are shown. All the studies are carried out in 1XPBS, pH7.4 at 25 °C.

| **Mutants** | **$k_d$(mutant) /$k_d$(WT)** |  |  |  |
| --- | --- | --- | --- | --- |
|  | **10 nM**<br>**CcdA<sub>45-72</sub>** | **25 nM**<br>**CcdA<sub>45-72</sub>** | **100 nM**<br>**CcdA<sub>45-72</sub>** | **200 nM**<br>**CcdA<sub>45-72</sub>** |
| **WT<sup>a</sup>** | 1.00 | 1.00 | 1.00 | 1.00 |
| **I24L** | 1.94 | 2.36 | 1.04 | 1.18 |
| **I24K** | 2.10 | 2.61 | 1.00 | 1.00 |
| **I24Y** | 2.85 | 2.66 | 1.04 | 1.00 |
| **I25A** | 2.11 | 2.35 | 1.00 | 1.00 |
| **I25N** | 3.77 | 2.67 | 1.00 | 1.11 |
| **D26A** | 1.45 | 2.12 | 1.02 | 1.02 |
| **D26S** | 2.21 | 2.67 | 1.00 | 1.03 |
| **D26V** | 2.15 | 2.02 | 1.02 | 1.04 |
| **I101N** | 1.65 | 2.33 | 1.02 | 1.05 |

<sup>a</sup>Parameters from previous study (Aghera et al., 2020).

**Table S5.** Related to Figure 4 and 5. The apparent dissociation rate constants (k_d_) for CcdB mutants, at residues that neither interact with CcdA nor with GyrA14 obtained at each CcdA_45-72_ concentration, have been normalised with respect to the WT rejuvenation rate constants obtained at the same concentrations. The fold change (k_d_Mutant/k_d_WT) with 10, 25, 100 and 200 nM of CcdA_45-72_ peptide measured by SPR^1^ are shown. Mutants with significantly slower and faster rejuvenation than WT are highlighted in blue and red boxes respectively and mutants with altered binding to GyrA14 are marked with an asterisk (*). All the studies are carried out in 1XPBS, pH7.4 at 25 °C.

| Mutants | $k_d(\text{mutant}) / k_d(\text{WT})$ | | | |
| --- | --- | --- | --- | --- |
|  | 10 nM<br>CcdA<sub>45-72</sub> | 25 nM<br>CcdA<sub>45-72</sub> | 100 nM<br>CcdA<sub>45-72</sub> | 200 nM<br>CcdA<sub>45-72</sub> |
| WT<sup>a</sup> | 1.00 | 1.00 | 1.00 | 1.00 |
| K9R* | 2.95 | 2.64 | 1.03 | 0.96 |
| E11P | 1.03 | 1.11 | 1.02 | 0.96 |
| S38N* | 0.94 | 1.57 | 1.00 | 0.93 |
| A39R* | 0.73 | 0.84 | 0.20 | 0.24 |
| R40E | 1.44 | 1.65 | 0.96 | 0.96 |
| D44H | 1.03 | 1.83 | 1.00 | 0.95 |
| D44V | 1.00 | 1.05 | 0.96 | 0.91 |
| R48G | 1.05 | 0.96 | 0.98 | 0.99 |
| R48T* | 0.97 | 0.95 | 0.96 | 0.95 |
| V53P* | 0.68 | 0.68 | 0.24 | 0.26 |
| V54G* | 0.65 | 0.73 | 0.52 | 0.50 |
| H55C | 1.03 | 1.65 | 0.93 | 1.02 |
| H55R* | 1.00 | 1.36 | 0.93 | 1.04 |
| G57C* | 0.66 | 0.80 | 0.62 | 0.57 |
| D58N* | 2.68 | 2.85 | 0.96 | 0.94 |
| E59P* | 0.48 | 0.39 | 0.21 | 0.23 |
| E59Q* | 0.50 | 0.65 | 0.22 | 0.31 |
| S60E* | 0.58 | 0.43 | 0.10 | 0.12 |
| S60G | 1.35 | 1.83 | 1.01 | 1.07 |
| P72L | 1.08 | 0.90 | 0.96 | 0.95 |
| E79I | 0.47 | 0.40 | 0.23 | 0.52 |
| D82I | 0.97 | 1.06 | 1.03 | 1.03 |
| S84G* | 0.95 | 1.95 | 0.89 | 1.05 |
| R86D* | 1.23 | 2.10 | 0.89 | 1.04 |
| R86E* | 0.60 | 0.68 | 0.41 | 0.54 |

<sup>a</sup>Parameters from previous study (Aghera et al., 2020).

**Table S6.**
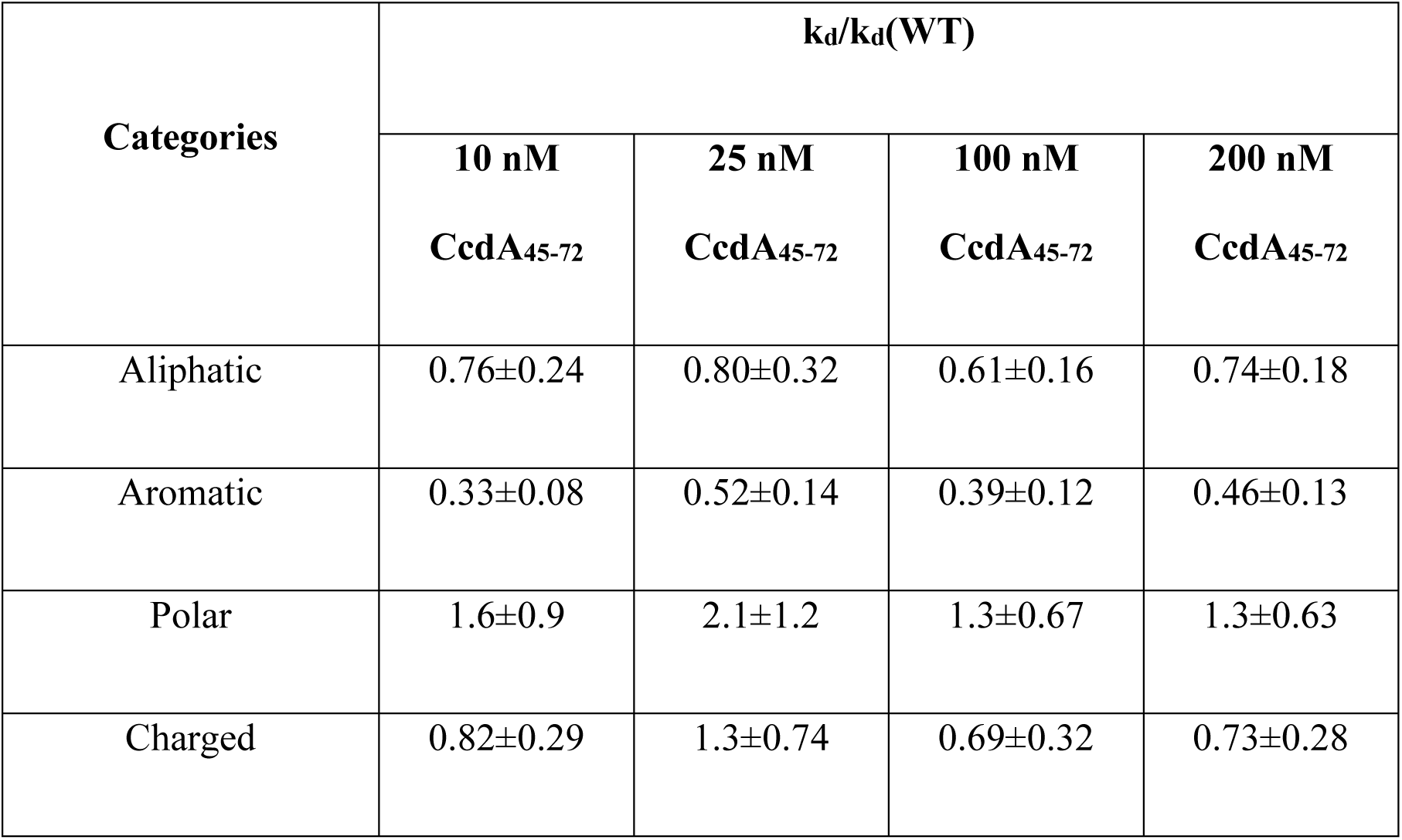
Related to Figure 4 and 5. Fold change in k_d_ resulting from amino acid substitutions (Mean±SEM) at positions used in this study. The rejuvenation rate constant for the CcdB mutants (k_d_) mediated by 10, 25, 100 and 200 nM of the CcdA_45-72_ peptide were normalized with the respective rejuvenation rate constant of WT CcdB (k_d_(WT)) obtained at each CcdA_45-72_ peptide concentration.

**Table S7.** Related to Figure 4 and 5. Kinetic parameters for binding to GyrA14 for CcdB mutants at CcdA-interacting sites measured by SPR^1^. Mutants with altered binding to GyrA14 are marked with a star (*). All studies are carried out in 1XPBS, pH7.4 at 25 °C.

| Mutants | $k_a$ (M<sup>-1</sup>s<sup>-1</sup>) | $k_d$ (s<sup>-1</sup>) | $K_D$ (nM) |
| --- | --- | --- | --- |
| WT<sup>a</sup> | 13±0.3×10<sup>5</sup> | 11.0±3.0×10<sup>-4</sup> | 1.4±0.6 |
| Y8D | 3.9±0.6×10<sup>5</sup> | 5.9±1.1×10<sup>-4</sup> | 5.5±3.2 |
| Y8L | 1.1±0.3×10<sup>5</sup> | 5.9±3.5×10<sup>-4</sup> | 6.9±0.5 |
| Y8V | 1.8±0.1×10<sup>5</sup> | 13.0±5.0×10<sup>-4</sup> | 7.1±0.3 |
| R10G | 1.8±0.2×10<sup>5</sup> | 19.0±6.0×10<sup>-4</sup> | 10.5±2.2 |
| R10K | 2.8±0.5×10<sup>5</sup> | 2.3±0.2×10<sup>-4</sup> | 0.9±0.01 |
| S12G | 12±0.4×10<sup>5</sup> | 17±0.1×10<sup>-4</sup> | 2.6±1.4 |
| R13F | 3.3±1.2×10<sup>5</sup> | 4.7±0.7×10<sup>-4</sup> | 1.2±0.3 |
| R13V | 4.8±0.7×10<sup>5</sup> | 6.6±2.0×10<sup>-4</sup> | 1.3±0.3 |
| Y14A | 14.0±4.0×10<sup>5</sup> | 18.0±2.0×10<sup>-4</sup> | 2.1±0.8 |
| Y14W | 4.7±0.7×10<sup>5</sup> | 5.0±1.5×10<sup>-4</sup> | 1.1±0.3 |
| P28N* | 0.7±0.2×10<sup>5</sup> | 21.9±2.0×10<sup>-4</sup> | 29.4±1.4 |
| P28S* | 2.2±0.2×10<sup>5</sup> | 33.5±1.0×10<sup>-4</sup> | 14.6±0.3 |
| P28Y* | 0.5±0.1×10<sup>5</sup> | 7.5±0.2×10<sup>-4</sup> | 14.5±1.3 |
| V33M | 79.9±13.0×10<sup>5</sup> | 5.3±0.3×10<sup>-4</sup> | 0.7±0.02 |
| A37L | 27.0±6.0×10<sup>5</sup> | 31.6±1.0×10<sup>-4</sup> | 1.4±0.4 |
| L41E | 3.8±1.7×10<sup>5</sup> | 7.7±0.4×10<sup>-4</sup> | 1.5±0.3 |
| L41M | 5.4±0.4×10<sup>5</sup> | 5.5±0.8×10<sup>-4</sup> | 1.0±0.2 |
| L42E | 6.2±1.4×10<sup>5</sup> | 4.2±0.6×10<sup>-4</sup> | 0.8±0.2 |
| L42K | 1.6 ±1.2×10<sup>5</sup> | 5.2 ±0.1×10<sup>-4</sup> | 3.4 ±0.9 |
| L42W | 5.0±2.0×10<sup>5</sup> | 6.9±1.1×10<sup>-4</sup> | 6.3±1.2 |
| S43T | 3.4±0.7×10<sup>5</sup> | 5.1±1.3×10<sup>-4</sup> | 7.0±0.4 |
| S43Y | 14.0±6.0×10<sup>5</sup> | 11.0±1.0×10<sup>-4</sup> | 1.3±0.04 |
| K45Y | 3.8±0.4×10<sup>5</sup> | 42.0±2.0×10<sup>-4</sup> | 10.6±4.0 |
| V46L | 5.1±1.8×10<sup>5</sup> | 15.0±2.0×10<sup>-4</sup> | 3.05±0.7 |
| V46R | 3.6±0.2×10<sup>5</sup> | 39.0±1.0×10<sup>-4</sup> | 5.8±0.01 |
| V46W | 3.8±2.0×10<sup>5</sup> | 14.0±3.0×10<sup>-4</sup> | 3.4±0.1 |
| V46W-Y51H<sup>a</sup> | 2.4±0.5×10<sup>5</sup> | 11.2±2.0×10<sup>-4</sup> | 2.2±0.1 |
| V46Y | 4.4±1.5×10<sup>5</sup> | 5.9±0.8×10<sup>-4</sup> | 2.0±0.7 |
| S47M | 2.9±0.7×10<sup>5</sup> | 26.6±2.0×10<sup>-4</sup> | 9.2±0.1 |
| S47R | 5.6±0.8×10<sup>5</sup> | 3.7±0.9×10<sup>-4</sup> | 0.7±0.2 |
| L50D* | 29.0±18.0×10<sup>5</sup> | 25.2±6.0×10<sup>-4</sup> | 2.1±0.1 |
| L50I* | 62.0±7.0×10<sup>5</sup> | 11.8±3.0×10<sup>-4</sup> | 1.9±0.4 |
| L50N | 2.5±0.8×10<sup>5</sup> | 2.5±0.3×10<sup>-4</sup> | 1.2±0.3 |
| L50R | 4.7±1.9×10<sup>5</sup> | 3.5±0.2×10<sup>-4</sup> | 1.5±0.3 |
| D67L | 2.0±1.5×10<sup>5</sup> | 9.1±1.2×10<sup>-4</sup> | 4.8±0.9 |
| D67S<sup>a</sup> | 5.5±1.5×10<sup>5</sup> | 3.1±0.9×10<sup>-4</sup> | 0.9±0.7 |
| D67T* | 34.0±12.0×10<sup>5</sup> | 9.4±1.2×10<sup>-4</sup> | 2.2±0.7 |
| A69R<sup>a</sup> | 7.6±1.4×10<sup>5</sup> | 24.4±1.0×10<sup>-4</sup> | 3.5±0.5 |
| A69T | 6.3±0.5×10<sup>5</sup> | 5.9±0.7×10<sup>-4</sup> | 0.9±0.02 |
| S70Q | 7.7±3.6×10<sup>5</sup> | 11.8±3.0×10<sup>-4</sup> | 1.5±0.4 |
| V71A | 9.6±0.4×10<sup>5</sup> | 32.7±3.0×10<sup>-4</sup> | 3.4±0.2 |

<sup>1</sup> The SPR experiments have been performed once with four different concentrations of the protein (in each experiment) and the listed error is the standard error derived from the values at multiple concentrations. <sup>a</sup>Parameters from previous study (Ahmed et al., 2022; Chattopadhyay et al., 2021).

**Table S8.** Related to Figure 4 and 5. Kinetic parameters for binding to GyrA14 for CcdB mutants at residues 24-26 and 101 involved in GyrA14 interaction, measured by SPR^1^. All studies are carried out in 1XPBS, pH7.4 at 25 °C.

| **Mutants** | **k<sub>a</sub> (M<sup>-1</sup>s<sup>-1</sup>)** | **k<sub>d</sub> (s<sup>-1</sup>)** | **K<sub>D</sub> (nM)** |
| --- | --- | --- | --- |
| **WT<sup>a</sup>** | 13±0.3×10<sup>5</sup> | 11.0±3.0×10<sup>-4</sup> | 1.4±0.6 |
| **I24E** | N.B. | N.B. | N.B. |
| **I24L** | 0.4±0.08×10<sup>5</sup> | 9.0±4.0×10<sup>-4</sup> | 27.2±6.0 |
| **I24K** | 0.2±0.06×10<sup>5</sup> | 30.5±7.0×10<sup>-4</sup> | 170.0±7.0 |
| **I24Y** | 1.8±2.2×10<sup>5</sup> | 64.5±7.0×10<sup>-4</sup> | 30.3±1.0 |
| **I25A** | 0.6±0.09×10<sup>5</sup> | 8.8±0.4×10<sup>-4</sup> | 15.2±2.4 |
| **I25N** | 0.7±0.2×10<sup>5</sup> | 26.6±7.0×10<sup>-4</sup> | 38.0±4.9 |
| **D26A** | 1.4±0.4×10<sup>5</sup> | 26.7±3.0×10<sup>-4</sup> | 24.6±7.0 |
| **D26S** | 0.1±0.2×10<sup>5</sup> | 73.5±2.0×10<sup>-4</sup> | 55.2±9.1 |
| **D26V** | 0.5±0.07×10<sup>5</sup> | 10.8±2.0×10<sup>-4</sup> | 23.0±2.3 |
| **I101N** | 0.9±0.2×10<sup>5</sup> | 27.3±8.0×10<sup>-4</sup> | 30.2±0.3 |

<sup>1</sup> The SPR experiments have been performed once with four different concentrations of the protein (in each experiment) and the listed error is the standard error derived from the values at multiple concentrations.
N.B.=No Binding
<sup>a</sup>Parameters from previous study (Chattopadhyay et al., 2021).

**Table S9.** Related to Figure 4 and 5. Kinetic parameters for binding to GyrA14 for CcdB mutants, at residues that neither interact with CcdA nor with GyrA14, measured by SPR^1^. Mutants with altered binding to GyrA14 are marked with a star (*). All studies are carried out in 1XPBS, pH7.4 at 25 °C.

| **Mutants** | **k<sub>a</sub> (M<sup>-1</sup>s<sup>-1</sup>)** | **k<sub>d</sub> (s<sup>-1</sup>)** | **K<sub>D</sub> (nM)** |
| --- | --- | --- | --- |
| **WT<sup>a</sup>** | 13±0.3×10<sup>5</sup> | 11.0±3.0×10<sup>-4</sup> | 1.4±0.6 |
| **K9R\*** | 0.9±0.1×10<sup>5</sup> | 8.1±1.2×10<sup>-4</sup> | 9.4±1.9 |
| **E11P<sup>a</sup>** | 1.3±0.3×10<sup>5</sup> | 4.5±0.6×10<sup>-4</sup> | 3.7±0.4 |
| **S38N\*** | 19.0±6.0×10<sup>5</sup> | 35.3±1.0×10<sup>-4</sup> | 1.9±0.2 |
| **A39R** | 29.0±7.0×10<sup>5</sup> | 81.9±2.0×10<sup>-4</sup> | 1.2±0.1 |
| **R40E** | 5.5±1.7×10<sup>5</sup> | 22.0±4.0×10<sup>-4</sup> | 4.9±1.1 |
| **D44H** | 9.4±3.2×10<sup>5</sup> | 1.2±0.9×10<sup>-4</sup> | 0.2±0.04 |
| **D44V<sup>a</sup>** | 2.9±0.7×10<sup>5</sup> | 4.8±1.9×10<sup>-4</sup> | 1.6±0.06 |
| **R48G** | 6.6±1.1×10<sup>5</sup> | 9.3±0.3×10<sup>-4</sup> | 1.7±0.7 |
| **R48T\*** | 3.1±0.5×10<sup>5</sup> | 7.0±1.6×10<sup>-4</sup> | 2.2±0.3 |
| **V53P\*** | 33.0±16.0×10<sup>5</sup> | 76.6±21.0×10<sup>-4</sup> | 4.1±0.2 |
| **V54G\*** | 18.0±3.0×10<sup>5</sup> | 32.7±5.0×10<sup>-4</sup> | 1.9±0.4 |
| **H55C** | 3.2±1.5×10<sup>5</sup> | 19.0±4.0×10<sup>-4</sup> | 1.4±0.2 |
| **H55R\*** | 26.0±11.0×10<sup>5</sup> | 31.0±5.0×10<sup>-4</sup> | 1.7±0.4 |
| **G57C\*** | 43.0±14.0×10<sup>5</sup> | 61.4±4.0×10<sup>-4</sup> | 3.0±0.2 |
| **D58N<sup>a</sup>** | 2.7±0.3×10<sup>5</sup> | 2.2±0.1×10<sup>-4</sup> | 1.5±0.8 |
| **E59P\*** | 22.0±6.0×10<sup>5</sup> | 37.9±3.0×10<sup>-4</sup> | 2.1±0.04 |
| **E59Q\*** | 30.0±10.0×10<sup>5</sup> | 33.4±5.0×10<sup>-4</sup> | 2.6±0.7 |
| **S60E\*** | 46.0±21.0×10<sup>5</sup> | 21.0±4.0×10<sup>-4</sup> | 2.4±0.2 |
| **S60G** | 8.9±0.4×10<sup>5</sup> | 3.5±0.6×10<sup>-4</sup> | 1.3±0.2 |
| **P72L** | 4.7±1.4×10<sup>5</sup> | 21.5±3.0×10<sup>-4</sup> | 4.6±0.1 |
| **E79I** | 27.0±7.0×10<sup>5</sup> | 24.9±5.0×10<sup>-4</sup> | 1.1±0.3 |
| **D82I** | 21.0±8.0×10<sup>5</sup> | 31.9±1.0×10<sup>-4</sup> | 2.1±0.7 |
| **S84G\*** | 31.0±12.0×10<sup>5</sup> | 58.1±8.0×10<sup>-4</sup> | 3.8±1.9 |
| **R86D<sup>a</sup>** | 4.2±0.5×10<sup>5</sup> | 3.9±0.5×10<sup>-4</sup> | 0.9±0.2 |
| **R86E\*** | 8.3±2.8×10<sup>5</sup> | 47.9±13.7×10<sup>-4</sup> | 6.1±0.5 |

<sup>1</sup> The SPR experiments have been performed once with four different concentrations of the protein (in each experiment) and the listed error is the standard error derived from the values at multiple concentrations. <sup>a</sup>Parameters from previous study (Ahmed et al., 2022; Chattopadhyay et al., 2021).

**Table S10.**
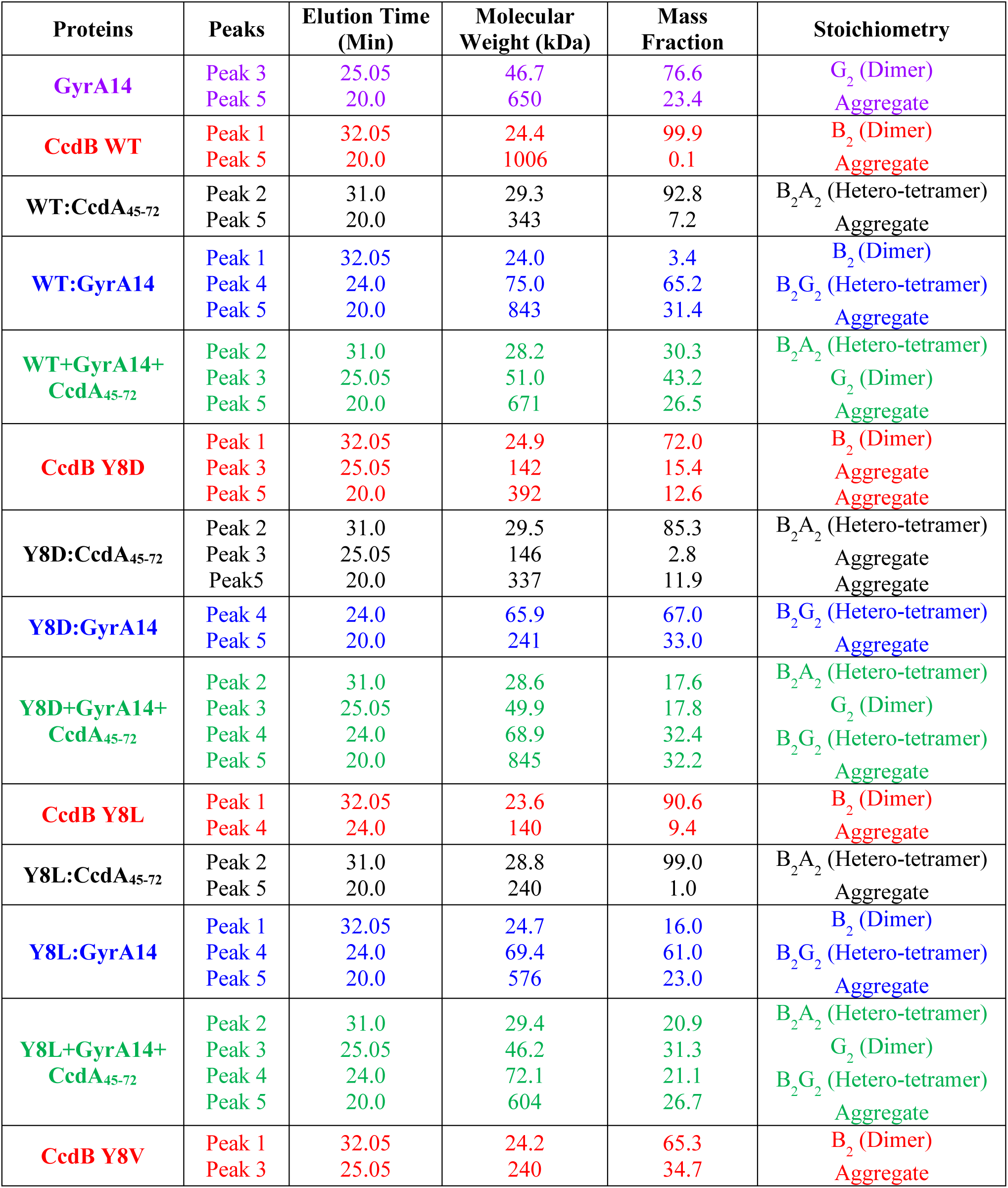
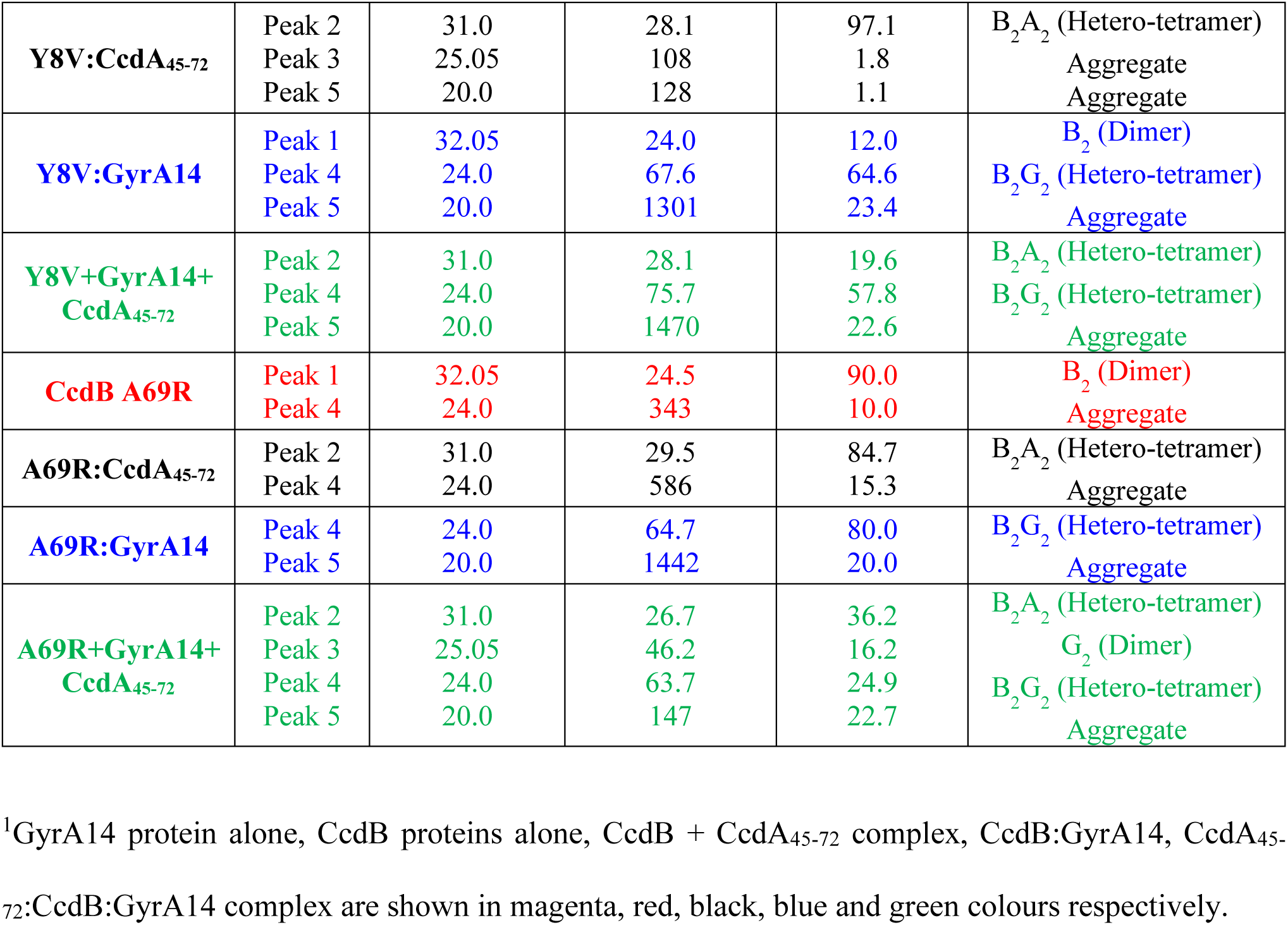
Related to Figure S15. The mass fractions and corresponding molecular weights of each of the peaks (Figure S15) observed for the proteins in isolation and when complexed, obtained from SEC-MALS and represented as (B:CcdB, A:CcdA, G:GyrA14)^1^.

